# Gene flow from the European wild apple and selection shaped the domesticated apple (*Malus domestica* Borkh.) genome

**DOI:** 10.1101/2025.09.18.676739

**Authors:** Xilong Chen, Ronan Dadole, Komlan Avia, Anthony Venon, Mathieu Brisson, Carine Remoué, Dong Zhang, Ivan Gabrielyan, Anush Nersesyan, Anamaria Roman, Tudor Ursu, Ammar Alhmedi, Dany Bylemans, Tim Beliën, Agnès Rousselet, Martine Le Guilloux, Enrique Dapena, Charles-Eric Durel, Thomas Kirisits, Gayle Volk, Frédérique Didelot, Arnaud Lemarquand, Thierry Hance, Amandine Cornille

## Abstract

How selection and demography shape genomes of long-lived crops remains largely unresolved. Using apple (*Malus domestica*) as a model, we integrate 218 whole genomes (68 cultivated dessert/cider; 150 wild: *M. sieversii, M. orientalis, M. sylvestris*), RNA-seq, and a flowering-time GWAS to resolve how these forces forged the cultivated apple genome. Despite weak neutral differentiation and widespread admixture, dessert and cider apples form distinct gene pools that derive primarily from *M. sieversii–M. orientalis* rather than European *M. sylvestris*. We find no evidence of a strong domestication bottleneck, as expected in perennials. Demography-aware selection scans reveal largely non-overlapping targets: dessert shows more hard sweeps at genes linked to fruit quality, disease resistance, and flowering, whereas cider shows proportionally more soft sweeps and balancing selection; RNA-seq differential expression supports these candidates. Wild-to-crop introgression from *M. sylvestris* is extensive but heterogeneous by context: some introgressed tracts concentrate in hard-sweep regions and approach fixation (consistent with rapid, targeted uptake), whereas others persist at intermediate frequencies with soft-sweep signatures (consistent with diffuse, recurrent introgression of adaptive alleles). Extending to the phenotype, the lead chromosome 9 flowering-time association lies within an introgressed segment near a transposable element and is separated from sweep peaks, consistent with regulatory/polygenic control. Cultivated apples carry a lower predicted deleterious load than wild relatives. Together, these results provide one of the most comprehensive population genomic portraits of a perennial fruit tree domestication, clarifying how selection and adaptive introgression jointly shaped the cultivated apple genome architecture and yielding actionable targets for breeding and conservation.

**Significance statement:** Perennial crops are underexplored compared to annuals, leaving open the question of how selection, gene flow, and demography shape their genomes. Using the apple tree (*Malus domestica*), we analyzed 218 genomes, along with expression and trait data. Despite weak genome-wide differences, dessert and cider apples form distinct gene pools. Widespread gene flow from the European wild apple supplied adaptive alleles, with contrasting dynamics: in cider, a few introgressed DNA segments rose rapidly; in dessert, many variants shifted gradually. Cultivated apples also carry a lower predicted burden of harmful mutations than wild relatives. Together, these results redefine perennial domestication and pinpoint genomic targets to accelerate breeding and conservation.

## Introduction

Domestication provides an experimental setting in evolution, revealing how selection and demography (including gene flow) shape genomes over a relatively short evolutionary timescale. Since Darwin first recognized domestication as a cornerstone model for understanding the origins of biodiversity (1), it has become clear that plant domestication involves both artificial selection on favorable traits and complex demographic events, including population bottlenecks and gene flow (2). Perennial fruit trees, however, differ fundamentally from annual crops in their domestication dynamics. Their long generation times, high outcrossing rates, and frequent reliance on clonal propagation lead to milder domestication bottlenecks, retention of gene flow with wild relatives, and fewer generations since domestication onset, resulting in unique genomic outcomes (3–5).

Despite growing genomic insights into perennials, significant knowledge gaps remain. The extent and dynamics of wild-to-crop gene flow, the nature of selection (positive vs. balancing (4)) acting on domesticated genomes, and the accumulation of deleterious mutations—the so-called “cost of domestication”—are still poorly understood in perennial fruit trees (3). While annuals often exhibit elevated genetic load post-domestication, the impact on long-lived, clonally propagated crops is less clear. For example, studies in grapevine suggest that somatic mutations may contribute significantly, but broader patterns across perennials remain elusive (6, 7).

The cultivated apple (*Malus domestica* Borkh.) is a powerful model for studying the domestication of perennial plants. Its extraordinary phenotypic diversity—ranging in fruit color, texture, and flavor—has led to its classification into dessert and cider cultivars (8, 9), with sometimes overlapping phenotypic characteristics (10). Genomic analyses reveal a continuum between dessert and cider apples, with two primary gene pools that are admixed yet genetically structured (11, 12). Previous studies have traced the origin of cultivated apples primarily to *M. sieversii* in Central Asia, with additional contributions from *M. orientalis* in the Caucasus and the European crabapple, *M. sylvestris*, through wild-to-crop introgression (13). Thus, the domestication history of cider and dessert apples, as well as wild-to-crop gene flow, is relatively well understood (11, 14). However, the genomic architecture and evolutionary consequences of this admixture remain poorly resolved, leaving open key questions about the relative roles of selection, introgression, and demographic history in shaping apple domestication (14–17).

Here, we aim to disentangle the evolutionary processes that shaped the cultivated apple genome, asking how selection, introgression, and demographic history jointly influenced perennial domestication. To address these questions, we analyzed a genome-wide dataset of 218 apple accessions (including 140 newly sequenced), including dessert and cider cultivars as well as their closest wild relatives (*Malus sieversii, M. orientalis*, and *M. sylvestris*). Unlike previous studies that focused primarily on Central Asian *M. sieversii*, our work is distinguished by the inclusion of an unprecedentedly comprehensive collection of European *M. sylvestris* sampled across its full geographic range. This is crucial because *M. sylvestris* has been a major contributor to recent wild-to-crop introgressions, making this dataset ideal for evaluating the adaptive role of introgression in the domestication of this perennial species. Specifically, we sought to (i) characterize the population structure and origin of apple genetic diversity, (ii) identify genomic signatures of selection—both positive and balancing—associated with domestication, (iii) investigate the extent, sources, and adaptive roles of wild-to-crop introgression, and (iv) evaluate the patterns and consequences of deleterious mutation load. To anchor these inferences at the trait and expression levels, we tested flowering-time differentiation via GWAS and profiled expression of selection candidates with RNA-seq, linking population genomic signals to phenotypic and transcriptional divergence. By integrating population genomics, selection scans, ancestry inference, gene expression analyses, and GWAS, this study provides a comprehensive view of the evolutionary forces—wild introgression, selection, and demography—that forged the cultivated apple genome, offering new insights into the evolutionary dynamics of perennial crop domestication.

## Results

### Cider and dessert apples formed distinct gene pools, admixed with wild relatives

We re-sequenced or retrieved whole-genome short-read data from 218 apple accessions, including 45 dessert and 23 cider cultivars, as well as 150 wild individuals: 87 *M. sylvestris* (Europe), 40 *M. sieversii* (Kazakhstan and China), 10 *M. orientalis* (Caucasus), and 13 *M. baccata* (China). After quality control (removing clones and individuals with SNP missing rate > 30%), 201 accessions were retained, in which we identified 28,377,551 SNPs (SI Appendix, Fig. S1, Dataset S1, Tables S1–S4) (18, 19).

Population structure and admixture analyses based on 31,300 unlinked synonymous SNPs revealed six major genetic clusters: dessert apples (orange), cider apples (pink), and four wild groups corresponding to their geographic origins (Fig. 1A, B; SI Appendix, Figs. S2, S3; Tables S5, S6). *M. sieversii* and *M. orientalis* were not separated at any K value (Fig. S2). Neighbor-net and principal component analyses confirmed that *M. baccata* was highly differentiated from the other species (Fig. 1C, D) and therefore used as the outgroup. Western and Eastern European *M. sylvestris* were genetically distinct from *M. sieversii-M. orientalis*. Both cider and dessert apples were closer to *M. sieversii*–*M. orientalis*, rather than *M. sylvestris, as* in a previous study (20).

**Figure 1.**
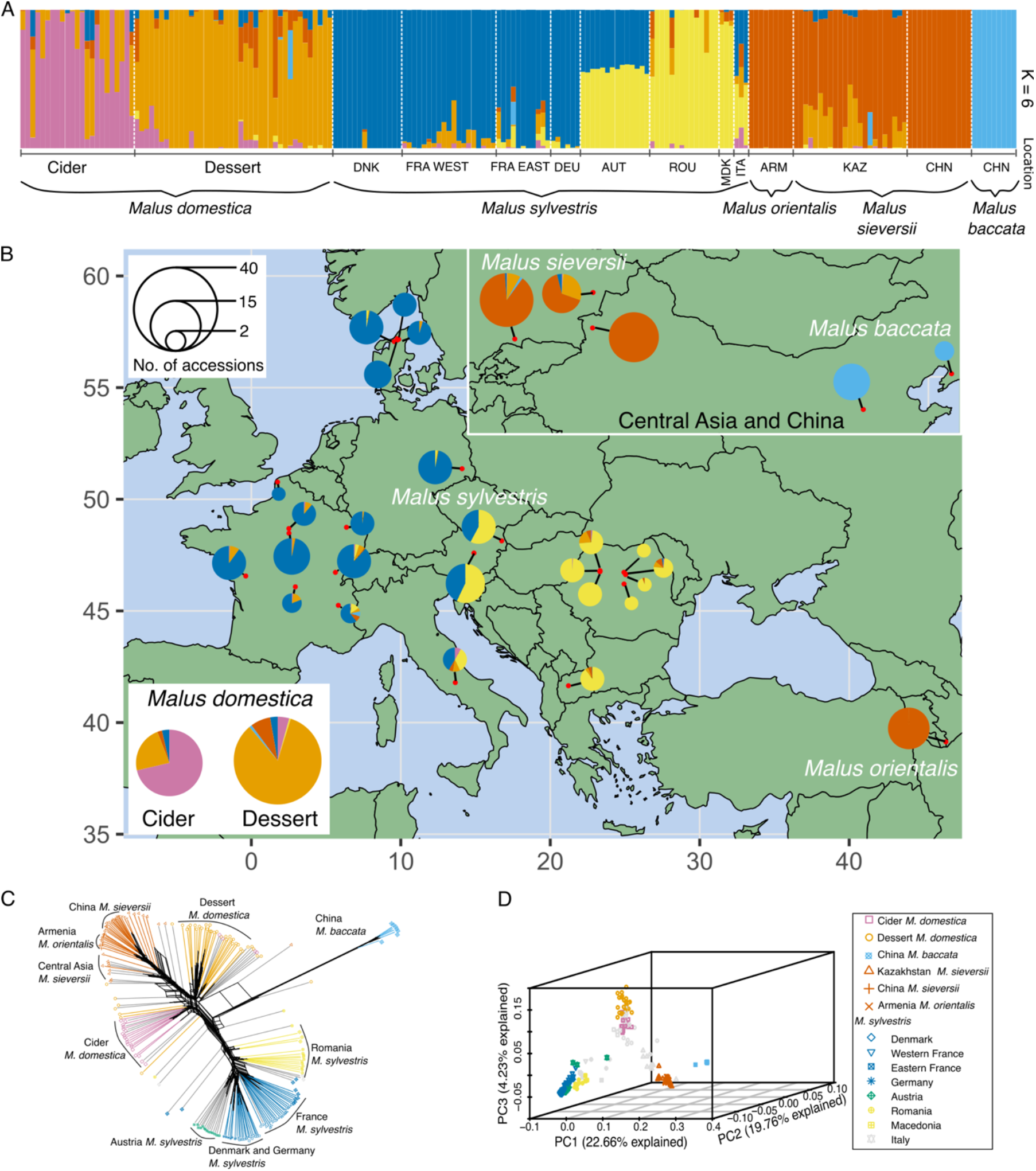
Population genetic structure and variation between cultivated and wild apples in Eurasia. (**A**) Population structure of cultivated and wild apple accessions inferred by fastSTRUCTURE with *K* = 6 (*N* = 201 accessions). Each individual is represented by a vertical bar, partitioned into *K* segments corresponding to the proportion of genetic ancestry from the *K* clusters. Wild accessions are sorted geographically from west to east in the plot, with the countries of origin listed: DNK (Denmark), FRA WEST (Western France), FRA EAST (Eastern France), DEU (Germany), AUT (Austria), ROU (Romania), MKD (Macedonia), ITA (Italy), ARM (Armenia), KAZ (Kazakhstan), and CHN (China). Distinct genetic clusters comprise cider apples (pink), dessert apples (orange), Western *M. sylvestris* (blue), Eastern *M. sylvestris* (yellow), *M. sieversii* and *M. orientalis* (red), and *M. baccata* (green). (**B**) Map showing the average proportions of fastSTRUCTURE membership coefficients at *K* = 6 for all wild individuals from each sampling location (see Dataset S1) and for cider or dessert varieties (lower left corner, outside the map, for cultivated apples). The size of each pie chart is proportional to the sample size. (**C**) Neighbor-net tree depicting relationships among wild and cultivated apple accessions and their associated populations, as identified by fastSTRUCTURE with *K* = 6. Colors correspond to the genetic groups inferred with *K* = 6, and admixed individuals (membership coefficient < 80% for any given gene pool) are shown in gray. (**D**) Principal component analysis (PCA) illustrating the genetic variation among wild and cultivated apple accessions. The total variance explained by each of the first three components is indicated along the respective axis. All of the analyses used the 31,300 unlinked and synonymous SNPs on 201 genotypes of apples.

FastSTRUCTURE membership coefficients (21) were used to assign genotypes to a given population (i.e., group of individuals including genotypes with membership coefficient > 0.8 to any cluster; SI Appendix, Fig. S4). Six populations were defined: DomC (cider apples), DomD (dessert apples), SiOr (*M. sieversii* + *M. orientalis*), SylW (Western *M. sylvestris*), SylE (Eastern *M. sylvestris*), and Bacc (*M. baccata*).

These results confirmed that cider and dessert apples represent genetically distinct but admixed gene pools, both of which are genetically closely related to Central Asian and Caucasian wild apples rather than European *M. sylvestris*.

### Divergence and demographic and histories of wild and cultivated apples

Genetic differentiation estimates confirmed that Bacc was the most divergent (SI Appendix, Fig. S5, Table S5). Using Bacc as an outgroup, the species-tree topology and coalescent-based ASTRAL phylogeny tree (SI Appendix, Fig. S6) showed SylW and SylE formed a clade, but could not resolve whether DomC was closer to SiOr or DomD. Repeated ASTRAL analyses using the same number of individuals per population consistently recovered two alternative species-tree topologies (SI Appendix, Fig. S7). SylE, SylW, SiOr, DomC, and DomD displayed weak genetic differentiation, further consistent with a recent divergence and/or gene flow (Figure S5, Table S5).

Cultivated apples (DomC and DomD) exhibited higher genetic diversity than the wild populations SiOr and SylW (SI Appendix, Table S5), indicating a lack of bottleneck during domestication. Demographic inference revealed a gradual effective population size (*Ne*) declined from the penultimate glaciation to the mid-Holocene (∼10,000 years ago), followed by divergent trends (Fig. 2A, B; SI Appendix, Fig. S8). SiOr and SylW expanded post-glacially, whereas SylE maintained a relatively stable *Ne*. Among cultivated apples, DomD showed a demographic expansion ∼2,500 years ago, whereas DomC showed a contraction (SI Appendix, Fig. S8).

**Figure 2.**
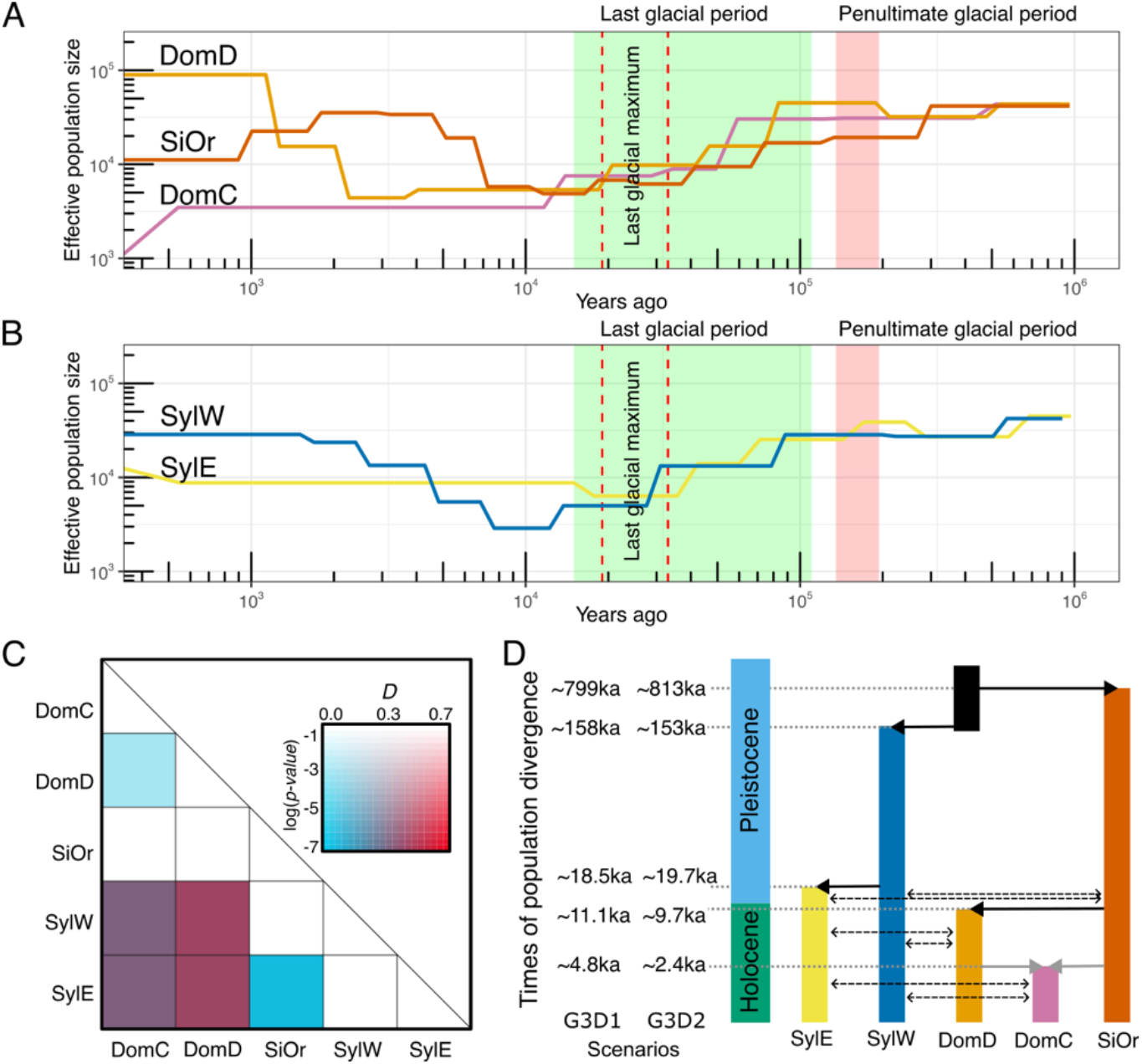
Demographic and divergence histories of wild and cultivated apple populations. (**A**) Changes in effective population size of cider apples (DomC), dessert apples (DomD), and *Malus sieversii* (SiOr) over time, as inferred by SMC++. (**B**) Changes in the effective population size of Western (SylW) and Eastern (SylE) European wild apples over time, as inferred by SMC++. Major climatic events are shown: the penultimate glacial period (red), the last glacial period (green), and the last glacial maximum (dashed vertical lines). (**C**) Patterson’s *D*-statistic (ABBA-BABA test) was estimated between five apple populations. DomC, cider apples; DomD, dessert apples; SiOr, *M. sieversii* and *M. orientalis*; SylE, Eastern European *M. sylvestris*; SylW, Western European *M. sylvestris*. Bacc (*M. baccata*) was used as the outgroup. (**D**) Schematic graph of the apple demographic history. Demographic parameters were estimated by fastsimcoal2. Solid lines with arrows represent divergence events, while dashed lines with arrows represent gene flow. The left side indicates the timing of divergence events (ka, thousands of years ago) for G3D1 and G3D2 scenarios (G3 includes gene flow between the ancestors of DomC/DomD and SiOr, gene flow between the ancestors of SylW/SylE and SiOr, and gene flow between DomC/DomD and SylW/SylE populations. Divergence scenarios include D1, where DomC and DomD both diverge from SiOr, with DomC diverging more recently and SylE diverging from SylW; D2, where DomD diverges from SiOr, followed by DomC diverging from DomD, and SylE diverging from SylW.).

Post-glacial climate likely shaped the demography of wild apples, while the expansion of DomD reflects its historical spread, and the contraction of DomC indicates contrasting domestication histories to DomD, that was taken into account for selection test below.

### Genome-wide introgression from European wild apples

Patterson’s *D*-statistics (22, 23) and *f*_4_-ratios (24) using Bacc as an outgroup revealed significant introgression from SylW into both DomD and DomC, with weaker signals from SylE. Minor gene flow was detected between SylE and SiOr, indicating elevated gene flow from an ancestor of *M. sylvestris* into the *M. domestica* gene pool (Fig. 2C; SI Appendix, Fig. S9; Table S6). The massive ‘proto-gene flow’ from ancestral *M. sylvestris* could also explain the observed *Ne* expansion above (16, 25).

Five-taxon *D*-statistics (*D*_FOIL_) (26) and topology weighting analyses (27) confirmed contributions from both Western and Eastern *M. sylvestris* to DomD and DomC (SI Appendix, Figs. S10–S12). The dominant species-tree topology indicated introgression from SylW into the common ancestor of DomC and DomD. This was further supported by *f*-branch statistics (28) and the best-fit qpGraph model (29), which inferred multiple admixture events, including one predating the DomC–DomD split (SI Appendix, Figs. S13, S14; Table S7). Among eight demographic models (SI Appendix, Fig. S15) tested with fastsimcoal2 (30), the best-fitting scenarios incorporated gene flow from SiOr into SylW/SylE and from SylW/SylE into DomC and DomD, with stronger introgression into DomD (Fig. 2D; SI Appendix, Fig. S16; Table S8).

Per-chromosome *D*-statistics and topology weighting revealed heterogeneous wild-to-crop introgression (SI Appendix, Figs. S17–S18). In DomC, chromosome 13 showed the highest SylE/SylW admixture, whereas in DomD, chromosomes 16 and 14 were most affected. Sliding-window *fd* scans (31) indicated highly introgressed regions enriched for genes linked to nitrogen metabolism, terpene biosynthesis, and steroid biosynthesis (SI Appendix, Figs. S19).

Wild-to-crop introgression was therefore extensive and heterogeneous, with DomD showing stronger signals. Introgressed regions overlap with genes associated with metabolism and adaptation, suggesting their potential functional role (SI Appendix, Figs. S20).

### Genomic regions under selection

Selection scans identified regions under positive selection (hard or soft sweeps) and balancing selection across all populations.

Most positive selection genes were population-specific; however, 27 were shared between DomC and DomD, including genes linked to fruit quality, disease resistance, and flowering time (Fig. 3A, B; SI Appendix, Dataset S2 and S3). We also identified 81 genes under balancing selection (Fig. S21; Dataset S4), 44 of which were shared among populations (SI Appendix, Fig. S22). These included stress-response genes such as *MdDREB2* (SI Appendix, Fig. S23) (32). Twenty-three balancing selection genes were specifically shared between DomC/DomD and *M. sylvestris* (SylE or SylW), including *MdVIP5* (MD04G1191400), a flowering-related gene (33). Cider apples (DomC) displayed more soft sweeps and balancing selection, whereas dessert apples (DomD) showed more hard sweeps (Table 1). DomC lacked the fruit size gene *fs15*.*1* (15, 34) but had unique balancing selection targets such as *RPW8-NBS* (SI Appendix, Fig. S24) (35). DomD contained two disease-resistance genes and eight flowering genes under hard sweeps. Expression analyses confirmed that 41 DomC and 54 DomD positively selected genes were differentially expressed compared to wild relatives (SI Appendix, Dataset S2).

**Table 1.** Size and gene count of regions under positive selection and balancing selection in the five cultivated and wild apple populations.

| Population | Positive selection |  |  |  |  |  | Balancing selection |  |
| --- | --- | --- | --- | --- | --- | --- | --- | --- |
|  | Hard sweep |  | Soft sweep |  | Total |  |  |  |
|  | Region size (Kb) | Number of genes | Region size (Kb) | Number of genes | Region size (Kb) | Number of genes | Region size (Kb) | Number of genes |
| DomC | 604.27 | 83 | 1,124.19 | 133 | 1,728.46 | 216 | 82.43 | 39 |
| DomD | 1,092.84 | 169 | 532.95 | 85 | 1,625.79 | 254 | 36.14 | 22 |
| SiOr | 276.31 | 21 | 74.71 | 11 | 351.02 | 32 | 21.11 | 14 |
| SylE | 335.73 | 26 | 17.64 | 3 | 353.37 | 29 | 62.56 | 36 |
| SylW | 644.75 | 62 | 123.93 | 12 | 768.68 | 74 | 139.04 | 75 |

Notes: DomC, Cider *M. domestica*; DomD, Dessert *M. domestica*; SiOr, *M. sieversii* and *M. orientalis*; SylE, Eastern European *M. sylvestris*; SylW, Western European *M. sylvestris*.

**Figure 3.**
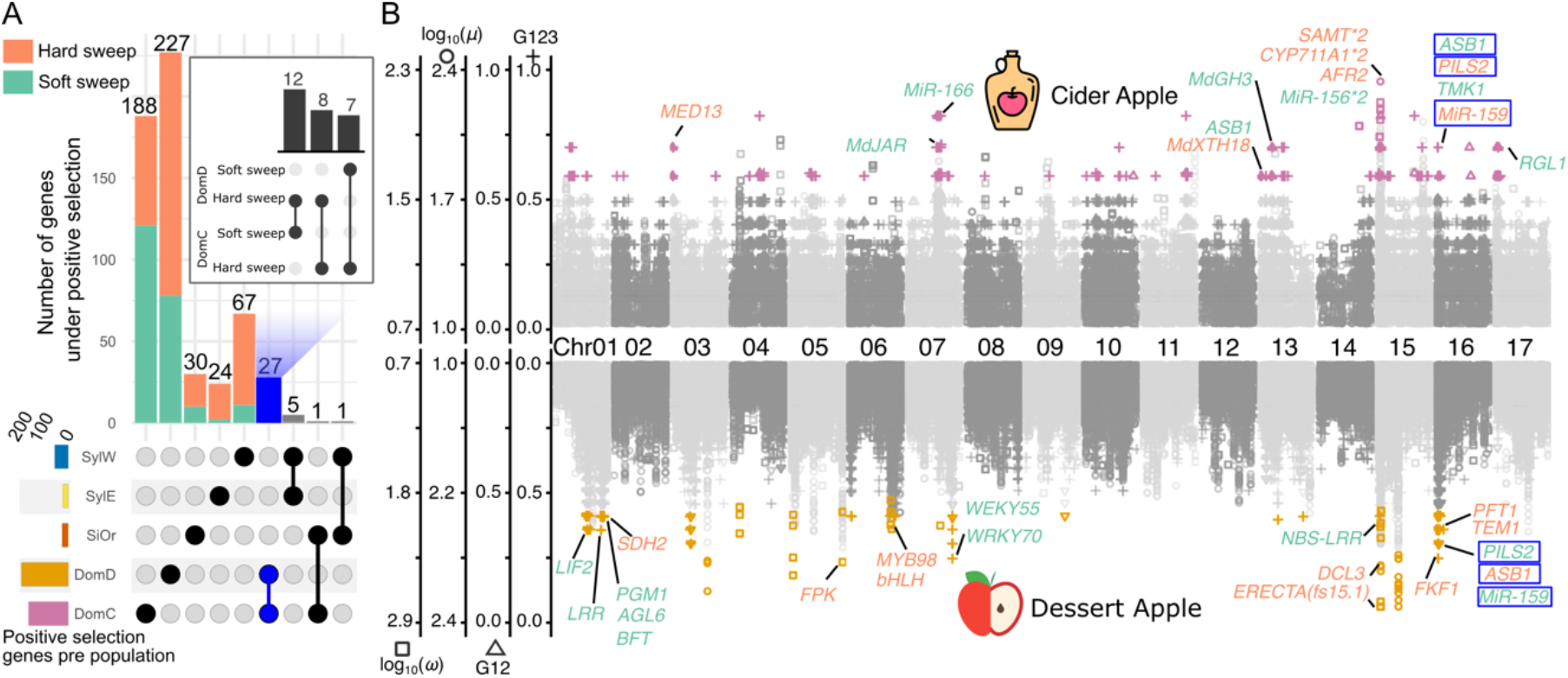
Positive selection along the genomes of cider and dessert cultivated apples. (**A**) UpSet plot of genes under positive selection between and among the five apple populations. The horizontal bars (bottom left) display the total number of these genes per population. The matrix (bottom right) shows the populations under consideration as black circles. When an intersection between two or more populations is displayed, these populations are connected by a solid black line. The top histogram shows the number of genes under positive selection according to population or the intersections between two or more populations, as illustrated in the matrix plot below. The blue-highlighted bar represents the positive selection genes shared by DomC and DomD, and the upset plot of the selection patterns (hard or soft sweep) of these genes can be found in the upper right. (**B**) Positive selection along the genomes of cider and dessert cultivated apple. Different symbol shapes and y-axes represent different types of selection tests. From left to right, log_10_(*ω*) (squares), log_10_(*μ*) (circles), G12 (triangles), and G123 (crosses) values are shown. Significant selective sweep signals are represented in pink (DomC) or orange (DomD). The positive selection patterns of the gene symbols are represented in red (indicating hard sweeps) and green (indicating soft sweeps), consistent with those depicted in Fig. 3A. The blue text box represents the positive selection genes shared by DomC and DomC. See Supplementary Text for the functions of genes highlighted in this figure. DomC, Cider apples; DomD, Dessert apples; SiOr, *M. sieversii* and *M. orientalis*; SylE, Eastern European *M. sylvestris*; SylW, Western European *M. sylvestris*.

Wild populations, especially SylW, exhibited a higher number of genes under balancing selection than the cultivated apples (Dataset S4). Positive selection in wild groups targeted multiple NBS-LRR genes (SI Appendix, Fig. S25).

Selection during domestication acted on both a small set of shared domestication-related traits and many population-specific targets. DomC exhibited a higher proportion of soft sweeps and balancing selection, consistent with the retention of broader genetic diversity, whereas DomD experienced stronger directional selection. RNA-seq supported these signals: a substantial subset of sweep candidates were differentially expressed in cultivated apples compared to their wild relatives, providing orthogonal validation of the selected candidates.

### Local ancestry in selected regions

Local ancestry inference using RFMix (36) revealed enrichment for DomC and DomD in SiOr ancestry. Still, DomC carried more SylW ancestry in hard sweeps, whereas DomD carried more SylE ancestry across sweep types (SI Appendix, Figs. S26–S28). Q95_*SiOr, DomC/DomD, SylW/SylE (10%, 100%*)_ statistics (i.e., 95th percentile of derived frequencies in DomC/DomD of SNPs that have a derived allele frequency 100% in a donor panel - SylW/SylE -, but where the derived allele is at a frequency smaller than 10% in SiOr) (37) showed a bimodal frequency distribution in DomC and lower peaks in DomD (SI Appendix, Figs. S29–S31). These patterns reflect distinct adaptive introgression histories.

Integrating *f*_d_ statistics (SI Appendix, Figs. S32 and S33), topology weights (SI Appendix, Figs. S34 and S35), RFMix (SI Appendix, Figs. S32, S36), and Q95, we identified 14 adaptive introgression regions in DomC and 23 in DomD (SI Appendix, Dataset S5, Fig. 5; Figs. S37, S38). DomC haplotypes clustered on chromosome 17 overlapping a fruit texture QTL (38) (Fig. 4A), while DomD haplotypes localized on chromosome 16 and included the acidity marker *Ma* (39) (Fig. 4A and B).

**Figure 4.**
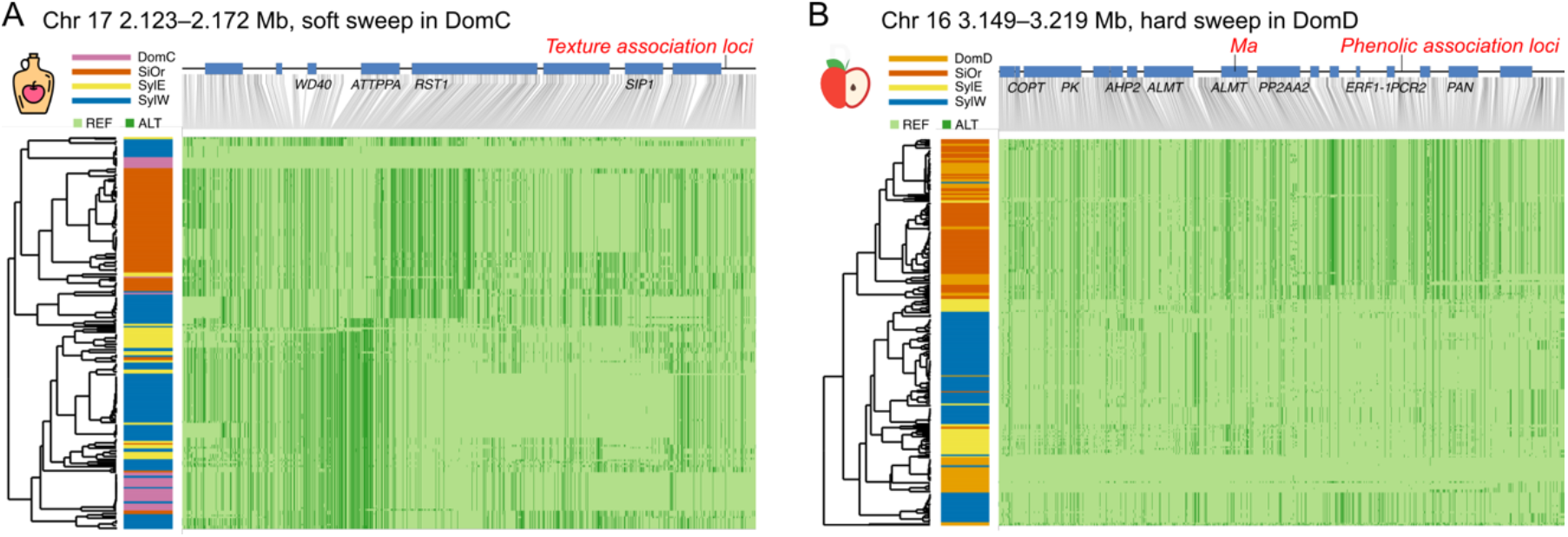
Two typical haplotype structures of adaptive introgression signals across positive sweep regions for cider and dessert apples. (A) Haplotype plot of the region on chromosome 17, spanning from 2.123 to 2.172 Mb with soft sweep in the cider apple genome. (B) Haplotype plot of region on chromosome 16, spanning from 3.148 to 3.220 Mb, hard sweep region of the dessert apple genome. The different colored bars on the left represent individuals from different populations. The left side is the dendrogram based on haplotypes clustering, and the bar plot represents the haplotypes from different populations in different colors. The upper part shows the genes in this region, and the gene IDs come from the annotations of the GDDH13 v1.1 genome (18). The QTL and GWAS regions are labeled in red above the gene models. DomC, cider apple; DomD, dessert apple; SiOr, *M. sieversii* and *M. orientalis*; SylE, Eastern European *M. sylvestris*; SylW, Western European *M. sylvestris*.

**Figure 5.**
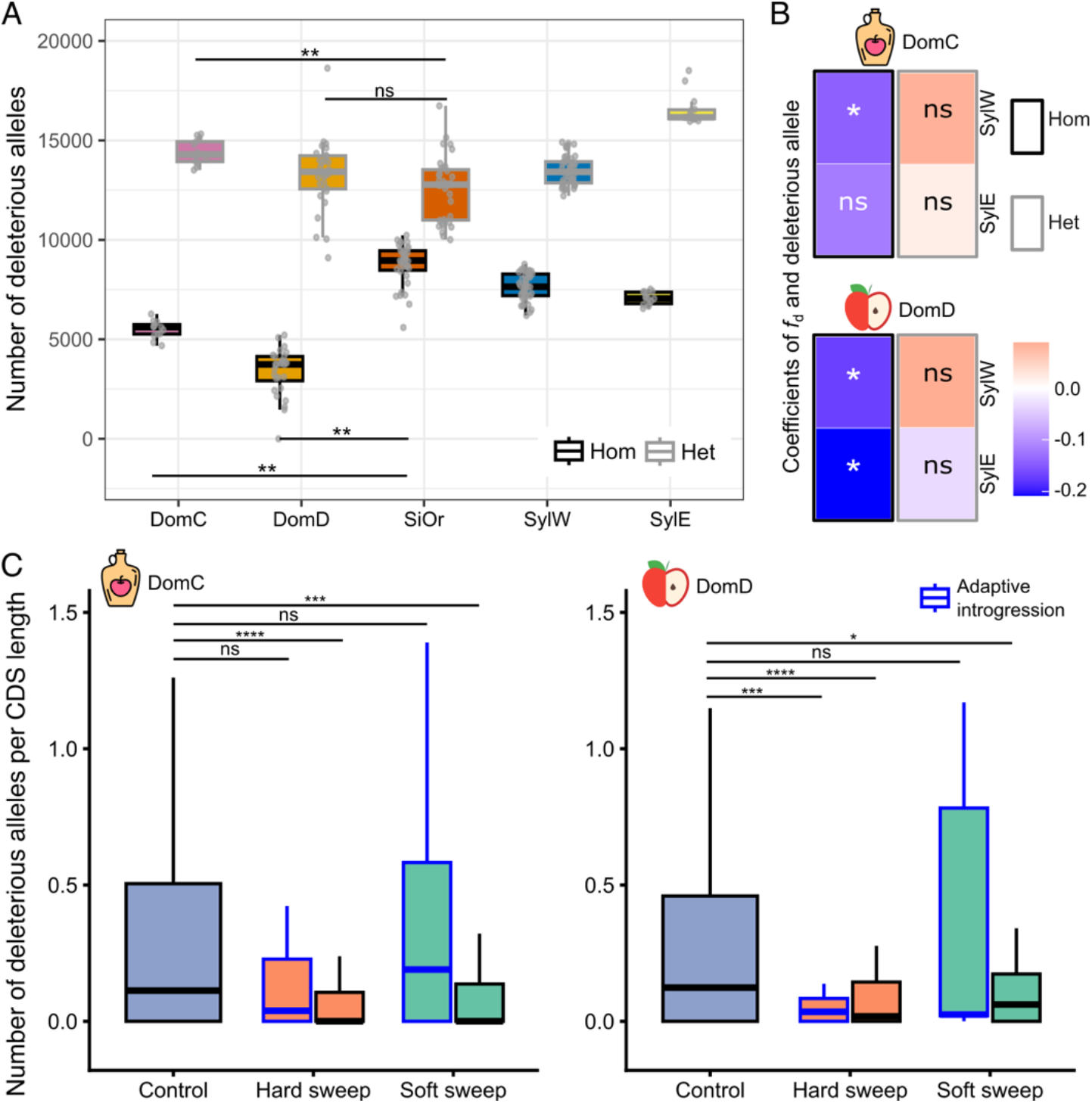
Distribution of mutation burden in cultivated and wild apple populations and the effect of introgression in cider and dessert apples. (A) Homozygote and heterozygote mutation burden across five apple populations, measured as the number of deleterious alleles in individuals within each population. (B) Heatmap of coefficients between *f*_d_ and the deleterious alleles. The population labels at the bottom and on the left are designated as P2 and P3 of *f*_d_ analysis. (C) Mutation burden in the cider (DomC) and dessert (DomD) apple populations across control genes, hard sweep genes, and soft sweep genes. Adaptive introgression genes are indicated with a blue border. The y-axis shows the number of deleterious alleles per 1000 bp of coding sequence (CDS). DomC, cider apple; DomD, dessert apple; SiOr, *M. sieversii* and *M. orientalis*; SylE, Eastern European *M. sylvestris*; SylW, Western European *M. sylvestris*. (ns: *p* > 0.05, ^*^: *p* ≤ 0.05, ^**^: *p* ≤ 0.01, ^***^: *p* ≤ 0.001, ^****^*p* ≤ 0.0001). Hom: homozygous state; Het: Heterozygous state.

Adaptive introgression shaped key fruit-quality loci, with cider and dessert apples drawing on different donor ancestries and experiencing distinct selection dynamics.

### Mutation load

No highly constrained sites with a GERP (40) score > 2 were detected in the apple genome (SI Appendix, Fig. S39), a threshold commonly used to identify deleterious mutations in constrained genomic regions (41, 42). Consequently, mutation load estimates relied on SIFT 4G predictions (43) (SI Appendix, Fig. S40). Cultivated apples carried fewer deleterious mutations than wild relatives. DomD showed the largest reduction (∼20.6%) compared to DomC (∼7.5%) (SI Appendix, Fig. S41). Most deleterious variants were heterozygous across populations (Fig. 5A, SI Appendix, Fig. S42).

Introgression from *M. sylvestris*, especially SylE into DomD and SylW into DomC/DomD, was associated with reduced homozygous deleterious mutations (Fig. 5B). Non-introgressed regions under hard and soft sweeps also had fewer deleterious mutations than controls (Fig. 5C). Hard sweep introgressed genes were depleted of deleterious mutations in DomD but not DomC. Domesticated apples—especially DomD—carry a lower deleterious load than wild relatives, likely due to the combined effects of selection and introgression.

### GWAS supports phenotypic divergence between cider and dessert cultivated apple

We ran a flowering-time GWAS in GAPIT v3 (BLINK/MLM; K and K+Q) on BLUPs from a subset of cider and dessert (44, 45). Two SNPs surpassed false discovery rate (FDR) and Bonferroni thresholds: Chr09:24,880,275 (P=1.11×10^−16^; PVE=92.4%) and Chr08:21,795,100 (P=1.51×10^−13^; PVE=87.7%) (Fig. S43). The Chr08 signal lies ∼2.6 Kb from *ACC1* (MD08G1178000), implicated in very-long-chain fatty acid biosynthesis and cold response (46), while the Chr09 SNP falls ∼10 Kb from an O-fucosyltransferase gene (MD09G1220000; SPINDLY-like hormone signaling (47)) in a region containing a HODOR transposable element (TEs) (ms537607_Chr09_HODOR-RLC_denovoMDO_kr-B-P814.288-Map5_reversed (18), suggesting potential regulatory effects. Genotype distributions at Chr09:24,880,275 clearly separate DomC from other groups, particularly *M. sylvestris* (Figs. S44–S45). Consistent with a wild-to-crop origin, *f*_d_ for DomC–SylW at this locus exceeds the genome-wide Q3 (the upper quartile of the *f*_d_ distribution), indicating localized introgression (Fig. S46). These two loci do not overlap sweep peaks, implying that flowering-time differences may stem from regulatory variation or polygenic bases not captured by classical selection scans.

## Discussion

Selection, introgression, and demography jointly forged the domesticated apple genome. Despite weak genome-wide differentiation at neutral sites and extensive shared admixture, cider and dessert cultivars of *M. domestica* form distinct gene pools that derive primarily from *M. sieversii*– *M. orientalis* rather than European *M. sylvestris* (13, 20). We find no evidence of a domestication bottleneck in *M. domestica*, as already observed in other perennials (3–5). Instead, demographic inference indicates a recent expansion in desert *M. domestica* and relative contraction in cider, alongside post-glacial expansions in *M. sieversii*–*M. orientalis* and Western *M. sylvestris*, with relative stability in Eastern *M. sylvestris* (48, 49). We accounted for these histories when inferring selection and in GWAS to limit confounding by relatedness and structure (44, 45).

Our results revealed contrasting selection modes in a perennial crop. Cider *M. domestica* shows proportionally more soft sweeps and balancing selection—consistent with broader retained diversity—whereas dessert *M. domestica* exhibits more hard sweeps on fruit-quality, disease-resistance, and flowering genes (15, 50–54). Wild populations, to a much larger extent than cider, carry massive balancing-selection signals, as expected in natural systems (55). Transcriptome profiling supports these inferences: a substantial subset of sweep candidates is differentially expressed in cultivated versus wild apples, indicating that selection has remodeled both sequence and expression.

In addition to selection, introgression from wild relatives played a central role in shaping diversity and adaptation. Local ancestry reveals a clear asymmetry: hard-sweep regions in cider are enriched for Western *M. sylvestris* ancestry, whereas dessert *M. domestica* carries more Eastern *M. sylvestris* ancestry across both hard and soft sweeps. Frequency spectra of introgressed variants further indicate contrasting dynamics: in cider apples, *M. sylvestris*–derived haplotypes cluster in hard-sweep tracts and show a bimodal distribution (many near fixation and many still rare), consistent with strong selection on a few advantageous tracts; in dessert apples, introgressed alleles more often remain at intermediate frequency and co-occur with soft sweeps, a pattern consistent with diffuse, polygenic adaptation in which recombination breaks up haplotypes and limits single-tract fixation (13, 56). A representative case is a lead flowering-time SNP on chromosome 9 that falls within an O-fucosyltransferase (SPINDLY-like) gene—an effector of hormone signaling linked to flowering pathways (47)—embedded in a cider–Western *M. sylvestris* introgressed tract and adjacent to a HODOR LTR element in the GDDH13 reference (18), suggesting cis-regulatory variation of wild origin may underlie phenological divergence. Notably, the flowering-time GWAS loci do not overlap classical sweep peaks, underscoring a regulatory and/or polygenic architecture that can elude hard-sweep scans.

Patterns of genetic load further distinguish perennial domestication from annual models. Cultivated apples—especially dessert *M. domestica*—carry fewer predicted deleterious alleles than wild relatives, contrary to trends in many annual crops and grapevine (3, 6, 7, 57, 58). Depletion of harmful variants in hard-sweep introgressed genes in dessert apples, together with higher burdens in soft-sweep regions, suggests that selection and wild gene flow jointly reduced genetic load while enabling adaptive change. While long-term clonal propagation could mask some deleterious alleles, the concordance between ancestry, selection mode, and load reduction argues for genuine purging in key regions.

Together, our integrative analyses—enabled by broad sampling of *M. sylvestris* across its range—refine wild-to-crop donor histories (13, 59, 60) and demonstrate that apple domestication combines classic hard sweeps (Mendelian-like) with widespread, polygenic adaptation from standing and introgressed variation—much of which escapes detection by standard sweep scans. Future steps include measuring genome-wide polygenic shifts among cultivar groups with more comprehensive sampling of *M. domestica*, particularly in Western and Northeastern Europe (60, 61). Concurrently, integrating allele-specific expression and methylation marks of nucleotide and structural variants—including TEs—will aid in distinguishing regulatory from coding regions involved in perennial domestication and improve prioritization of alleles for breeding and conservation amid global change.

## Materials and Methods

Newly sequenced samples were complemented with publicly available high-coverage genomes (62, 63) (SI Appendix, Table S1; SI Appendix, Fig. S1). DNA was extracted from young leaves using the Macherey–Nagel DNA Mini Kit® and quality-checked via agarose gel electrophoresis and Qubit quantification. Sequencing libraries were prepared with NEBNext® DNA Library Prep kits, checked for size distribution using an Agilent 2100 Bioanalyzer, and sequenced on Illumina platforms. RNA-seq data were generated for 59 leaf samples spanning cider, dessert, and wild genotypes sequences for short reads to examine domestication-related expression changes.

Reads were trimmed with fastp v0.21.0 (64), checked with FastQC v0.11.8 (65), and aligned to the *M. domestica* GDDH13 v1.1 reference genome (18) using bwa-mem v2.1 (66). Variant calling followed the GATK Best Practices workflow using HaplotypeCaller (19, 67). Variants were filtered in multiple steps (SI Appendix, Fig. S1) and refined with custom scripts. Genomic kinship was computed using Plink2 (68), and individuals with high genetic relatedness (kinship > 0.354) were excluded (69). Sites with more than 30% missing data were excluded. Phasing was performed using Beagle v5.1 (70), and recombination rates were estimated with MareyMap (71) based on high-density integrated genetic maps (72, 73). Pairwise linkage disequilibrium (LD) between SNPs were computed by PopLDdecay v3.42 (76) with decay distances ranging from 6.6 Kb to 123.5 Kb across populations (SI Appendix, Table S9). After filtering for synonymous and unlinked sites (Plink v1.9, vcftools v0.1.16) (74, 75), 31,300 high-quality SNPs were retained for downstream population and demographic analyses.

Population structure was inferred using fastSTRUCTURE v1.0 (21), with the optimal K determined by cross-validation and visualized with CLUMPAK (76) and plotted by pophelper v2.3.1 (77). Genetic clustering was also explored via PCA (Plink v1.9 (74); visualized in ggplot2 (78)) and neighbor-net analyses (Splitstree v4.12.3 (79)). Genetic diversity (*π*), differentiation (*F*_*ST*_), and divergence (*D*_*XY*_) were calculated using pixy v1.0.0 (80) in 10 kb windows across the genome.

The *Ne* dynamics were inferred using SMC++ v1.15.2 (81), assuming a mutation rate of 3.9 × 10^−9^ per bp per generation and a 10-year generation time (13). Coalescent species trees were inferred from 100-SNP windows using PhyML v3.0 (82) under the GTR model, re-rooted with *M. baccata* via the ETE toolkit (83), and summarized using ASTRAL-III v5.7.1 (84). Demographic modelling was based on the site frequency spectrum using fastsimcoal2 v2.8 (30). The best-fitting demographic model was identified on likelihood distributions obtained based on 50 expected SFS, each approximated using 100,000 coalescent simulations under the parameters that maximize the likelihood for each model.

Wild-crop gene flow was tested using *D*-statistics (ABBA–BABA test; (22, 27)) and *f*_4_-ratios (24) implemented in Dsuite v0.4 (85), as well as branch-specific tests (f-branch). Admixture graphs were explored using qpGraph in ADMIXTOOLS v2.0.4 (29). Chromosome-level introgression signals were examined with Twisst (27), sliding-window *fd* statistics(31), and DFOIL (26). Local ancestry was inferred with RFMix v2.03 (36).

Signatures of selection were identified using complementary methods: OmegaPlus (86) and RAiSD (87) for positive selection, G12/G123 (88) for soft sweeps, and BetaScan2 (89) for balancing selection. Neutral expectations were simulated with ms (98) under demographic models inferred by MSMC2 (90) to control for background demography. Candidate genes KEGG pathways were annotated with eggNOG-mapper v2.1.8 (91) and enriched using and ClusterProfiler v4.0 R package (92).

GERP++ (40) was used to identify conserved regions in *M. domestica* GDDH13 v1.1 reference genome. Mutation load was estimated using SIFT4G (43) for deleterious variant prediction, polarized with ancestral alleles inferred from *M. baccata* and *Pyrus pyrifolia* outgroups using est-sfs v2.0 (93). Mutation loads were summarized as derived deleterious allele counts per 1 Kb CDS length. The relationship between introgression, selection sweeps, and mutation load was assessed using Wilcoxon tests and generalized linear models (glm.nb in MASS R package (94).

GWAS for flowering time was conducted on a subset of 30 cultivated accessions (six replicates per accession in a random design, measured in three common gardens over three years) (Dataset S6). We used the GAPIT R package (v3.0) (44) implementing the BLINK method (45), with kinship (K) and population structure (Q) as covariates. SNPs were filtered for minor allele frequency (MAF ≥ 0.05) and missing rate < 20%. Significance thresholds were calculated using both FDR < 0.05 and Bonferroni correction (95). We examined nearby genes TEs using the GDDH13 v1.1 genome annotations (18) within the 10 Kb, corresponding to the average linkage disequilibrium (LD) decay estimations (SI Appendix, Table S9).

Full methodological details, including software parameters, filtering criteria, simulation strategies, and additional figures and tables, are provided in SI Appendix.

## Acknowledgments

The authors thank the INRAE Biological Resource Center “Pome Fruits and Roses” (RosePom: https://www6.angers-nantes.inrae.fr/irhs/Ressources-mutualisees/Ressources-genetiques/CRB-Fruits-a-pepins-et-rosier) and associated staff, especially the INRAE Horticulture Experimental Facility (Unité Expérimental Horticole: https://horti.angers-nantes.hub.inrae.fr), for maintaining the plant materials and associated datasets used in the present study. The authors thank Simon Martin for insightful discussions on detecting gene flow in genomes. We are grateful to the genotoul bioinformatics platform Toulouse Occitanie (Bioinfo Genotoul, https://bioinfo.genotoul.fr) for providing help, computing, and/or storage resources. USDA is an equal opportunity provider, employer, and lender. Mention of trade names or commercial products in this article is solely for the purpose of providing specific information and does not imply recommendation or endorsement by the U.S. Department of Agriculture.

## Funding

ATIP-CNRS Inserm

IDEEV

LabEx BASC

Tamkeen, under the New York University, Abu Dhabi Research Institute grant AD 454

European LEADER project 1.1.359 Proverbio

ANR-21-CE20-0005 PLEASURE

Ministry of Research, Innovation and Digitization, CNCS/CCCDI-UEFISCDI, project number PN-IV-P8-8.1-PRE-HE-ORG-2024-0223, within PNCDI IV

## Data, Materials, and Software Availability

All sequencing data generated in this study have been deposited in the NCBI under BioProject accession numbers PRJNA1252109 (whole-genome sequencing, Illumina platform) and PRJNA1297608 (RNA-seq data, Illumina platform).

The analysis code and workflow are available at https://github.com/CornilleEclecticLab/Apple-domestication_population_genomics.

## Supporting Information

Appendix 01

Datasets S1 to S6

