## Supplementary material for "Gene flow from the European wild apple and selection shaped the domesticated apple (*Malus domestica* Borkh.) genome": SI Appendix: BioRxiv_SI_AppendixDEF2.pdf

**This PDF file includes:**

Supporting text
Figures S1 to S46
Tables S1 to S8
SI References

**Other supporting materials for this manuscript include the following:**

Datasets S1 to S6

### Supporting Information Text

**Gene flow detected using five populations by $D_{FOIL}$.** We assessed the direction of gene flow using the five-taxon *D*-statistic ($D_{FOIL}$) approach, which counts the number of shared alleles based on a five-taxon symmetric phylogeny and tests for the occurrence and direction of gene flow among populations (1). The estimated SylW-SylE divergence time from SMC++ (2) was between the divergence time of DomC and DomD from SiOr. Consequently, we first performed $D_{FOIL}$ analysis using SiOr as P3, DomD or DomC as P4, and the two *M. sylvestris* groups as P1 and P2 (Fig. S10). $D_{FOIL}$ detected admixture in 9.3% and 9.4% of 100 Kb windows (mean ± s.d. = 569 ± 31 and 576 windows) in dessert and cider apples, respectively. Most of these windows exhibited gene flow between *M. domestica* and the ancestors of both *M. sylvestris* populations (P1⇄P4; 29.45% from DomD and 32.2% from DomC). We identified the second-highest number of introgression windows from SylW to *M. domestica* populations (P3→P2; 27.95% of DomD and 23.9% of DomC). The number of introgression windows from SylE to *M. domestica* populations was roughly half that from SylW (P4→P2; 12.1% of DomD and 12.5% of DomC). Given that there were no significant differences (*p*-value > 0.01, Fig. S8) in divergence times between SiOr–DomC or SiOr–DomD and SylE–SylW and based on the SMC++ split time results (Fig. S9), we also swapped P1–P2 and P3–P4. Using DomC or DomD as P4 revealed a similar pattern of gene flow (Fig. S11).

**Migration events detected by qpGraph.** We further investigated admixture among wild and cultivated apple populations using an admixture graph with 31,300 unlinked SNPs. We explored admixture graphs with 0 to 4 admixture events and conducted 200 iterations for each to identify the best graph (Fig. S14). The best fitting admixture graph (Fig. S14) had 3 migrations and was significantly (*p*-value < 0.05) better than all graphs with 0 and 1 migration, while not significantly worse than 6 graphs with two migrations, 22 graphs with 3 migrations, and 76 graphs with 4 migrations (Table S7).

**The gene flow detected by the *f*-branch statistics.** We based our analysis on $f_4$-ratios results to investigate gene flow events further and used the most common species-tree topology and the coalescent species-tree inferred by ASTRAL (Fig. S6) as input for *f*-branch analysis. The results indicate that SylE and SylW have strongly infiltrated the ancestral populations of DomD and DomC (Fig. S13).

**Topology weightings indicate gene flow between cultivated and wild apples.** The topology weightings were similar for both cider and dessert apples at the genome-wide level (Fig. S12). The common species tree topology (Fig. S18) accounted for 27.8% and 28.1% of the windows in cider and dessert apples, respectively. This was followed by the topology indicating gene flow from SylW to DomC and DomD (20.1% and 19.9%), and from SylE to DomC and DomD (15.1% and 15.5%). Other topologies represented 26.5% and 26.4% of the windows for DomD and DomC, respectively.

**Demographic modeling proved the gene flow events, but the origin of DomC is still uncertain.** To investigate gene flow between cultivated and wild apples and reconstruct the history of apple domestication. We designed eight scenarios based on various gene flow and species divergence events (Fig. S15). The models we compared included: (G0) no gene flow, (G1) gene flow only from the ancestor of DomC/DomD (SiOr) to SylW/SylE, (G2) gene flow only between DomC/DomD and SylW/SylE, and (G3) simultaneous gene flow as described in both G1 and G2. We conducted 100,000 coalescent simulations for each scenario, repeating the simulations 50 times. We then calculated the maximum likelihood values to compare models and identify the best-fitting scenario. The highest maximum likelihood was achieved for G3, in which G1 and G2 gene flow occurred simultaneously, showing a significant difference from the other gene flow scenarios (Fig. S16). However, the likelihood values of the two G3 scenarios did not exhibit significant differences (Fig. S16). Based on the estimated parameters of all G3 group scenarios, gene flow from the ancestor of DomD to DomD through SylE/SylW was greater than for other gene flow events. G3D1 indicated that the DomC diverged from SiOr at 4.75 thousand years ago. G3D2 indicated that DomC diverged from DomC at 2.40 thousand years ago. And all the demographic parameters are detailed in Table S8. And the schematic graph based on G3D1 and G3D2 scenarios were shown in Fig. 2D.

**Description of gene function under positive selection in the five apple populations.**
In DomD, genes under positive selection were primarily linked to fruit quality traits, as well as resistance to parasites and flower phenology (Details of genes under selection are provided in Dataset S2). Importantly, we detected a hard selective sweep within the QTL *fs15.1* on chromosome 15, encompassing *ERECTA* (MD15G1049300), a gene previously associated with fruit size (3, 4). Another hard sweep was identified in a cluster of four *SDH* genes on chromosome 1, including *SDH2* (MD01G1195200), which shows reduced expression during apple fruit development (5). On chromosome 5, a strong sweep was observed in MD05G1354900, encoding a phosphofructokinase (PFK) family protein linked to apple soluble sugar content (6). Additionally, on chromosome 6, a hard sweep was found in a *bHLH* gene (MD06G1199400), whose homologs are known regulators of acid accumulation in apples (7). We also identified a hard sweep in *MYB98* (MD06G1172900), a gene previously connected to flesh browning disorder in apples (8). Disease resistance is another important aspect of apple domestication. A gene encoding an NBS-LRR protein (MD15G1042600) and one encoding an LRR protein (MD01G1181100) were under hard and soft selection in DomD, respectively. Two *WRKY* genes (MD07G1234600, *WRKY55*; MD07G1234700, *WRKY70*) were also under soft selection and played important roles in defense against apple tree canker disease (9). We also identified up to eight flower-related genes under positive selection, involved in general signaling, sugar metabolism, and photoperiodism/light perception pathways. Four of these genes underwent soft selection, while the other four showed evidence of hard selection. Notably, three flower-related genes on chromosome 16 (MD16G1018000, *FKF1*; MD16G1047200, *PFT1*; MD16G1047700, *TEM1*), which are under hard selection, are specifically associated with photoperiodism and light perception pathways (10). In DomC, genes under positive selection were also associated with fruit quality but differed from those in dessert apples. Notably, we did not detect *fs15.1*, the gene linked to fruit size in dessert apples. While cider apples are larger than crabapples (e.g., *M. baccata*), they remain smaller than dessert apples, supporting a multi-step domestication model (11, 3). We identified two *SAMT* (MD15G1023600 and MD15G1023700) genes under hard selective sweeps on chromosome 15, which are involved in the biosynthesis of methyl salicylate, a key volatile compound contributing to tomato aroma (12). Additionally, two cytochrome P450 genes, MD15G1057500 and MD15G1057600, homologous to *CYP711A1/MAX1* in *Arabidopsis*, were under hard sweep and are implicated in strigolactone biosynthesis and the flavonoid pathway (13). A hard selective sweep was also detected in *MdXTH18* (MD13G1016500), a gene associated with fruit firmness (14). Several genes under positive selection in DomC were functionally linked to auxin regulation. These include three *TMK1* genes (MD16G1015000, MD16G1015100, MD16G1015200), which are key regulators in auxin signaling in root growth(soft sweeps) (15); two *ASB1* genes (MD13G1016400, MD16G1013900), involved in IAA biosynthesis (hard and soft sweeps) (16); one *PILS2* gene (MD16G1030900), an auxin efflux carrier protein (17); and two *MdGH3* genes (MD07G1145900, MD13G1132300), which are early auxin-responsive genes (soft sweeps) (18). We also identified three miRNAs (one *MiR-166*, two *MiR-156*) under soft selection and one MiR-159 under hard selection in DomC. While *MiR-159* was also shared with the DomD population soft selection genes. Furthermore, we found only two flower-related genes under hard selection and one under soft selection in DomC, significantly fewer than in DomD (Dataset S2). In the three wild apple populations, genes under positive selection were primarily associated with resistance and developmental processes (Fig. S25, Dataset S2, and Dataset S3). The *SiOr* population exhibited limited signatures of positive selection, with only 32 detected genes (Fig. 3A). Among them, one resistance gene (*CNL*, MD04G1041500), and two genes related to root development (*ASPR1*, MD04G1217500 and *XEG113*, MD06G1008500), under hard positive selection(19, 20). The number of genes under positive selection in *SylE* was also limited (Table 1 and Dataset S2). Hard sweep regions included three clustered *CRK* genes (MD01G1096800, MD01G1096900, MD01G1097000), known regulators of plant stress responses (21) as well as a *TNL* resistance gene (MD05G1136700). In contrast, *SylW* exhibited signatures of selection in hormone-related genes, including *GAS2* (MD07G1074700) and *KAO1* (MD07G1083200), which regulate gibberellin biosynthesis (22), *EOL1* (MD03G1258600), involved in ethylene signaling (23), and *PCN* (*MD08G1080300*) gene, which integrates auxin signaling (24). Furthermore, SylW exhibited hard positive selection in four genes associated with plant defense or immune responses. These genes

include *PUB23 (*MD07G1083300*)*(25), *FER (*MD09G1069300*)* (26)*, GFS12 (*MD13G1143700*)* (27), and *LIK1 (*MD10G1273300*)* (28)*.* Soft sweep regions in *SylW* included a *CNL* gene (MD17G1204900) and *SES1* (*MD02G1074400*), involved in salt and heat stress responses (29). SylW and SylE shared four specific hard sweep genes: two wall-associated kinase-like genes, *RFO1* (MD09G1145200), which provides broad-spectrum resistance to *Fusarium* species, and *SKB1* (MD15G1111800), which influences flowering by regulating the *FLC* pathway. Between cultivars and wild populations, only one hard sweep gene was identified between SiOr and DomC populations (Fig. S30). Soft selection genes were largely unique to each population, with just one overlapping gene (*SWEETIE*-like) found between SiOr and SylW (Fig. 3A).

**Description of gene function under balancing selection in the five apple populations.**
In addition to the balancing selection genes described in the main text, SylW exhibited the highest number (32 genes) of unique genes under balancing selection (Fig. S21), including an *RVE8* (MD12G1126100), regulation of circadian clock gene (30), a *TNL* disease resistance gene, and others related to critical biological processes such as plant growth, development, response to environmental stress, and cell signaling, demonstrate the complex adaptation and regulatory mechanisms of SylW to its environments (Dataset S4). We also found an *F-box* gene under balancing selection only shown in the SylE and SylW populations (Fig. S24). In contrast, the cultivar populations possessed only a few unique balancing selection genes (Fig. S22). DomD only had one unique gene (MD06G1060700) with an unknown function, while SylE had two, including a root development gene, *XEG113* (MD06G1008500) (20), and a cellulase protein gene (MD13G1104700). Other genes are involved in plant biological processes such as metabolism, transcription, signal transduction, disease resistance, and material transport (Dataset S4).

Because gene functional annotation relies on inference based on homology, it is essential to acknowledge that these apple genes need further validation to confirm their actual roles.

### Materials and Methods

**Sample preparation, DNA extraction, and sequencing.**
A total of 140 newly sequenced individuals originated from leaves of 43 cultivated apple samples, including 21 dessert and 22 cider apple varieties, 75 *Malus sylvestris* (European crabapple), nine *Malus orientalis*, three *Malus sieversii,* and ten *Malus baccata*. The status of cider and dessert cultivars was assessed using the New Book of the Apple (31) and the expertise of the ResPom consortium (INRAE Angers). DNA was extracted with the Macherey-Nagel DNA Mini Kit® using manufacturer instructions. DNA quality controls were performed using two methods: (1) Agarose gel electrophoresis for testing DNA degradation and potential contamination and (2) Qubit® 2.0, which quantifies the DNA concentration precisely. A total amount of 1.0μg DNA per sample was then used as input material for the DNA library preparations. Sequencing libraries were generated using NEBNext® DNA Library Prep Kit following the manufacturer's recommendations, and indices were added to each sample. The genomic DNA was randomly fragmented to a size of 350bp by shearing, then DNA fragments were end polished, A-tailed, and ligated with the NEBNext adapter for Illumina sequencing, and further PCR enriched by P5 and indexed P7 oligos. The PCR products were purified (AMPure XP system), and resulting libraries were analyzed for size distribution by Agilent 2100 Bioanalyzer and quantified using real-time PCR. After pooling, the qualified libraries were fed into Illumina sequencers according to their effective concentration and expected data volume.

This newly sequenced dataset was complemented with publicly available sequencing data with at least 15X coverage (78 other genotypes in (32, 33), including 24 dessert and one cider apple cultivars, one *M. orientalis*, 37 *M. sieversii*, 12 *M. sylvestris* and three *M. baccata*).

Therefore, the total dataset included 218 genotypes and was composed of 45 dessert and 23 cider apple varieties and the four main wild relatives of the cultivated apple: *M. orientalis* (*N*=10), *M. sieversii* (*N*=40), *M. sylvestris* (*N*=87), and *M. baccata* (*N*=13) (Dataset S1 and Fig. S1). All plant material used in this study was collected or sourced in full accordance with applicable international regulations, including the Nagoya Protocol and the International Treaty on Plant Genetic Resources for Food and Agriculture (ITPGRFA). Proper permits and agreements were secured for the collection, use, and analysis of both cultivated and wild apple genetic resources.

**Variant calling and filtering**

Quality control and pre-processing of the reads were performed using FastQC v0.11.8 (34) and fastp v0.21.0 (35), respectively. Variant calling was carried out following the Best Practices procedure (36). Clean DNA reads were aligned to the GDDH13 Version 1.1 reference genome (37) using bwa-mem v2.1 (38) (https://github.com/bwa-mem2/bwa-mem2). Mapping statistics are shown in Table S1. Raw variants were called using GATK HaplotypeCaller (39) (GATK version 4.1.7, https://github.com/broadinstitute/gatk). The raw variants were filtered using the GATK variant filter module with a hard filter (Level 1 filters) and further refined with a custom script (Level 2 filters) (Fig. S1). Genomic kinship was calculated for all pairs of individuals using Plink2 (40). Duplicate or clonally related individuals were excluded from further analysis (i.e., individuals with pairwise KING-robust kinship estimates greater than 0.354) (41), using 'Plink2 –make-king'. Individuals with a missing rate > 30% were also removed. All filtering parameters are listed in Table S2, and the removed samples are detailed in Dataset S2, representing 17 genotypes and retaining 201 individuals. The dataset was phased using Beagle v5.1 (42) with default settings and genetic maps (43) for each chromosome. The genetic map position of each SNP was obtained by MareyMap pipeline. The recombination rate variation across the genome was estimated using high-density integrated genetic maps (53, 14). We downloaded the data of the marker genetic distance and physical distance on GDDH v1.1 reference genome (7) and imported it into MareyMap Online (54) (https://lbbe-shiny.univ-lyon1.fr/MareyMapOnline/), then followed MareyMap pipeline and get recombination rates along the genome.

Pairwise linkage disequilibrium (LD) between SNPs was computed by PopLDdecay v3.42 (44). The decay distance of LD was defined as the physical distance at which $r^2$ decreased to half of its maximum value. The LD decay distance varied among populations, ranging from 6.6 Kb to 123.5 Kb (Table S9). For genetic diversity ($\pi$) (45), genetic distance ($D_{XY}$) (45), differentiation ($F_{ST}$) (46), and $f_d$ scans analysis, a window size of 10 kb was chosen, providing a compromise between resolution and statistical robustness given the SNP density across the genome. For the $D_{FOIL}$ analysis, a window size of 100 kb was used because the information content in five individual sets of 10 kb windows was insufficient.

Linked and non-synonymous SNPs were filtered out for population structure and demographic inferences to avoid bias due to potential SNPs linked to genomic regions under selection (Level 3 filters, Table S2). SNPs were annotated with SnpEff v5.0 (47), and only synonymous SNPs were retained. SNPs in LD were removed. Linked SNPs were filtered using Plink v1.09 (48) with the '–indep-pairwise' function, applying the following parameters: window = 5 Kb, step = 1 SNP, and $r^2$ = 0.2. SNPs with distances less than 5 Kb were removed using the 'thin' function of vcftools v0.1.16 (49). In total, 31,300 unlinked and synonymous SNPs were used for population structure, genetic variation, and differentiation analyses on 201 genotypes.

**Population structure, genetic variation, and differentiation**

The population structure and admixture were inferred using the Bayesian clustering method implemented in fastSTRUCTURE v1.0 (50) using the 31,300 unlinked and synonymous SNPs and the 201 genotypes. Analyses were performed with default parameters, running 20 iterations for each $K$ value from $K$ = 1 to $K$ = 10 clusters. The consensus solution for each $K$ value was generated using CLUMPAK (51), and the optimal $K$ value was determined based on the cross-validation error (50) . The bar plots were visualized using the R package pophelper v2.3.1 (52), and the $K$ value that resulted in well-assigned individuals for all clusters was selected.

Two additional methods were employed to further explore the genetic variation and differentiation among the genetic groups detected by fastSTRUCTURE. Principal component analysis (PCA) was conducted using the Plink v1.9 PCA module (48), and the first three principal components were visualized using the R package "ggplot2" (53). A Hamming distance matrix among individuals was calculated using Plink v1.9 (48) , followed by the construction of a neighbor net, which was visualized with Splitstree v4.12.3 (54). For both PCA and neighbor-net, individuals assigned to a given cluster with a membership coefficient ≥ 0.8 (Figure S4) were color-coded according to their genetic cluster inferred with fastSTRUCTURE, while admixed individuals were colored gray. This threshold was selected based on the distribution of maximum membership coefficients inferred by fastSTRUCTURE (see Results, Fig. S4).

**Population genetic diversity and differentiation**

Descriptive population genetic estimates were computed for each population (i.e., each cluster inferred with fastSTRUCTURE, excluding admixed individuals with a membership coefficient < 0.80). Genetic diversity ($\pi$) (45), genetic distance ($D_{XY}$) (45), and differentiation ($F_{ST}$) (46) between populations were estimated using pixy (55) with the "all-sites" variant calling format (VCF). Population genetic diversity statistics—including the number of private alleles ($A_p$), number of polymorphic sites (S), observed heterozygosity($H_o$), expected heterozygosity ($H_e$), and inbreeding coefficient ($F_{IS}$)—were calculated for each population using the *populations* module in Stacks (56). Genome-wide $\pi$, $D_{XY}$, and $F_{ST}$ were calculated using all sites (332,335,748 variant and invariant sites) in 10 Kb window regions of high mappability.

**Divergence time estimates and effective population size (*Ne*) over time**

To infer population divergence history, we first reconstructed a coalescent species tree using ASTRAL-III (57). ASTRAL is a tool for inferring an unrooted species tree from a set of unrooted gene trees under the multispecies coalescent model. This method is particularly suited for addressing incomplete lineage sorting, a common phenomenon in tree species (58). Phylogenetic trees per 100 SNP windows were calculated with PhyML (59) with the GTR model. The output trees were re-rooted with one Bacc individual by the ETE Toolkit (60), which were used as input for ASTRAL-III. These trees were also used as the input for Twisst (61) to get the most common species-tree topology of cultivated and wild apple populations.

Effective population size (*N*e) over time and population divergence estimates were inferred using two methods: SMC++ (2). First, the reference genome mappability scores were calculated with GenMap v1.3.0 (62) using a *k*-mer size of 140 (the average length of filtered sequencing reads) for 20 Kb windows with 10 Kb steps. Any mean scores of 20 Kb higher than 0.8 were defined as high mappability regions. SMC++ demographic inference employed SNPs within high mappability regions with all individuals for each population. A mutation rate of $3.9 \times 10^{-9}$ per bp per generation and a generation time of 10 years were used to convert generations into years (63). The divergence time between DomD and SiOr, DomC and SiOr, and SylW and SylE was also inferred by *SMC*++ using the "*split*" command.

**Genome-wide estimates of gene flow among wild and cultivated populations**

*D*-statistics and $f_4$-ratio were employed to detect gene flow among cultivated and wild populations. The *D*-statistic (ABBA-BABA test) (61, 64) and related statistics, such as the $f_4$-ratio (65), are widely used parsimony-like methods for distinguishing gene flow from incomplete lineage sorting. While the *D*-statistic tests for introgression, the $f_4$-ratio estimates the proportion of admixture or introgression using population allele frequencies. We computed genome-wide four-taxon *D*-statistics and $f_4$-ratios using the Dtrios command in Dsuite (66) with unlinked 31,300 SNPs. Tests were performed on populations identified by fastSTRUCTURE, excluding admixed individuals with membership coefficients <0.80, and using Bacc as the outgroup. *D*-statistics and $f_4$-ratios were computed for all possible trios and organized according to the tree topology. The significance of *D*-statistics was assessed using a jackknife over 20 blocks (67, 64). Results were visualized using the Ruby scripts plot_d.rb and plot_f4ratio.rb available from M. Matschiner's repository. After that, *D*-statistics and $f_4$-ratios were calculated for each branch of the population tree using the "Fbranch" module in Dsuite using the most common species-tree topology and the coalescent species-tree inferred by ASTRAL. The *f*-branch result was visualized using the dtools.py script provided in the Dsuite.

We investigated the directionality and extent of gene flow with *qpGraph* implemented in the R package ADMIXTOOLS2 (68). As input, allele frequencies and $f_2$-statistics (65) were calculated for each population using the 31,300 unlinked and synonymous SNPs. The function "find_graphs" in qpGraph was then executed to explore admixture graphs with the assumed number of admixture events ranging from 0 to 4 using Bacc as the outgroup population. A testing procedure was utilized where the best scoring graph out of 200 candidate graphs for a given number of migrations was first identified, and graphs with the same number of migrations were not significantly worse ($p$-value > 0.05). It was also tested whether the best scoring graph for each number of migrations could be significantly rejected in favor of a graph with a higher number of migrations. The test score was calculated by optimizing a topology from a subset of the $f$-statistics and evaluating it on the remaining data. A test of significance was performed using a jackknife approach for each obtained graph, which was then compared to the remaining graphs using a nominal $p$-value of 0.05. The best number of admixture events was determined by identifying the best-scoring graph significantly better than all graphs with fewer migrations (Table S7).

To further characterize the gene flow between cultivated and wild apple, full genome SNPs (28,377,551) were used for $D_{FOIL}$, Twisst, and chromosome-level $D$-statistic, and $f_d$ genome scan analysis. The $D_{FOIL}$ analysis (1) was conducted on a selection of five populations (*P1, P2, P3, P4, O,* and the divergence time between P1 and P2 occurred earlier than that between P3 and P4). The *M. baccata* population was used as the outgroup (*O*). The SMC++ divergence time estimates suggests that the SylW and SylE divergence time was earlier than DomC from SiOr but later than DomD from SiOr. So, all four possible populations set, set1 (P1 = SiOr, P2 = DomC, P3 = SylW, P4 = SylE, O = Bacc), set2 (P1 = SiOr, P2 = DomD, P3 = SylW, P4 = SylE, O = Bacc), set3 (P1 = SylW, P2 = SylW, P3 = SiOr, P4 = DomC, O = Bacc), and set4 (P1 = SylW, P2 = SylW, P3 = SiOr, P4 = DomD, O = Bacc) were used as the input for $D_{FOIL}$ analysis. $D_{FOIL}$ analysis was applied by the mvftool v0.6.2.1 (69) 'CalcPatternCount' function. A total of 100 non-repeating individual sets were randomly generated, and the test was applied on 100 KB-sized windows along the genome. The total number of significant introgression windows in each set was then summarized. The Twisst analysis (61) was applied to two groups of five populations: Set DomD, comprising SiOr, DomD, SylE, SylW, and Bacc; and Set DomC, consisting of SiOr, DomC, SylE, SylW, and Bacc. The analysis was constrained to a five-taxon topology. Phylogenetic trees were inferred for windows containing 100 SNPs by PhyML (*75*) using the same parameters as above, which served as the input data. Chromosome-level topology weighting analysis was applied separately. Chromosome-level $D$-statistics were calculated for each chromosome using the Dtrios command in Dsuite in four population groups (SiOr-DomC-SylE-Bacc, SiOr-DomC-SylW-Bacc, SiOr-DomD-SylE-Bacc, and SiOr-DomD-SylW-Bacc). To identify introgression within specific genomic regions, we also employed the $f_d$ statistic in 10-kb sliding windows using the ABBABABAwindows.py (https://github.com/simonhmartin/genomics_general/tree/master) on the same four population groups in Chromosome-level $D$-statistics analysis. We identified the potential introgressed regions as those falling within the top 5% of $f_d$ values across four population groups and extracted the genes located in these areas. The list of genes was then subjected to KEGG enrichment analysis, and the significant KEGG pathways were visualized by R (see details in 'Gene function annotation and enrichment analysis' section).

**Demographic modeling based on site frequency spectrum (SFS)**

Based on the inferred topology from qpGraph, the most common ASTRAL phylogenetic tree, and the gene flow events identified using $D$-statistics and $D_{FOIL}$, fastsimcoal v2.8 (70) was employed to further investigate the demographic history of apple populations. The analysis considered four possible gene flow scenarios: G0, where no gene flow occurs within populations; G1, involving gene flow between the ancestors of DomC/DomD and SiOr, as well as SylW/SylE populations; G2, where gene flow occurs between DomC/DomD and SylW/SylE populations; and G3, which combines both G1 and G2. Additionally, four divergence scenarios were also included; D1, where DomC and DomD both diverge from SiOr, with DomC diverging more recently and SylE diverging from SylW; D2, where DomD diverges from SiOr, followed by DomC diverging from DomD, and SylE diverging from SylW; D3, where DomC and DomD both diverge from SiOr, with DomC diverging more recently, and SylW diverging from SylE; D4, where DomD diverges from SiOr, followed by DomC diverging from DomD and SylW diverging from SylE. This framework resulted in a total of 16 distinct scenarios being evaluated (Fig. S15).

A folded 2D site-frequency spectrum (SFS) was generated using 31,300 SNPs with the program easySFS (using the -a option) (71, 72). Each scenario was optimized through 50 independent runs with specific settings, including 100,000 coalescent simulations for each likelihood estimation (-n 100,000), 40 cycles of the conditional maximization algorithm (-L40), and a minimum of 10 observed SFS entry counts for likelihood computation (-C10). The maximum likelihood values obtained from these runs were used to determine the best-fitting scenario or scenario group. Once the most likely scenario or scenario group was obtained, the parameter median estimates from the 50 runs and calculate the 95 percentile confidence intervals.

**Detection of signatures of positive and balancing selection**

Signatures of positive selective sweeps in the genomes of cultivated and wild apples were detected using OmegaPlus (73) and RAiSD (74), which are a linkage disequilibrium (LD) method and a composite method, respectively, employing 28,377,551 SNPs (Fig. S1). OmegaPlus computes the $\omega$ statistic, which can be used for synonymous and non-synonymous linked SNPs while considering the LD between SNPs. For the selective sweep analysis, OmegaPlus was applied to each chromosome individually per population with the parameters -minsnps 5, −minwin 5,000, −maxwin 100,000, and a grid size set to the number of chromosome lengths divided by 1000 bp (chosen given a median gene length of ~2,000 bp, to ensure coverage of all genes for detection). RAiSD calculates the $\mu$ statistic, a composite evaluation test that scores genomic regions by quantifying local polymorphism reduction, shifts in the SFS towards low- and high-frequency variants, and localized patterns of LD across the genome. We performed $\mu$-statistics using 50 SNPs per chromosome per population. Soft sweeps were detected using the *G12* and *G123* statistics (75). Positive selection can reduce genomic diversity at both the selected and linked sites, producing a characteristic signature of elevated expected haplotype homozygosity. These selective sweeps can be classified as either hard or soft. In a hard selective sweep, a single adaptive haplotype rises to high population frequency. In contrast, in a soft sweep, multiple haplotypes sweep through the population simultaneously, producing distinct patterns of genetic variation near the selected site. VCF files were converted to the format required by SelectionHapStats (https://github.com/ngarud/SelectionHapStats). The *G12* and *G123* statistics were calculated for each chromosome of each population using overlapping windows of 200 SNPs with a step size of 50 SNPs. The above threshold definitions for these positive selection signals are referenced in the "Control of demographic history for selection analyses" section.

Signatures of balancing selection were inferred using BetaScan2 (76), which identifies the enrichment of variants in regions with low deviation from allele frequency and a deficit of substitutions. VCF files were converted to the input format required by BetaScan2 using a custom script, excluding folded allele frequencies lower than 0.15 (-m 0.15). The $\beta$ scores were calculated for all remaining SNPs using a window size of 1000 bp (-w 1000) for each SNP position. LD blocks were defined as regions with LD ($r^2 > 0.8$), computed using LDBlockShow v1.39 (77). Similar to the positive selection analysis, neutral model to define the threshold of $\beta$ scores were applied.

**Control of demographic history for selection analyses**

The effect of population demographic history on the SFS, LD, and genetic diversity along the genome was controlled for all selection analyses presented above. Significance thresholds for deviations from the neutral model in selection analyses were assessed based on simulated datasets under the neutral demographic model. The population history (*Ne*) estimated by MSMC2 (78) was converted into ms parameters using the msmc2ms.py script (https://github.com/stschiff/msmc-tools). A total of 10,000 datasets, each including five populations of the same number of pure individuals defined by fastSTRUCTURE analyses, were simulated with the ms program (79). Each dataset consisted of a 1 Mb chromosome fragment from the respective model parameters. Subsequently, 10,000 simulations of 1 Mb chromosome under the neutral demographic model were conducted to assess the significance of $\omega$, $\mu$, G12/G123, and $\beta$ statistics using the same parameters for the real genome. The top 1% value for $\omega$, $\mu$, G12/G123, and $\beta$ was considered to significantly deviate from the neutral model simulations, thus serving as thresholds for each selection analysis. The low mappability regions (see above) were excluded from the selection signal results.

**Gene function annotation and enrichment analysis**

Enrichment analyses of genes under selection were then performed. KEGG pathway annotations were obtained using eggNOG-mapper v2.1.8 (80), with the GDDH v1.1 genome gene protein file (37) as input. KEGG pathway names were downloaded from https://www.kegg.jp. Enrichment analysis was conducted using the ClusterProfiler v4.0 R package (81), with significantly enriched KEGG pathways identified as those with a *q*-value < 0.05. Enrichment plots were generated using the Enrichplot v3.17 R package (82). Flowering genes were identified by aligning the GDDH v1.1 protein sequence with the Flowering Interactive (FLOR-ID) database(10). Genes were retained if they exhibited an e-value of less than 1e-5, matched bases of at least 50 base pairs, and an identity of 80% or higher. Transcription factors (TFs) were identified in the GDDH v1.1 genome using PlantTFDB (http://planttfdb.cbi.pku.edu.cn/) (83) by HMMER3.0 (84).The TFs were classified according to the consensus rules, including the required and prohibited protein domains for each TF gene family, summarized on the PlantTFDB website. Resistance gene analogs (Rgene) based on differences in their domains identified by NLGenomeSweeper (85), combine the information from the GDDH v1.1 released annotation(37).

**RNA extraction, library preparation, and sequencing**

Leaf samples were immediately flash-frozen in liquid nitrogen after harvesting and subsequently stored at –80 °C. In total, 59 leaf samples were collected from cider *M. domestica* (n = 15, five genotypes * three clonal replicates), dessert *M. domestica* (n = 15, 5 genotypes * three clonal replicates), *M. sylvestris* (n = 15, five genotypes representing SylW * three clonal replicates), and *M. sieversii* (n = 12, four genotypes * three clonal replicates). Samples were pulverized using a QIAGEN TissueLyser II (R2) with the addition of Polyvinylpyrrolidone (PVP40) to reduce interference from phenolic compounds. Total RNA was extracted using the Nucleospin RNA Plant Kit (Macherey-Nagel, Düren, Germany), following the manufacturer's protocol, which includes RNA binding, DNase treatment, and centrifugation steps. Again, all plant material used was collected or sourced in full accordance with applicable international regulations, including the Nagoya Protocol and the International Treaty on Plant Genetic Resources for Food and Agriculture (ITPGRFA). Proper permits and agreements were secured for the collection, use, and analysis of both cultivated and wild apple genetic resources.

RNA quality and quantity were initially assessed using a NanoDrop spectrophotometer®, with purity determined via absorbance ratios at 260/280 nm and 260/230 nm. Samples with A260/A230 ratios between 1.8 and 2.2 were considered of poor quality and discarded. RNA integrity was further evaluated using a Bioanalyzer (Agilent Technologies, USA). High-quality RNA was used for cDNA library construction with the Novogene NGS Stranded RNA Library Prep Set (PT044), along with rRNA depletion. Sequencing was performed on a NovaSeq 6000 system (Illumina, San Diego, CA, USA) using the S4 Reagent Kit v1.5 (300 cycles), producing 150 bp paired-end reads targeting 20 million reads per samples.

**RNA-sequenced data processing and differential expression analysis**

Raw reads were assessed using FastQC v0.11.2 (34), followed by adapter trimming and quality filtering with Fastp v0.23.2 (35), and ribosomal RNA filtering using SortMeRNA v4.3.2 (86). Cleaned reads were aligned to the *M. domestica* GDDH13 v1.1 reference genome (14) using STAR v2.7.10b (87), and gene quantification was performed in alignment-based mode using Salmon v1.10.0 (88). Differential gene expression analysis was conducted with the R-based DiCoExpress pipeline (89). Read counts were first normalized by collection year and then merged. Low-expression genes were filtered using the "NbConditions" strategy with a CPM_cutoff of 5. TMM normalization was applied (90), and differential expression was tested using the edgeR method (91) with an FDR threshold (Alpha_DiffAnalysis) of 0.05. Genes differentially expressed between cultivated (cider or dessert) and wild apples were cross-referenced with genes under positive selection in DomC and DomD to identify domestication-related candidate genes.

**Genomic landscape of introgressions and adaptive introgressions**

The genomic landscape of introgression was investigated as a first step to studying the adaptive introgression. The $f_d$ statistic was calculated using 100-SNP sliding windows with ABBABABAwindows.py (https://github.com/simonhmartin/genomics_general) to detect introgression regions across the whole genome. The analysis considered the models (((SiOr, DomD or DomC), SylW or SylE), Bacc). Topological weighting was applied to examine local genomic topological relationships using Twisst (61). Sub-trees were constructed with PhyML (59) under the GTR model based on phased data in 100-SNP windows. A smoothing approach was then applied, using a 20 Kb window size to refine the inferred topologies. The resulting topology weighting data were categorized into five distinct patterns (Fig. S18): no introgression (one topology), DomC/DomD introgressed by the ancestor of SylE and SylW (one topology), SylE introgression (three topologies), SylW introgression (three topologies), and others (seven topologies).

To investigate adaptive introgression further, two new analyses were introduced. The Q95 statistic (92) was first calculated in the form of $Q95_{SiOr,\ DomC/DomD,\ SylW/SylE\ (10\%,\ 100\%)}$, which pertains to the 95% quantile frequency of derived alleles in the DomC or DomD populations that have a derived allele frequency of 100% in a donor panel in the SylW and SylE individuals and are present at <10% frequency in the combined SiOr panel. Local ancestry was inferred using RFMix (v2.03) (93) with phased data from phased sites of all DomC and DomD individuals. The Bacc, SiOr, SylW, and SylE populations, retaining only very pure individuals (membership coefficient > 0.99 in fastSTRUCTURE analysis), served as reference populations. A comprehensive analysis integrating results from four analytical methods— $f_d$, topology weighting, Q95 statistic, and local ancestry inference—was conducted to identify potential regions resulting from adaptive introgression. For each selection region, $f_d$, introgression topology weighting, and SylW/SylE ancestry proportion were calculated. Regions exhibiting outlier values based on the interquartile range (IQR) method were subsequently extracted, with thresholds for extreme values defined as (Q3 + 1.5IQR) to identify outliers (Fig. S32 to Fig. S35 and Fig. S36). The distribution of Q95 values exhibited a bimodal pattern, prompting using the k-means algorithm to categorize the signals into two clusters. To prevent overinterpretation of the Q95 statistic's efficacy, the cluster centroid value of the higher Q95 values was utilized as the threshold (Fig. S31). Any selected region containing two or more signals (including signals of the same type but from different wild donor populations) was designated as an adaptive introgression area based on the four identified signals.

**Mutation load estimates**

The constrained regions of the apple genome were identified using GERP++ software (94). First, genome sequences of eight species within the Rosales order (i.e., apple (*M. domestica*), pear (*Pyrus pyrifolia*), loquat (*Eriobotrya japonica*), Bowman's root (*Gillenia trifoliata*), almond (*Prunus dulcis*), peach (*Prunus persica*), strawberry (*Fragaria vesca*), and marijuana (*Cannabis sativa*)) (37, 95–101) were collected for multiple sequence alignments. Multiple sequence alignments were performed with LastZ v1.02 (102). A phylogenetic tree for the 4-fold conserved loci was then constructed based on the maximum likelihood method using IQ-TREE v2.1 software (103). Finally, the number of "rejected substitutions" (GERP score) was estimated using the 'gerpcol' module from the GERP++ software.

Since no sites GERP > 2 were found in the apple genome (see Supplementary Text above), we estimated genome-wide mutation load using the number of derived deleterious alleles identified in *Malus* accessions based on Sorting Intolerant From Tolerant 4G (SIFT 4G) scores (104). We constructed a SIFT 4G database specific for the apple (*Malus domestica*) genome using the SIFT4G_Create_Genomic_DB tool. The database was built based on the reference genome sequence (FASTA) and genome annotation (GFF3) files. To enhance the evolutionary conservation information, we incorporated the UniRef90 protein clustering database (105) as a source of homologous protein sequences during the database construction. This integration provided a comprehensive set of homologs for multiple sequence alignment, improving the accuracy of functional impact predictions. Following database construction, we employed the SIFT4G_Annotator tool to annotate genetic variants (in VCF format) with SIFT scores. We then polarized derived and ancestral alleles using *M. baccata* and *P. pyrifolia* as outgroups. The software est-sfs v2.04 (106) was used to estimate the ancestral alleles of *M. domestica, M. sieversii, M. orientalis,* and *M. sylvestris* individuals. First, short read sequences from five *M. baccata* (outgroup 1) and five *P. pyrifolia* (outgroup 2) were mapped to the GDDH v1.1 apple reference genome (37). Alleles with a frequency greater than 0.8 in the outgroup were considered as putatively ancestral, as high frequency in these related species likely reflects the ancestral state. The Kimura two-parameter model (107) was then applied to estimate the probability of ancestral alleles, and only those with an ancestral site probability greater than 0.9 were considered. Candidate deleterious mutations were screened based on the SIFT4G results, with a threshold set at a "SIFT score < 0.05" of any nonsynonymous position (108). We observed a reference bias, where sites with the derived allele in the reference apple genome were more often categorized as tolerated *versus* sites where the reference was ancestral, similar to findings in human and soybean genomes (Fig. S40).

We adjusted the derived deleterious allele count at sites where the reference genome contained the derived allele using the probability estimation method described by Simons (109). The mutation load for each accession was quantified by counting the derived deleterious alleles.

To examine the relationship between gene flow or selection sweep between mutation load. The mutation load of each gene was quantified by the average of derived deleterious alleles per 1,000 bp CDS region in DomC or DomD populations, which standardizes for gene length and facilitates comparison among genes. The introgression level ($f_d$) was calculated for each gene in the four population groups (SiOr-DomC-SylE-Bacc, SiOr-DomC-SylW-Bacc, SiOr-DomD-SylE-Bacc, and SiOr-DomD-SylW-Bacc). The deleterious alleles distribution per gene was fitted to a negative binomial distribution, the coefficients with $f_d$ were estimated by the *glm.nb* function from the MASS package (110). Genes from various selection regions, including soft or hard sweeps and adaptive introgression or non-introgression, were extracted, while the remaining genes function used as controls. The statistical significance of the difference between selection genes and control genes was assessed with a Wilcoxon test.

**Genome-wide association study (GWAS) of flowering time in cultivated apples**

To investigate phenotypic d simethingivergence between cider and dessert apples, we conducted a GWAS for flowering time using a subset of 30 cultivated accessions, each with six replicates in a randomized design measured across three common gardens over three years, planted in 2021. We analyzed genotype–phenotype associations using the GAPIT R package (v3.0) (111) implementing the BLINK method (112), with kinship (K) and population structure (Q) as covariates. SNPs were filtered for minor allele frequency (MAF ≥ 0.05) and missing rate < 20% prior to analysis. Significance thresholds were determined using both false discovery rate (FDR < 0.05) and Bonferroni correction (113). For significant SNPs, we examined average LD decay distance (10 Kb) nearby genes and transposable elements using the GDDH13 v1.1 genome gene annotation and TE annotations (37). To assess population-specific differentiation at these loci, we visualized genotype distributions with custom scripts generating barplots of allele frequencies by individual and genetic group (DomC, DomD, SiOr, SylE, SylW) based on fastSTRUCTURE assignments, excluding individuals with >20% missing data or admixture coefficients < 0.8. We assessed the distribution of flowering-associated alleles across selective sweep and introgressed regions in the DomC and DomD populations.

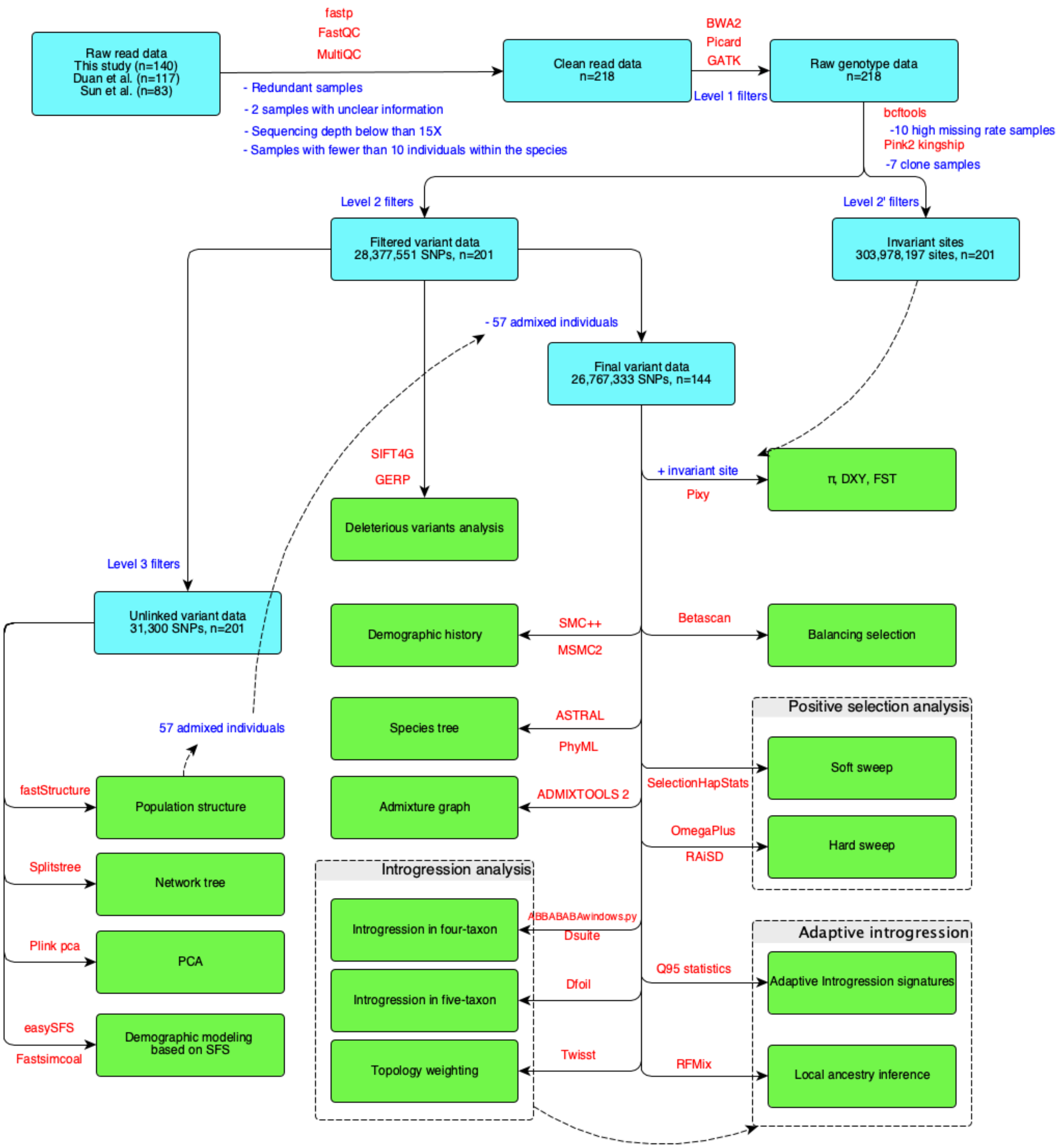


**Fig. S1. Bioinformatic workflow with the origin of data used, number of individuals, number of single-nucleotide polymorphisms (SNPs) used for each step, and method in this study.** Text in blue indicates data filtering and sample adjustment steps, and text in red indicates software used. All filtering parameters are provided in Table S2. Gene expression and GWAS analyses are not represented here as they were used anchor population genomics inferences and different number of samples and sequencing or phenotypes dataset were used.

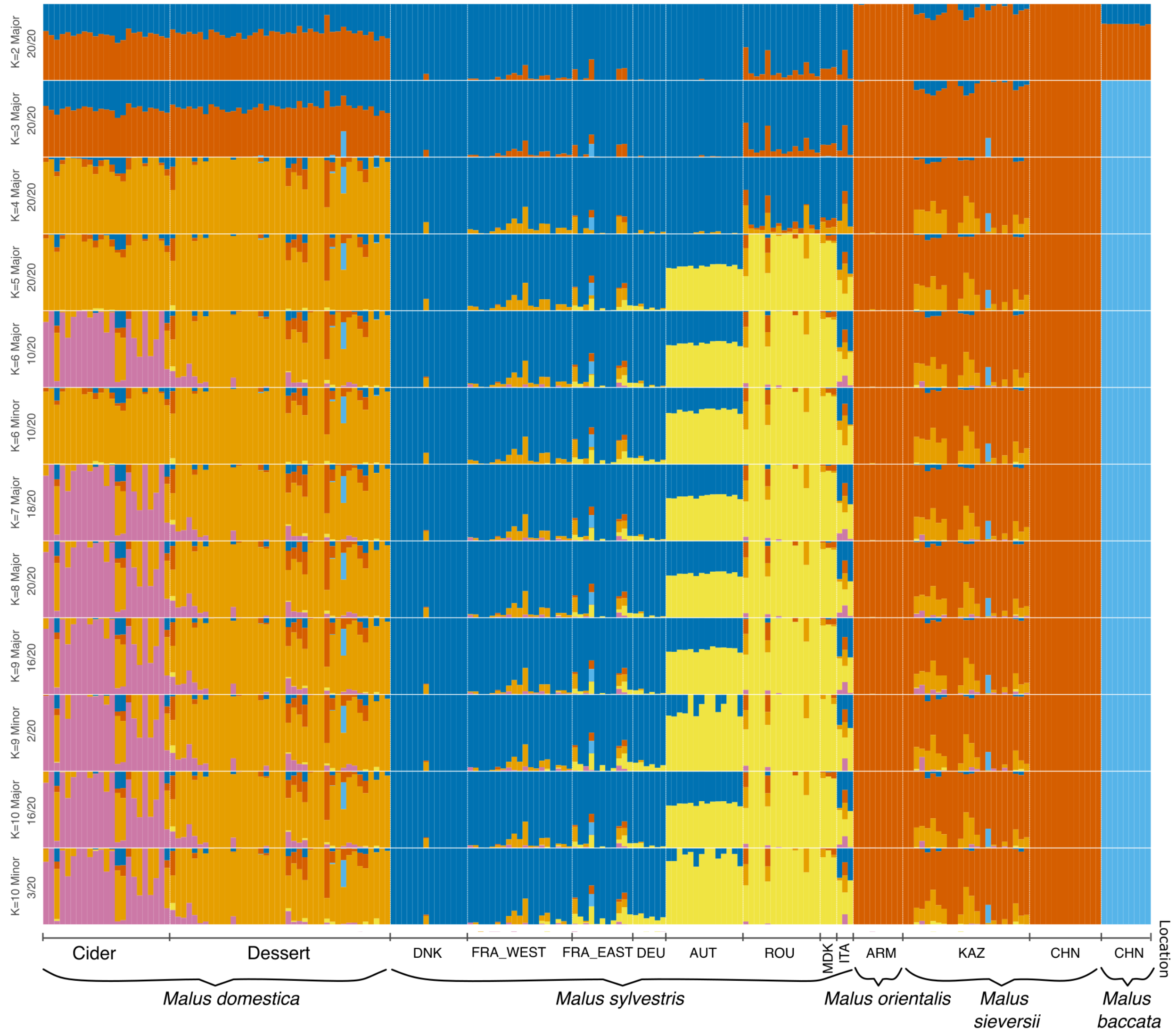


**Fig. S2. Population structure of the wild and cultivated apples inferred with fastStructure from *K* = 2 to *K* = 10 (N= 201, 31,300 unlinked synonymous SNPs).**

Each individual is represented by a vertical bar partitioned into *K* segments that represent the proportions of the ancestry of its genome in K clusters. Colors indicate the inferred ancestry from *K* ancestral genetic clusters). For each *K* values, minor and major modes and division of runs by mode are provided, when detected, with their respective percentage observed over 20 repetitions. *Malus domestica* cultivars were sorted according to their uses (cider or dessert). For the wild apples (*Malus sylvestris*, *M. orientalis*, *M. sieversii* and *M. baccata,* countries of sampling are provided (see details in Dataset S1). DNK: Denmark; FRA_WEST: Western France; FRA_EAST: Eastern France; DEU: Germany; AUT: Austria; ROU: Romania; MKD: Macedonia; ITA: Italy; ARM: Armenia; KAZ: Kazakhstan; CHN: China.

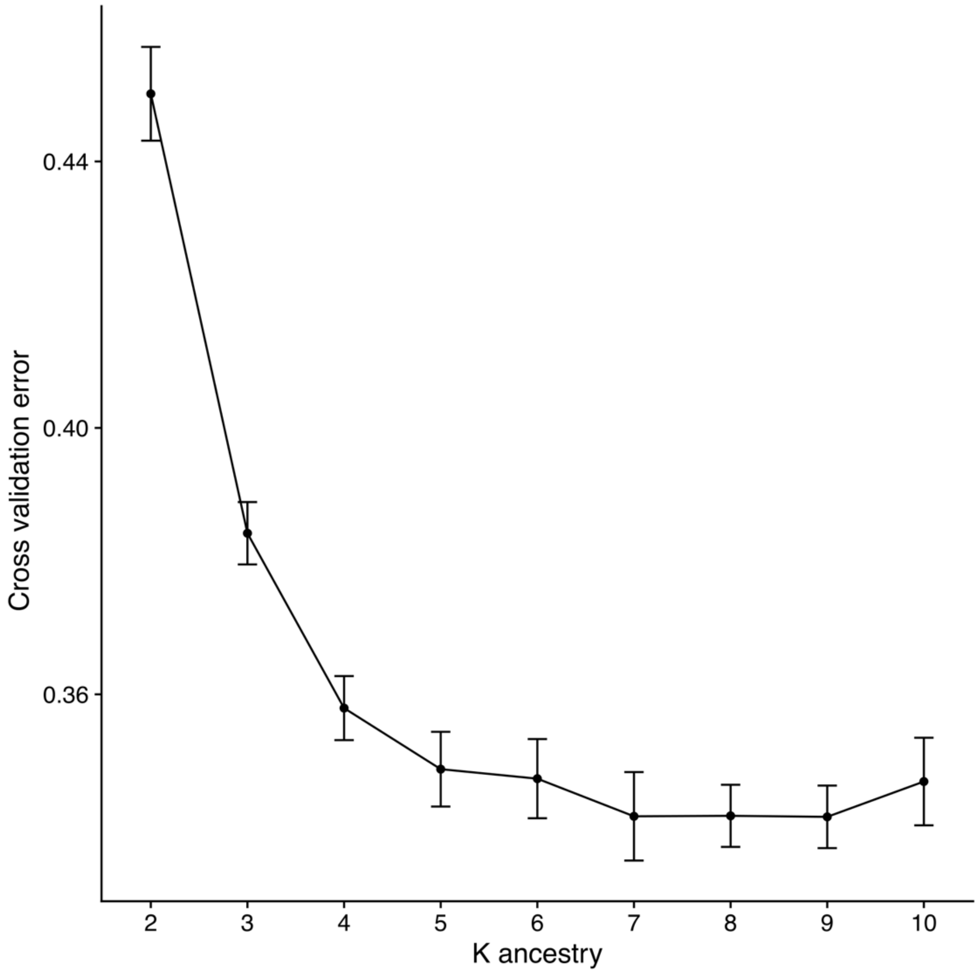


**Fig. S3. Cross-validation error for each *K* value obtained with fastSTRUCTURE analyses for the *Malus* accessions (*N*=201, 31,300 SNPs).**
The cross-validation error, designed to help identify the most relevant number of clusters, monotonically decreased up to K = 7, and no further substructures were observed at *K* > 6 (Fig. S2).

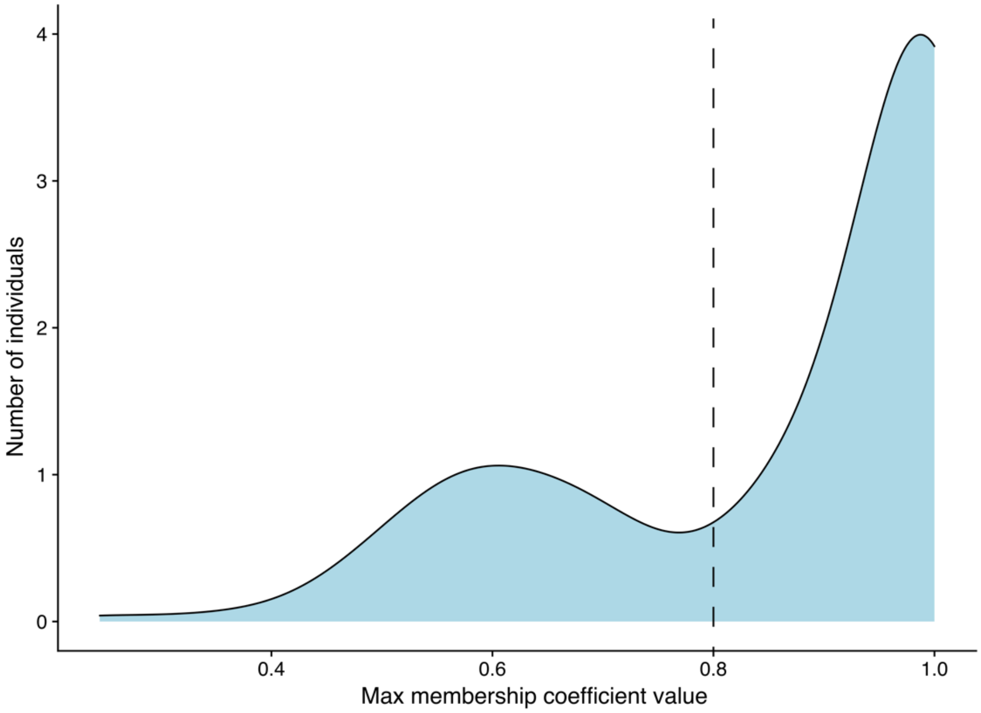


**Fig. S4. Distribution of the max membership coefficient inferred with fastSTRUCTURE for the 201 individuals of wild (*M. sylvestris*, *M. orientalis*, *M. sieversii*, *M. baccata*) and cultivated apples (*M. domestica*, N=68) at K=6.**

Individuals with a membership coefficient ≥ 0.8 were considered as admixed genotypes. The vertical line at 0.8 represents the threshold to assign an individual to a given population (i.e., genetic cluster excluding admixed individuals with a membership coefficient < 0.8).

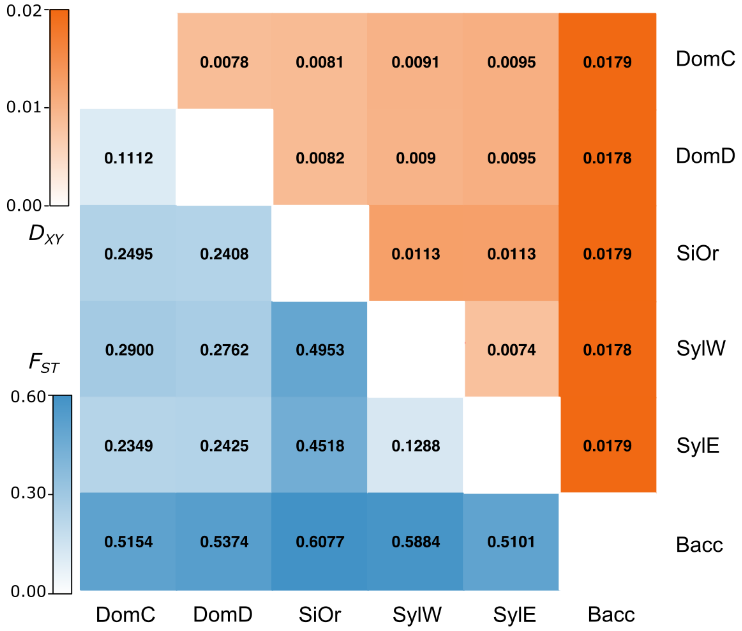


**Fig. S5. Heatmap of pairwise $F_{ST}$ and $D_{XY}$ estimates among wild and cultivated apple populations.**
DomC, Cider *M. domestica*; DomD, Dessert *M. domestica*; SiOr, *M. sieversii* and *M. orientalis*; SylE, Eastern Europe *M. sylvestris*; SylW, Western Europe *M. sylvestris*; Bacc, *M. baccata.*

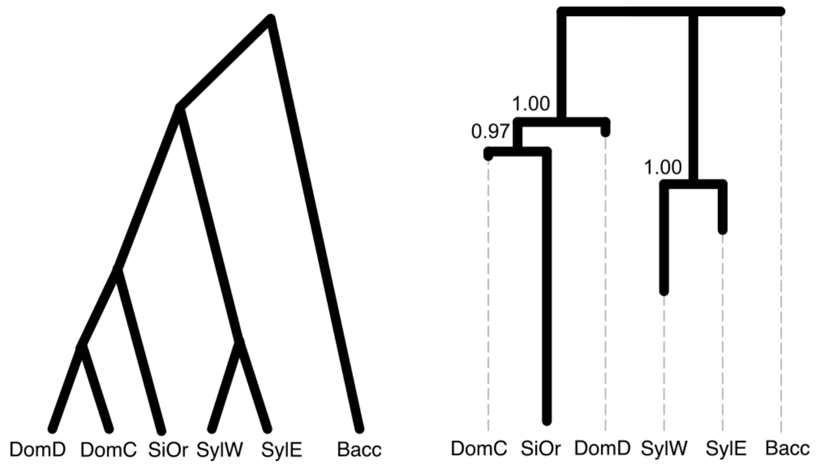


**Fig. S6. The most common species-tree topology (left) and the coalescent species-tree (right) among wild and cultivated apple inferred by ASTRAL.**
DomC, Cider *M. domestica*; DomD, Dessert *M. domestica*; SiOr, *M. sieversii* and *M. orientalis*; SylE, Eastern Europe *M. sylvestris*; SylW, Western Europe *M. sylvestris*; Bacc, *M. baccata.*

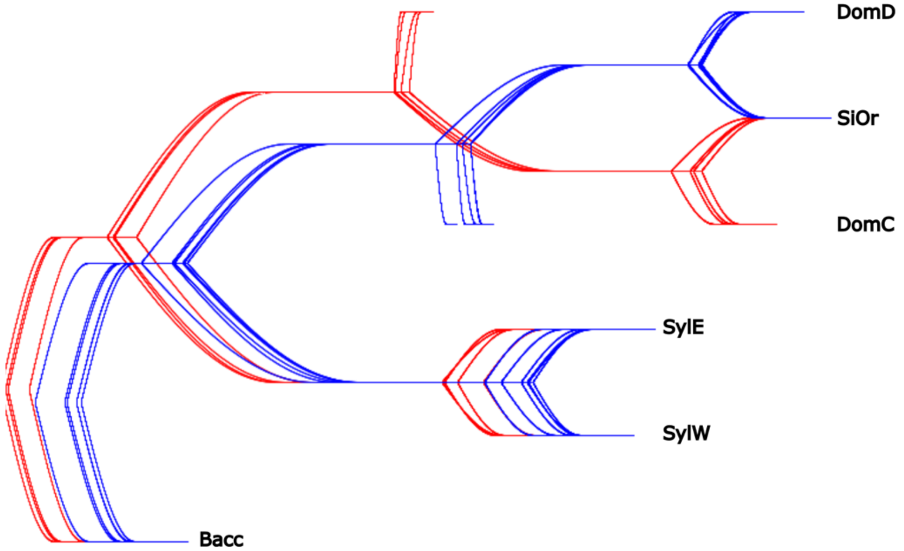


**Fig. S7. DensiTree visualization of 10 coalescent species trees of wild and cultivated apples inferred by ASTRAL, each based on a random set of nine individuals per populations.**
Blue and red lines indicate two alternative species-tree topologies. DomC, Cider *M. domestica*; DomD, Dessert *M. domestica*; SiOr, *M. sieversii* and *M. orientalis*; SylE, Eastern Europe *M. sylvestris*; SylW, Western Europe *M. sylvestris*; Bacc, *M. baccata.*

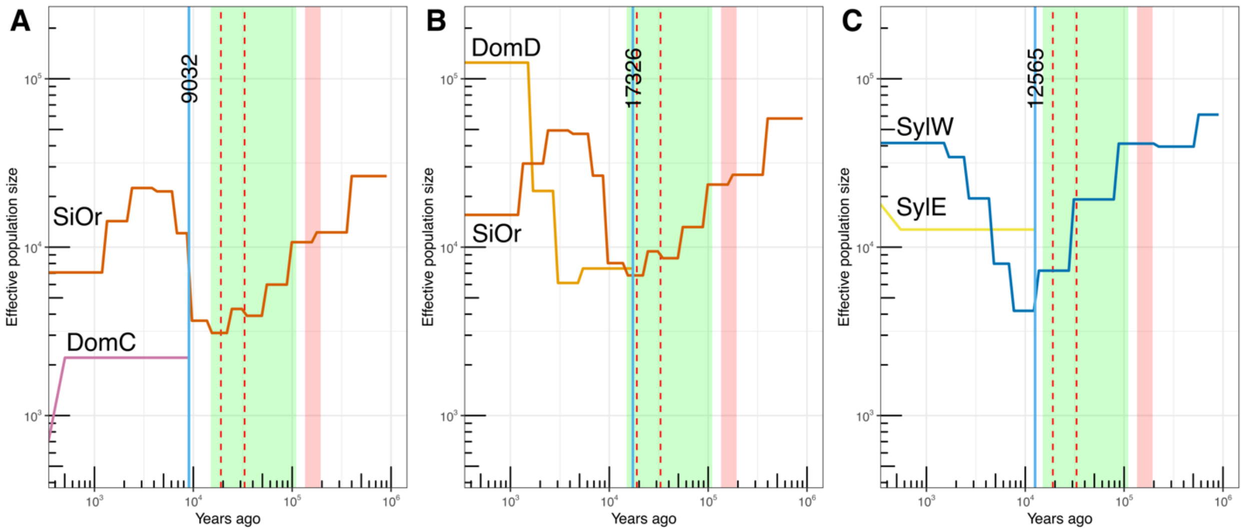


**Fig. S8. Estimated split times among wild and cultivated apple populations inferred with SMC++.**

Red highlights: Penultimate glacial period; green highlights: last glacial period, LGP; between dashed line: last glacial period, LGM; Blue vertical line and the numbers: estimate split times between two populations. DomC, Cider *M. domestica*; DomD, Dessert *M. domestica*; SiOr, *M. sieversii* and *M. orientalis*; SylE, Eastern Europe *M. sylvestris*; SylW, Western Europe *M. sylvestris*; Bacc, *M. baccata.*

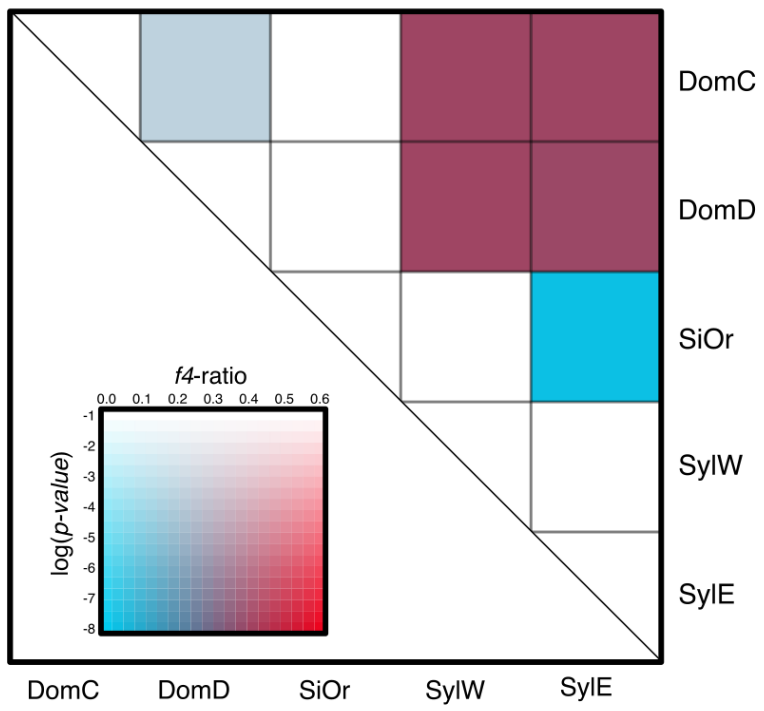


**Fig. S9. *f*$_4$-ratio statistic estimated with Dsuite among the five wild and cultivated apple populations using Bacc as the outgroup.**
DomC, Cider *M. domestica*; DomD, Dessert *M. domestica*; SiOr, *M. sieversii* and *M. orientalis*; SylE, Eastern Europe *M. sylvestris*; SylW, Western Europe *M. sylvestris*; Bacc, *M. baccata.*

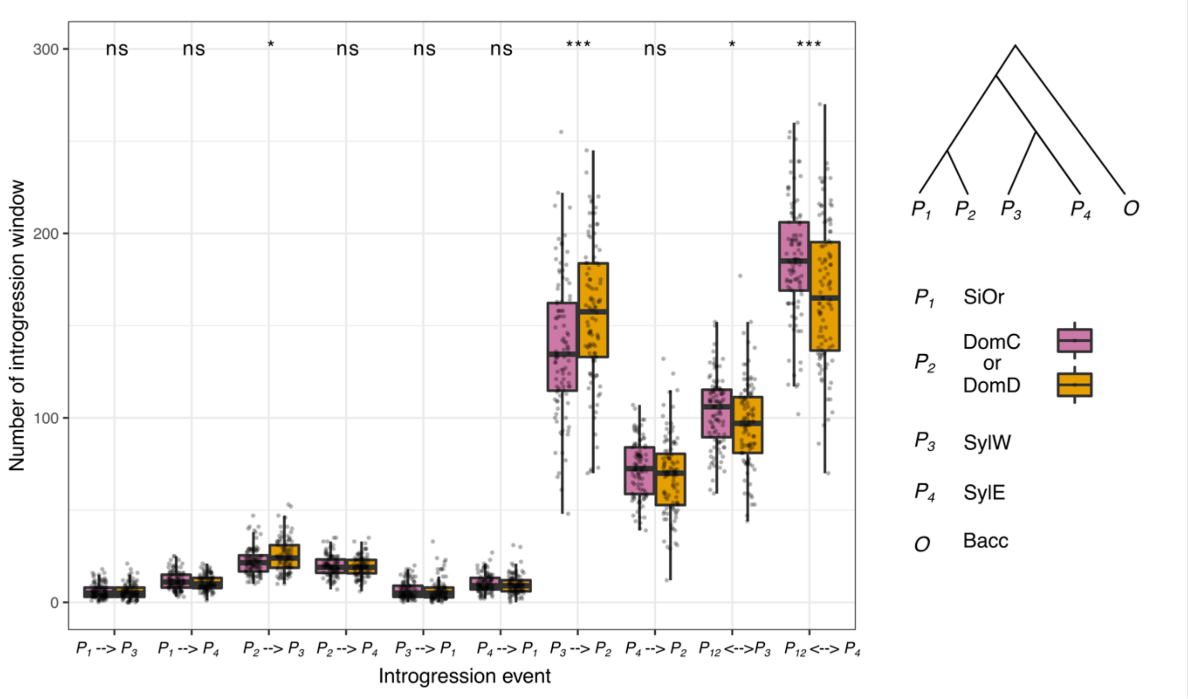


**Fig. S10. Results of the $D_{\mathrm{FOIL}}$ analyses of 100 independent runs (when $P_2$ represents DomC or DomD.**

The topologies used in the analysis are shown on the right (*M. sylvestris* divergence time is earlier than *M. domestica*). Populations $P_1$ to $P_4$ and $O$ (outgroup) are indicated in the legend. $P_{12}$ represents the ancestral population of $P_1$ and $P_2$. We analyzed a total of 6,125 100 Kb windows per run in which $D_{\mathrm{FOIL}}$ detected gene flow between lineages as indicated on the x-axis. The unidirectional arrows on the x-axis depict the direction of gene flow, while bidirectional arrows indicate that the direction of gene flow cannot be distinguished. The box limit indicates the 25th and 75th percentiles of the window numbers, and the center line shows the mean, respectively. The dots indicate the number of windows with a signal of gene flow from independent runs. DomC, Cider *M. domestica*; DomD, Dessert *M. domestica*; SiOr, *M. sieversii* and *M. orientalis*; SylE, Eastern Europe *M. sylvestris*; SylW, Western Europe *M. sylvestris*; Bacc, *M. baccata.*

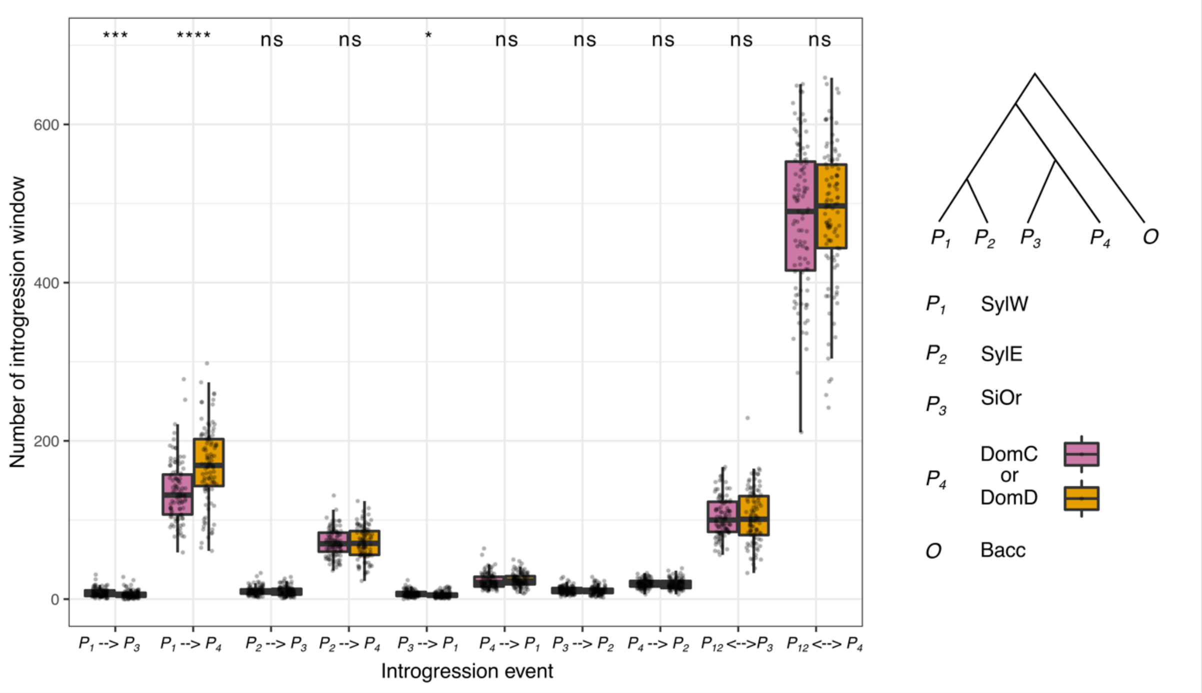


**Fig. S11. Results of the *D*FOIL analyses of 100 independent runs (when *P*$_4$ represents DomC or DomD).**

The topologies used in the analysis are shown on the right (*M. domestica* divergence time is earlier than *M. sylvestris* ). Populations *P*$_1$ to *P*$_4$ and *O* (outgroup) are indicated in the legend. *P*$_{12}$ represents the ancestral population of *P*$_1$ and *P*$_2$. We analyzed a total of 6,125 100 Kb windows per run in which *D*FOIL detected gene flow between lineages as indicated on the x-axis. The unidirectional arrows on the x-axis depict the direction of gene flow, while bidirectional arrows indicate that the direction of gene flow cannot be distinguished. The box limit indicates the 25th and 75th percentiles of the window numbers, and the center line shows the mean, respectively. The dots indicate the number of windows with a signal of gene flow from independent runs. DomC, Cider *M. domestica*; DomD, Dessert *M. domestica*; SiOr, *M. sieversii* and *M. orientalis*; SylE, Eastern Europe *M. sylvestris*; SylW, Western Europe *M. sylvestris*; Bacc, *M. baccata.*

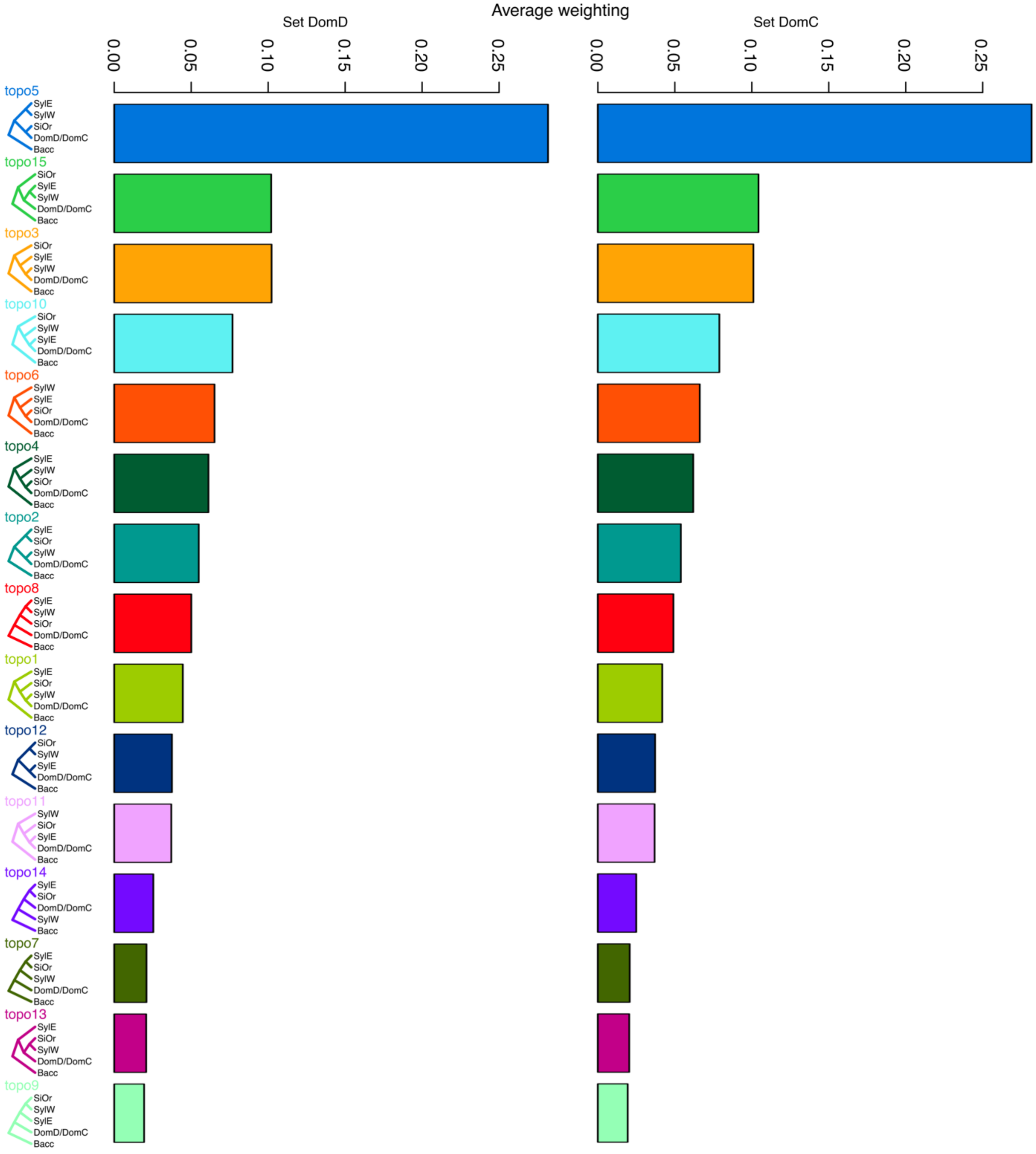


**Fig. S12. Topology weighting averages across the genomes of apple populations.**
Left are the 15 possible topologies from two sets of five-taxa (Set DomD and DomC). Right, the average weightings (%) for each of the 15 topologies are included in each bar. DomC, Cider *M. domestica*; DomD, Dessert *M. domestica*; SiOr, *M. sieversii* and *M. orientalis*; SylE, Eastern Europe *M. sylvestris*; SylW, Western Europe *M. sylvestris*.

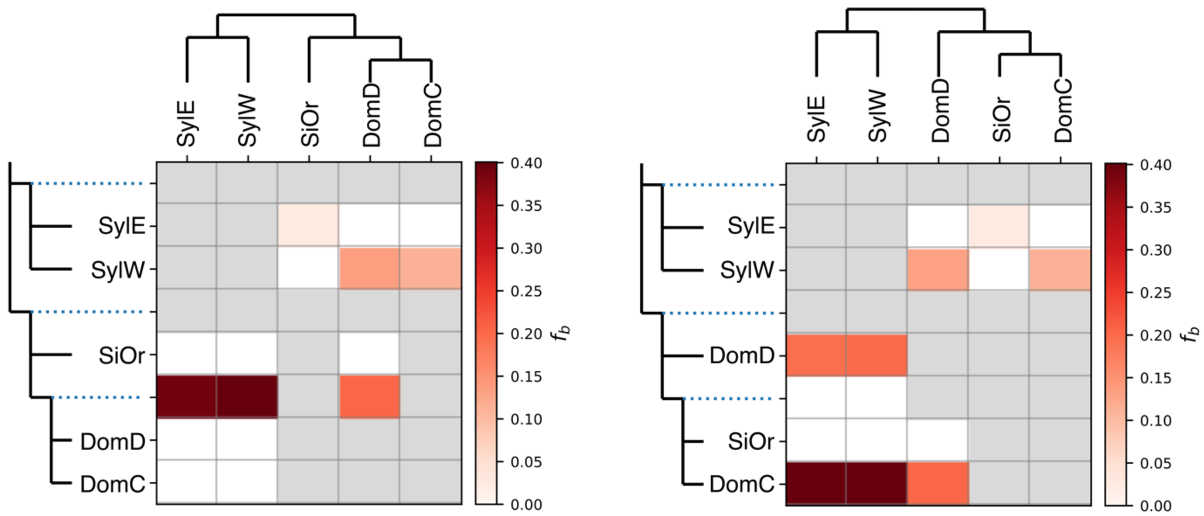


**Fig. S13. The branch-specific statistic *f*-branch identifies excess sharing of derived alleles between the branch of the tree on the *y*-axis and the population on the *x*-axis**
The left is the *f*-branch result based on the most common species-tree topology, and the right is the *f*-branch result based on the coalescent species-tree inferred by ASTRAL. The rows depict the tree topology nodes, while the columns represent the tips. Each cell displays the *f*-branch statistic value between a tree node (rows) and each tip (columns). Empty grey cells indicate comparisons that could not be made within the dataset.

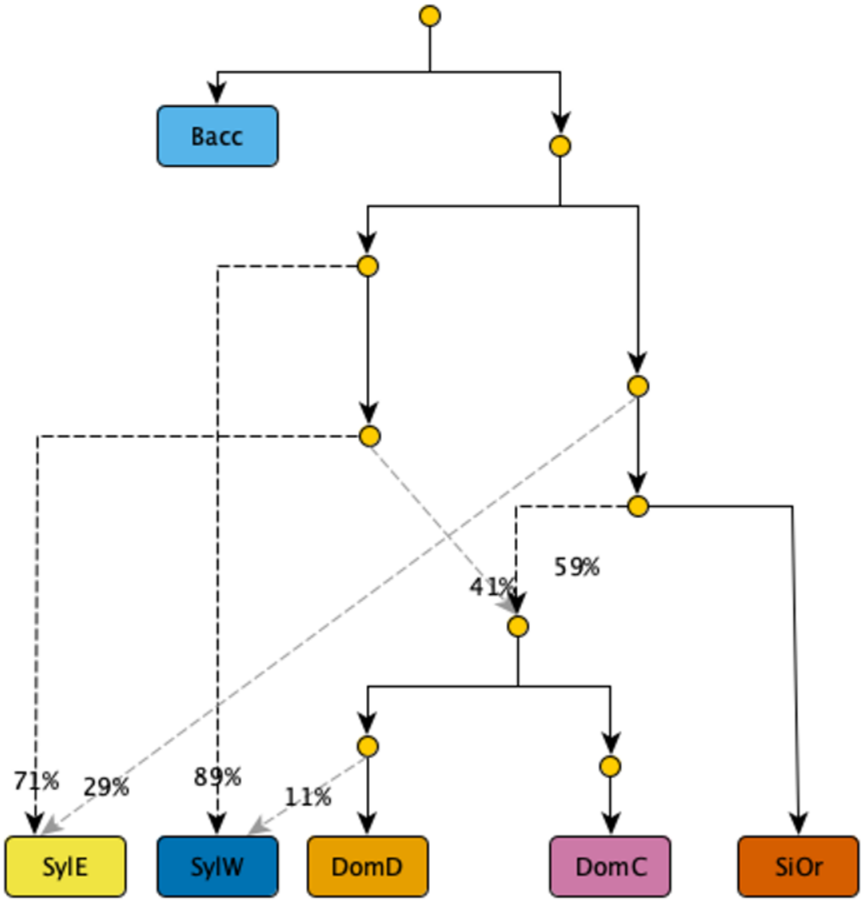


**Fig. S14. Best-fitting admixture graph based on the *qpGraph* models.**
Dashed lines represent admixture edges, and the admixture proportions are indicated next to the dashed line arrows. DomC, Cider *M. domestica*; DomD, Dessert *M. domestica*; SiOr, *M. sieversii* and *M. orientalis*; SylE, Eastern Europe *M. sylvestris*; SylW, Western Europe *M. sylvestris*; Bacc, *M. baccata.*

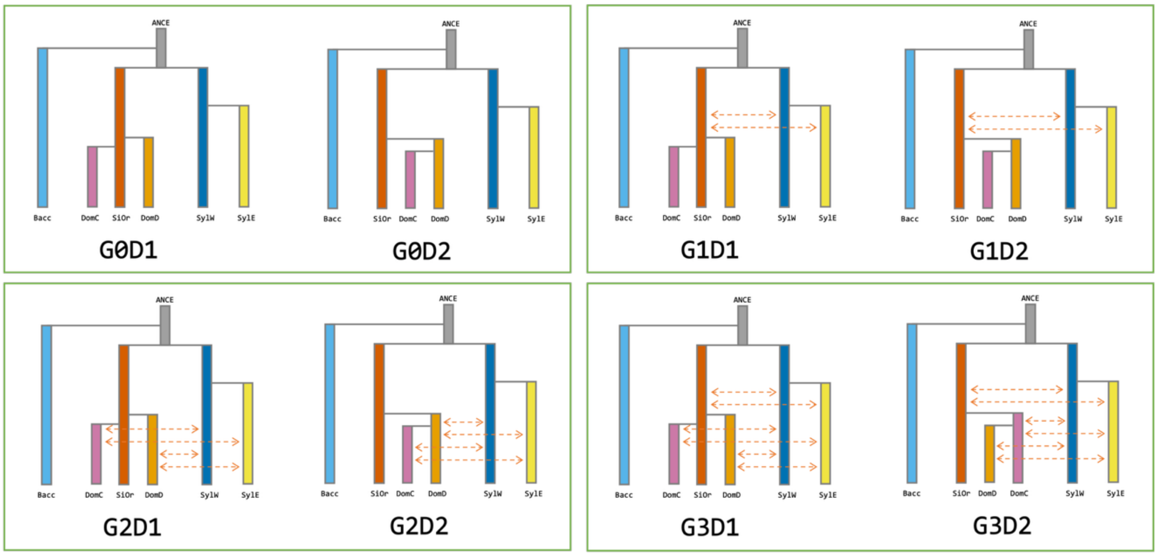


**Fig. S15. Apple demographic history scenarios simulated in fastsimcoal.**
G0 represents no gene flow within populations. G1 involves gene flow between the ancestor of DomC/DomD and SiOr and SylW/SylE populations. G2 represents gene flow between DomC/DomD and SylW/SylE populations, while G3 combines scenarios G1 and G2. Divergence scenarios include D1, where DomC and DomD both diverge from SiOr, with DomC diverging more recently and SylE diverging from SylW; D2, where DomD diverges from SiOr, followed by DomC diverging from DomD, and SylE diverging from SylW. Population identifiers are as follows: DomC refers to Cider *M. domestica*, DomD to Dessert *M. domestica*, SiOr to *M. sieversii* and *M. orientalis*, SylE to Eastern Europe *M. sylvestris*, SylW to Western Europe *M. sylvestris*, and Bacc to *M. baccata*.

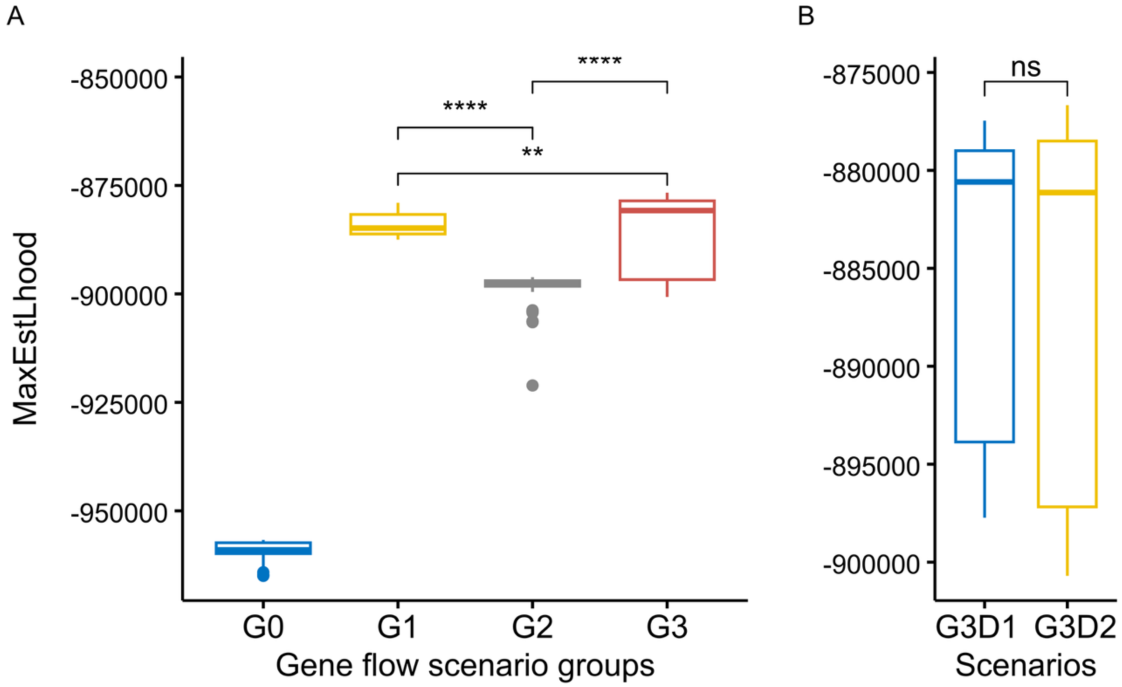


**Fig. S16. Maximum likelihood distributions for each gene flow scenario group (A) and two G3 scenarios (B) simulated with fastsimcoal2.**

Distributions were derived from 50 expected site frequency spectra (SFS), with each approximation based on 100,000 coalescent simulations. G0, no gene flow within populations; G1, gene flow between the ancestor of DomC/DomD and SiOr and SylW/SylE populations; G2, gene flow between DomC/DomD and SylW/SylE populations; G3, G1 + G3. Four stars: Wilcoxon test $P < 0.0001$.

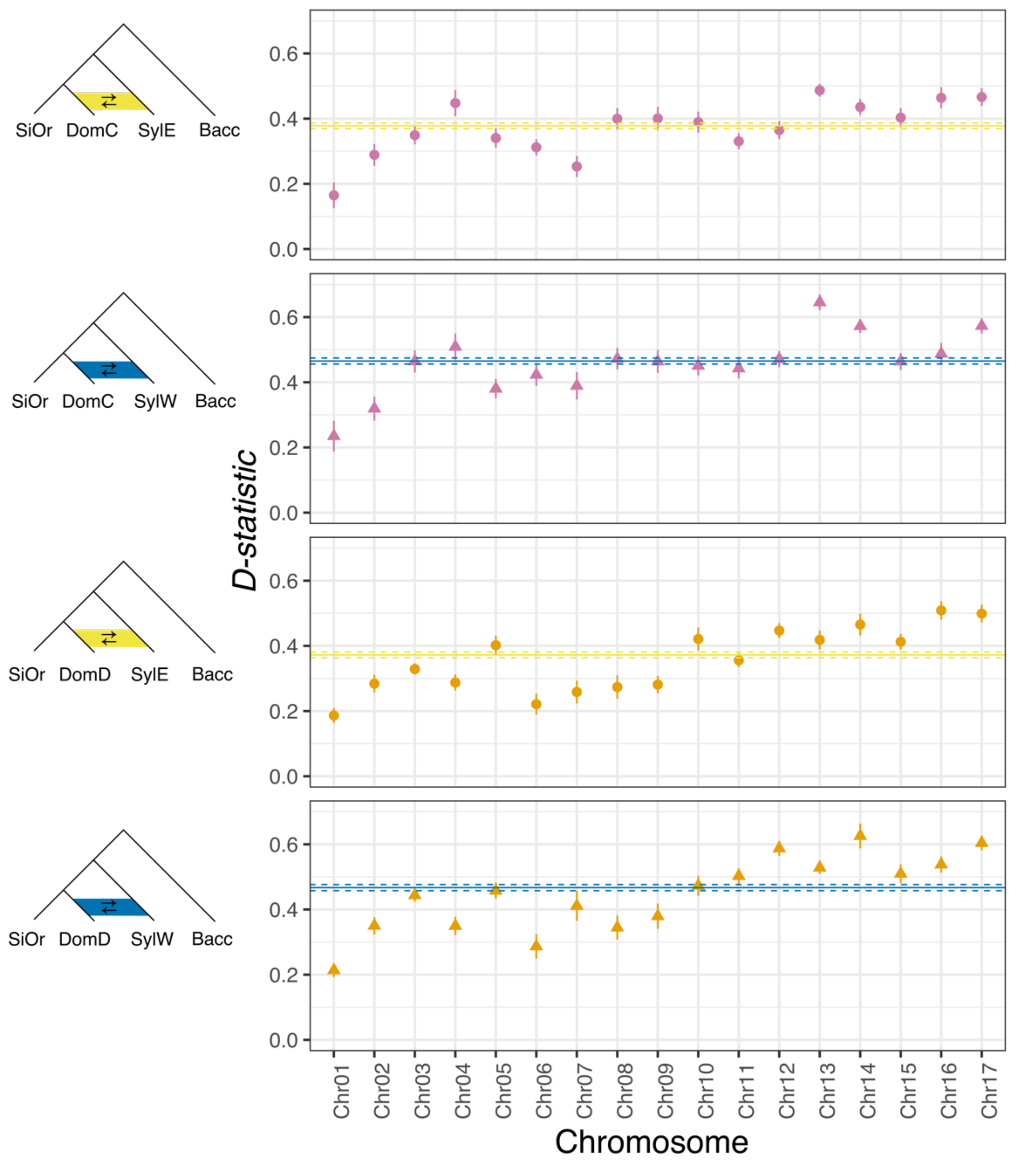


**Fig. S17. *D*-statistic for P1 (SiOr), P2 (DomC or DomD), P3 (SylE or SylW), and outgroup (*Bacc*) estimated per chromosome.**

The positive *D* value indicates the introgression of P2 and P3. The line crossed by the chromosomes represents the *D* value at the genome level. The topologies used in the analysis are shown on the left. DomC, Cider *M. domestica*; DomD, Dessert *M. domestica*; SiOr, *M. sieversii* and *M. orientalis*; SylE, Eastern Europe *M. sylvestris*; SylW, Western Europe *M. sylvestris*; Bacc, *M. baccata.*

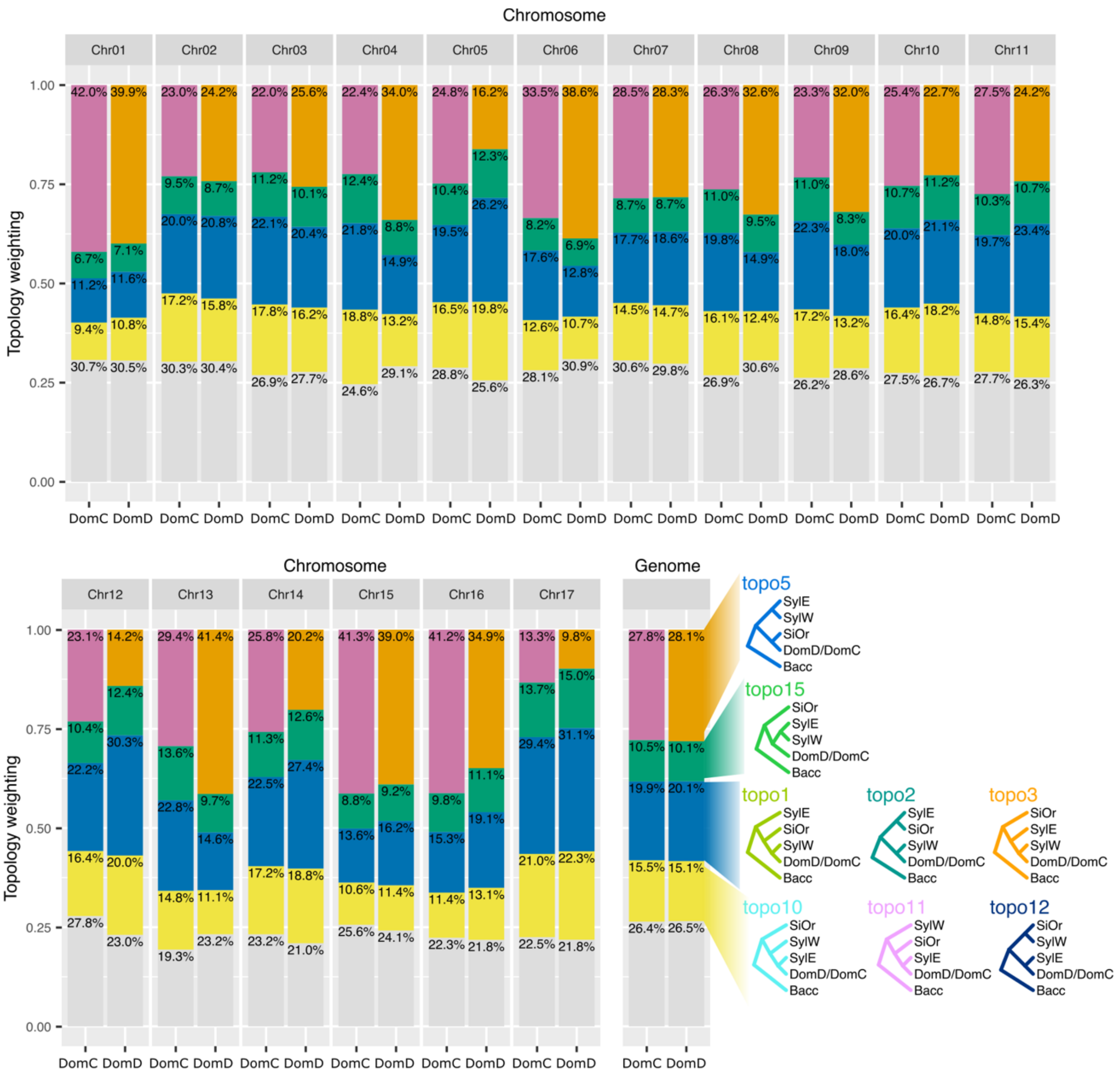


**Fig. S18. Phylogenies inferred from regions on the 17 chromosomes with Twisst.**
NJ-rooted phylogenies were inferred from n = 264,168 trees, 100 SNPs genomic windows for Set DomC and Set DomD. Colors correspond to the topology groups in the bottom left. Grey bars represent the other topologies. The four phylogenies groups are shown. Pink or Orange species tree; Green, DomC, or DomD are introgressed by the ancestor of SylE and SylW. Blue, DomC or DomD are introgressed by the SylW; Yellow, DomC or DomD are introgressed by the SylE. DomC, Cider *M. domestica*; DomD, Dessert *M. domestica*; SiOr, *M. sieversii* and *M. orientalis*; SylE, Eastern Europe *M. sylvestris*; SylW, Western Europe *M. sylvestris*; Bacc *M. baccata*.

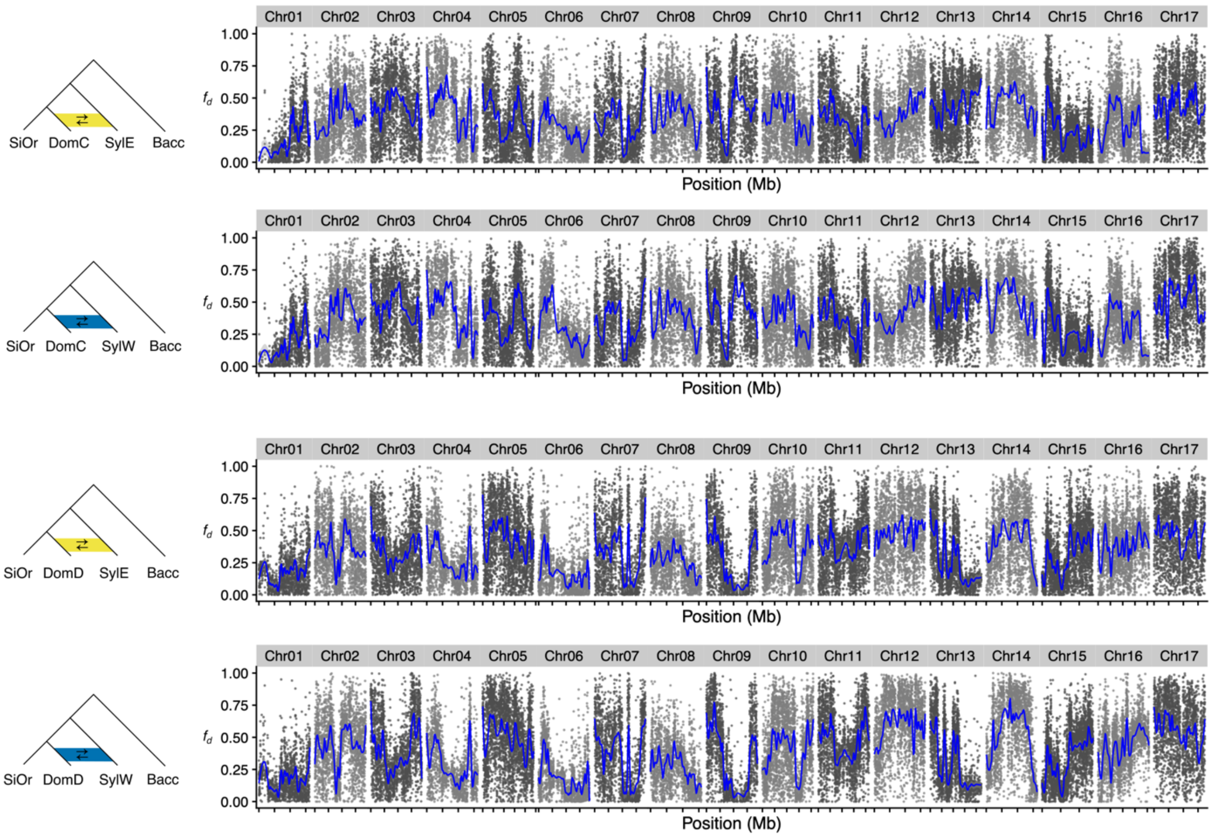


**Fig. S19. Genome-wide distribution of $f_d$ values calculated for 10 Kb windows across the P1 (SiOr), P2 (DomC or DomD), P3 (SylE or SylW), and outgroup (Bacc).**

The topologies used in the analysis are shown on the left. The blue line represents a loess fit with a 95% confidence interval. DomC, Cider *M. domestica*; DomD, Dessert *M. domestica*; SiOr, *M. sieversii* and *M. orientalis*; SylE, Eastern Europe *M. sylvestris*; SylW, Western Europe *M. sylvestris*; Bacc, *M. baccata*.

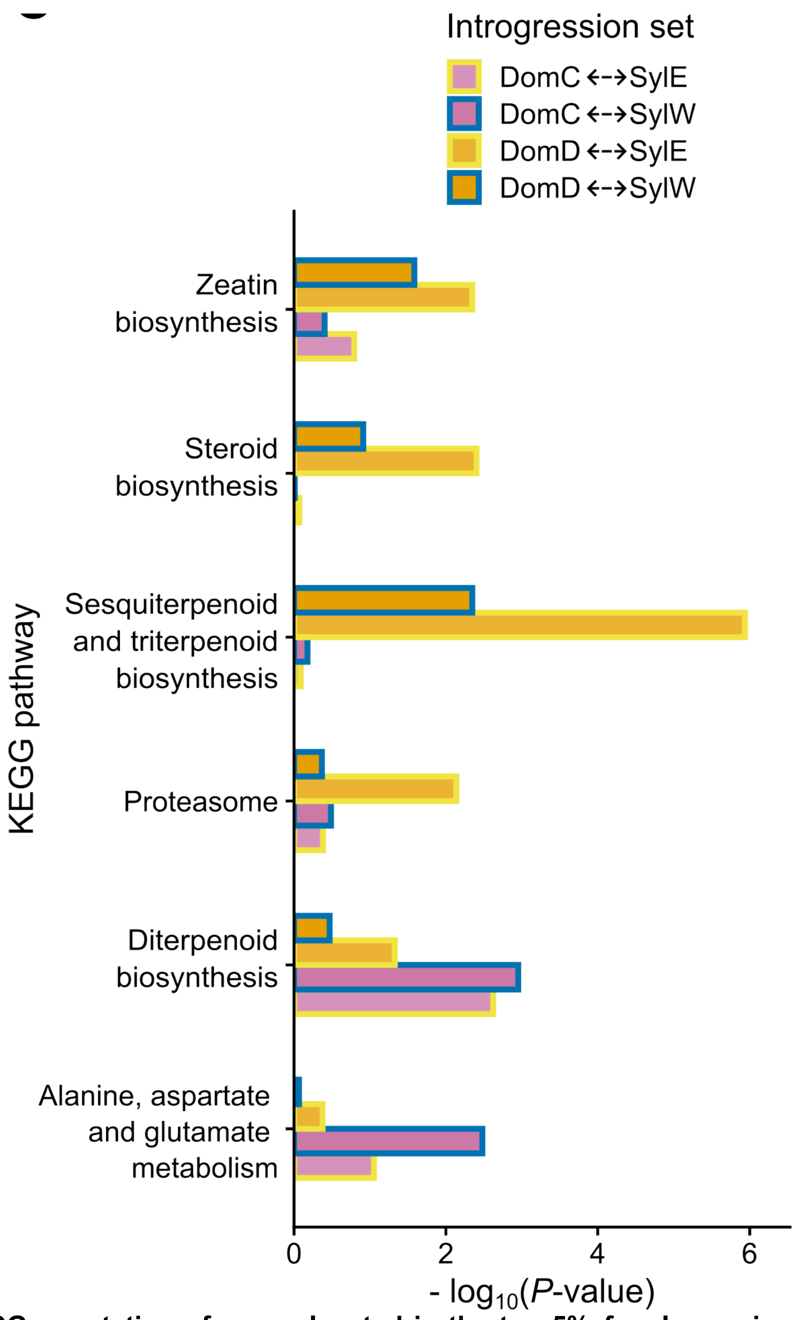


**Fig. S20. KEGG annotation of genes located in the top 5% $f_d$ value regions of cider and dessert apples.**

Frames and fill colors represent different introgressed population sets. The x-axis displays *p*-values on a $\log_{10}$ scale. DomC, Cider *M. domestica*; DomD, Dessert *M. domestica*; SiOr, *M. sieversii* and *M. orientalis*; SylE, Eastern Europe *M. sylvestris*; SylW, Western Europe *M. sylvestris*.

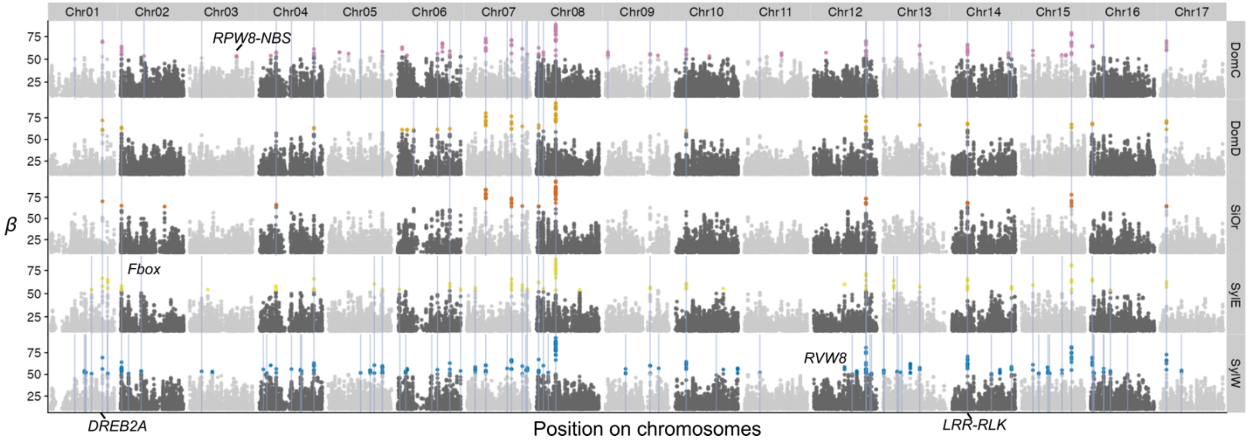


**Fig. S21. Genomic landscape of $\beta$ values, highlighting regions under balancing selection in wild and cultivated apple populations.**

Vertical lines indicate genes with shared balancing selection signals across populations. Population identifiers are as follows: DomC refers to Cider *M. domestica*, DomD to Dessert *M. domestica*, SiOr to *M. sieversii* and *M. orientalis*, SylE to Eastern Europe *M. sylvestris*, and SylW to Western Europe *M. sylvestris*. Gene ID are documented when available in the litterature.

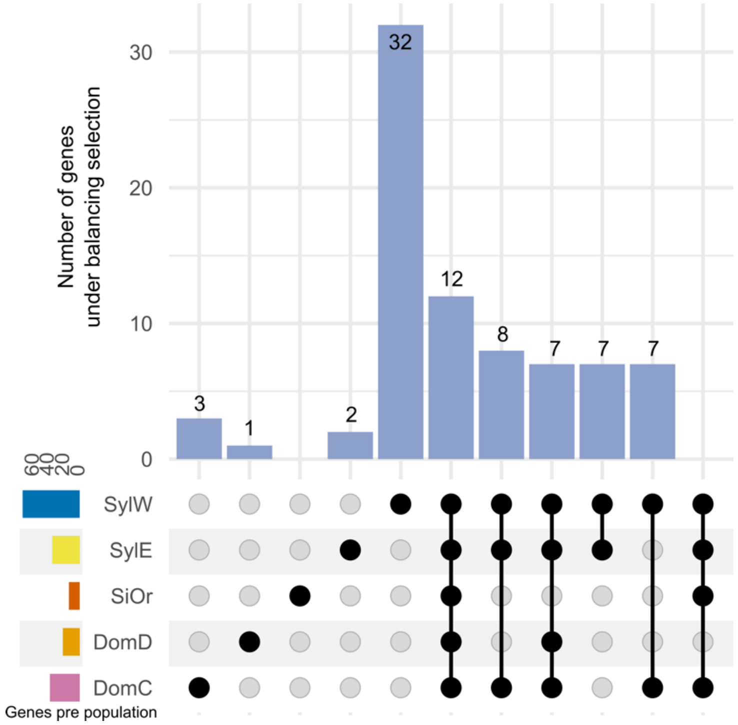

**Fig. S22. Upset plot of genes under balancing selection among five apple populations.** The horizontal bars (bottom left) display the total number of genes under balancing selection for each population. The matrix (bottom right) shows the populations under consideration as black circles. When an intersection between two or more populations is displayed, these populations are connected by a solid black line. The bar graph (top) represents the number of genes under balancing selection in a population or the intersections between two or more populations. Population identifiers are as follows: DomC refers to Cider *M. domestica*, DomD to Dessert *M. domestica*, SiOr to *M. sieversii* and *M. orientalis*, SylE to Eastern Europe *M. sylvestris*, and SylW to Western Europe *M. sylvestris*.

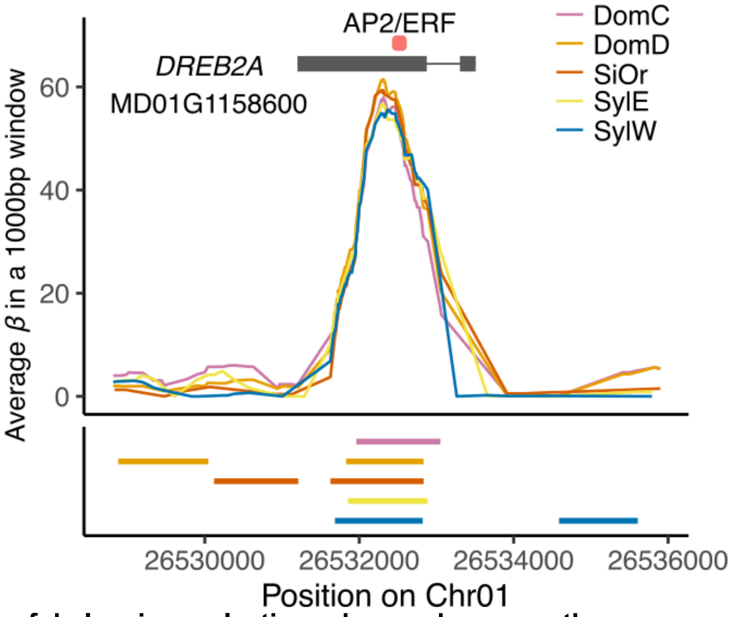


**Fig. S23. Signal of balancing selection observed across the sequence encoding the AP2/ERF domain of the *DREB2A* gene in five apple populations.**

Top, gene model, and protein domains; Bottom, linkage disequilibrium (LD) blocks spanning each population. Population identifiers are as follows: DomC refers to Cider *M. domestica*, DomD to Dessert *M. domestica*, SiOr to *M. sieversii* and *M. orientalis*, SylE to Eastern Europe *M. sylvestris*, and SylW to Western Europe *M. sylvestris*.

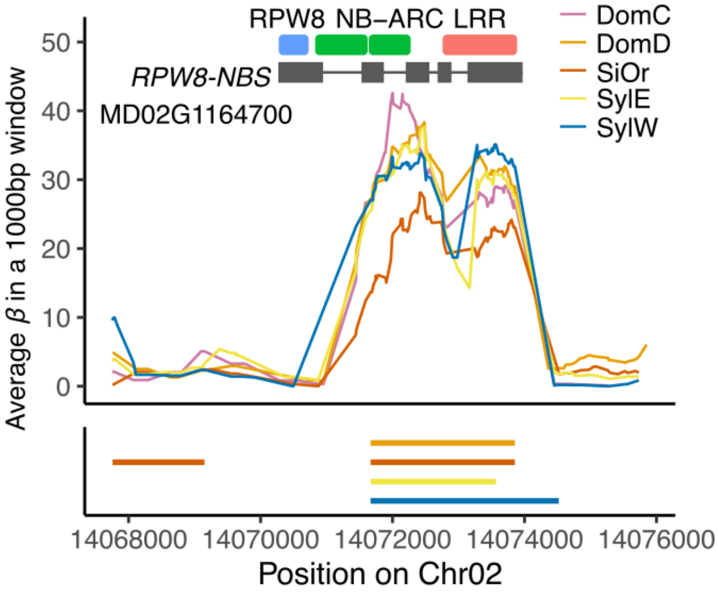


**Fig. S24. Signal of balancing selection observed across the sequence encoding the RPW8-NBS domain of the *RPW8-NBS* gene in five apple populations.**

Top, gene model, and protein domains; Bottom, linkage disequilibrium (LD) blocks spanning each population. Population identifiers are as follows: DomC refers to Cider *M. domestica*, DomD to Dessert *M. domestica*, SiOr to *M. sieversii* and *M. orientalis*, SylE to Eastern Europe *M. sylvestris*, and SylW to Western Europe *M. sylvestris*.

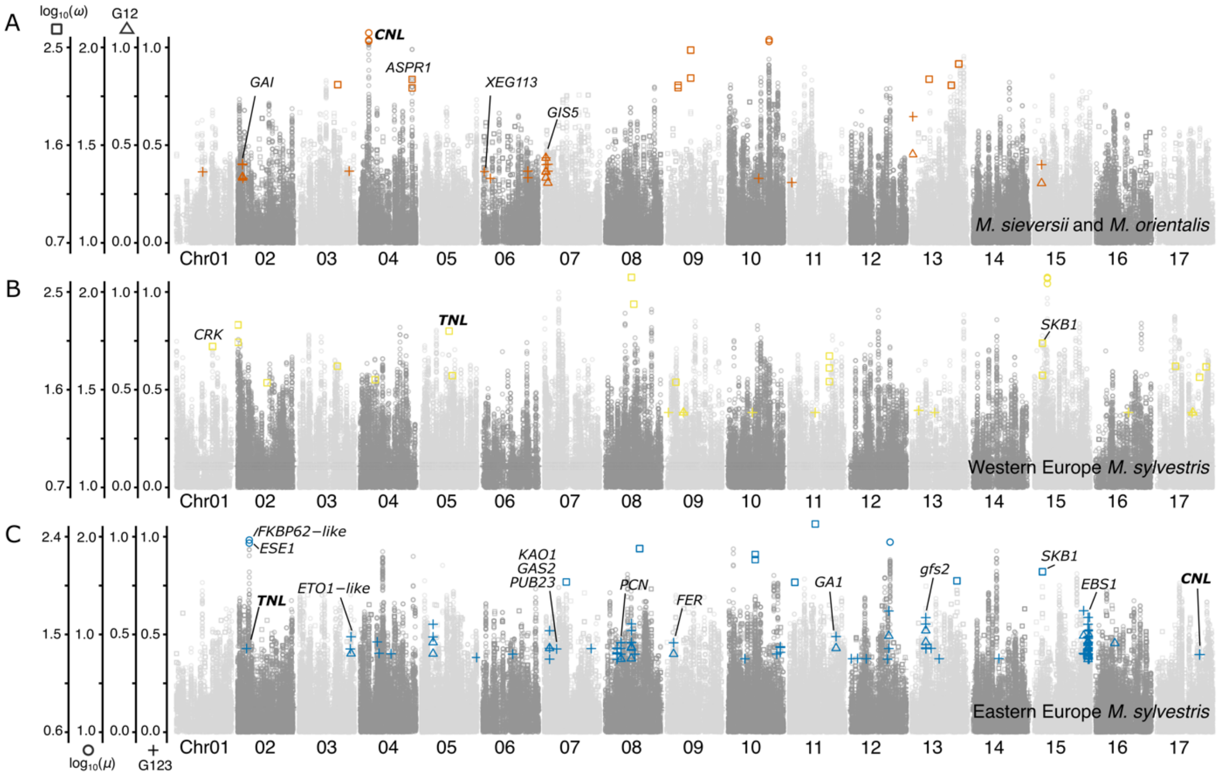


**Fig. S25. Positive selection scores across the genome of three wild apple populations.**
(A) *M. sieversii* and *M. orientalis* population (SiOr) (B) Western Europe *M. sylvestris* (C) Eastern Europe *M. sylvestris.* Different shapes and y-axis represent different types of selection tests. They are the Log$_{10}$($\omega$), Log$_{10}$($\mu$), G12, and G123 values from left to right. Significant selective sweep signals are represented in colors. Bold text represents disease resistance genes (R genes) from different families, including TIR-domain-containing (TNL) and CC-domain-containing (CNL) subfamilies.

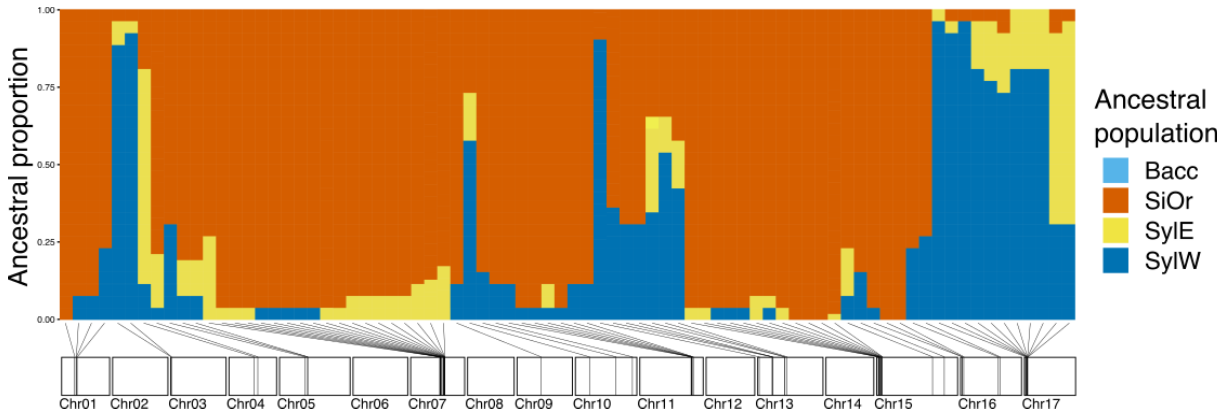


**Fig. S26. Ancestral population proportions across selective sweep regions in DomC (cider apple) estimated with RFMix.**
Colors represent ancestral populations. Chromosome diagrams below the plot are drawn to scale based on physical positions, with selective sweep regions positions along each chromosome. SiOr, *M. sieversii* and *M. orientalis*; SylE, Eastern European *M. sylvestris*; SylW, Western European *M. sylvestris*; Bacc to *M. baccata*.

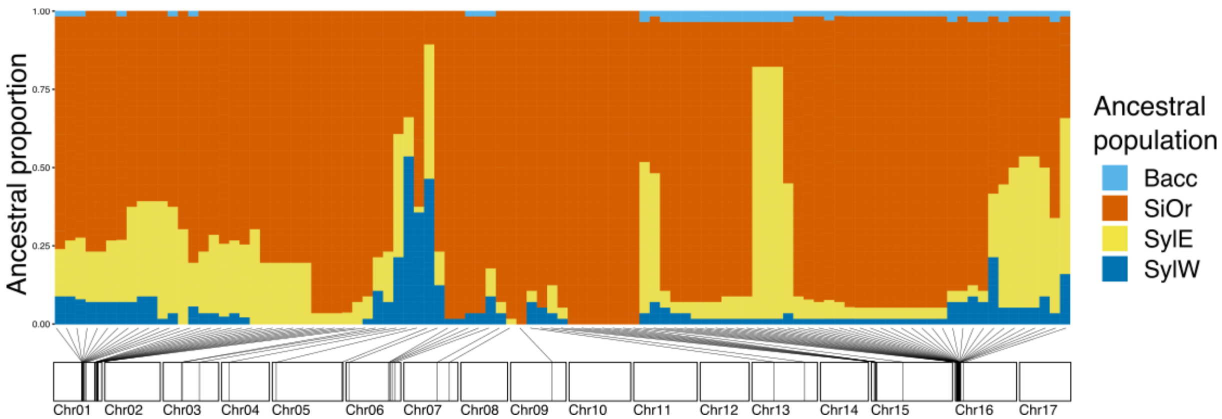


**Fig. S27. Ancestral population proportions across selective sweep regions in DomD (cider apple) estimated with RFMix.**

Colors represent ancestral populations. Chromosome diagrams below the plot are drawn to scale based on physical positions, with selective sweep regions positions along each chromosome. SiOr, *M. sieversii* and *M. orientalis*; SylE, Eastern European *M. sylvestris*; SylW, Western European *M. sylvestris*; Bacc to *M. baccata*.

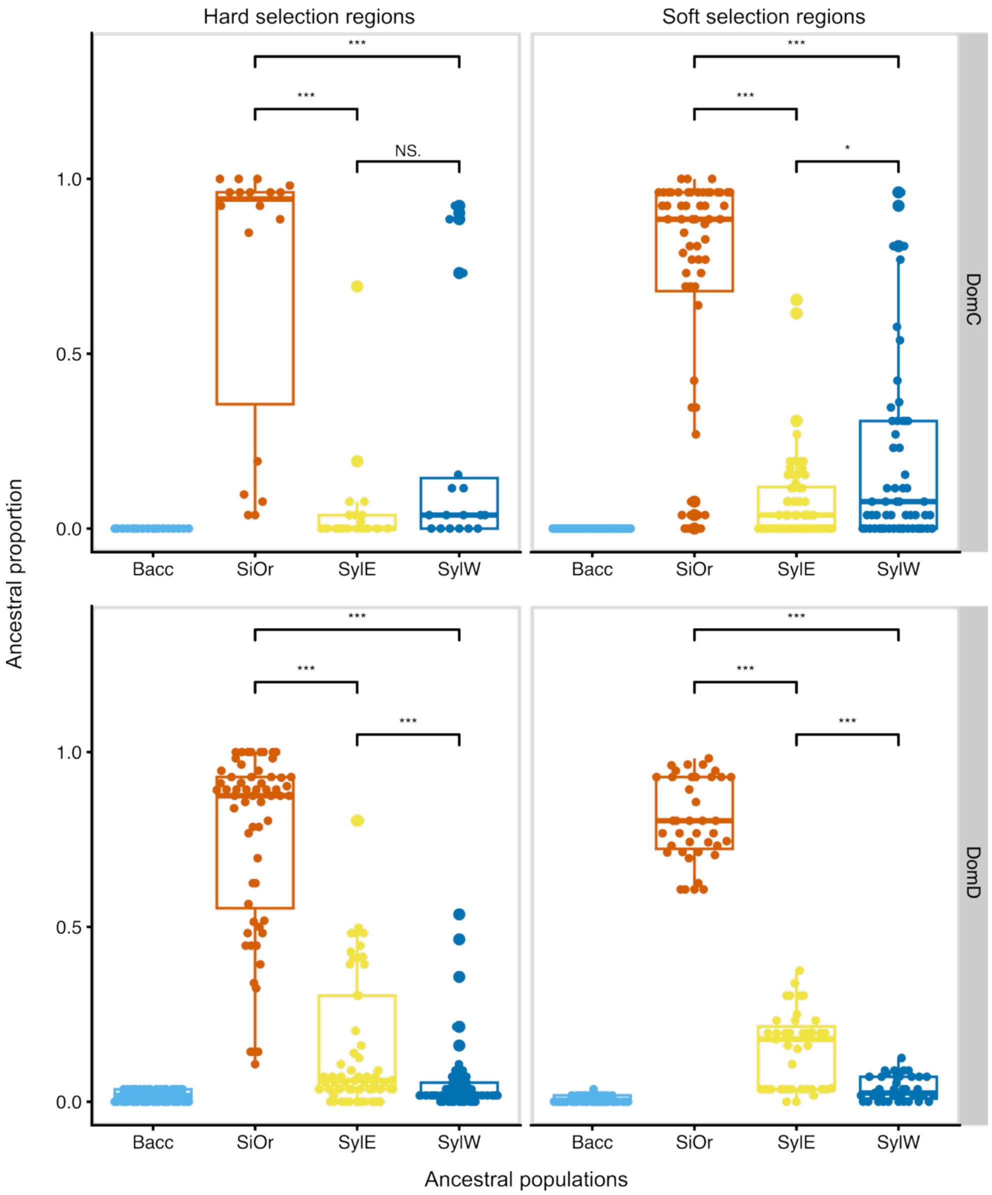


**Fig. S28. Boxplot of different wild ancestry population proportions under hard/soft sweep regions in DomC and DomD population estimated using RFMix.**

Colors represent ancestral populations. Boxplots show the proportions of different wild ancestry populations (Bacc, SiOr, SylE, SylW) within hard and soft sweep regions in DomC and DomD populations. Asterisks indicate significant differences between groups (*** $p < 0.001$). DomC, Cider *M. domestica*; DomD, Dessert *M. domestica*; SiOr, *M. sieversii* and *M. orientalis*; SylE, Eastern Europe *M. sylvestris*; SylW, Western Europe *M. sylvestris*; Bacc, *M. baccata.*

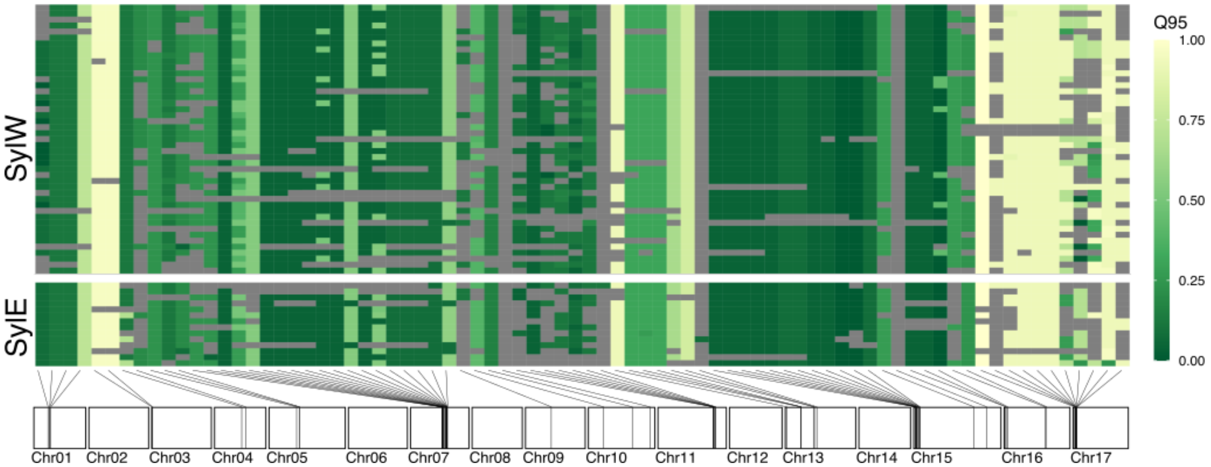


**Fig. S29. Heatmaps of Q95$_{\text{SiOr, DomC, SylW/SylE (10\%, 100\%)}}$ values using each *M. sylvestris* individual as the donor of each selective sweep region in DomC (cider apple).**

The Q95$_{\text{SiOr, DomC, SylW/SylE}}$ (10%, 100%) refers to the 95th percentile frequency of derived alleles in the DomC populations that are fixed (100% frequency) in the donor panel of SylW and SylE individuals, and occur at a frequency of less than 10% in the combined SiOr panel.The color scale indicates the value of Q95. Gray indicates sites within the region cannot calculate Q95. Chromosome diagrams below the plot are drawn to scale based on physical positions, with selective sweep regions positions along each chromosome. DomC, Cider *M. domestica*; SiOr, *M. sieversii* and *M. orientalis*; SylE, Eastern Europe *M. sylvestris*; SylW, Western Europe *M. sylvestris.*

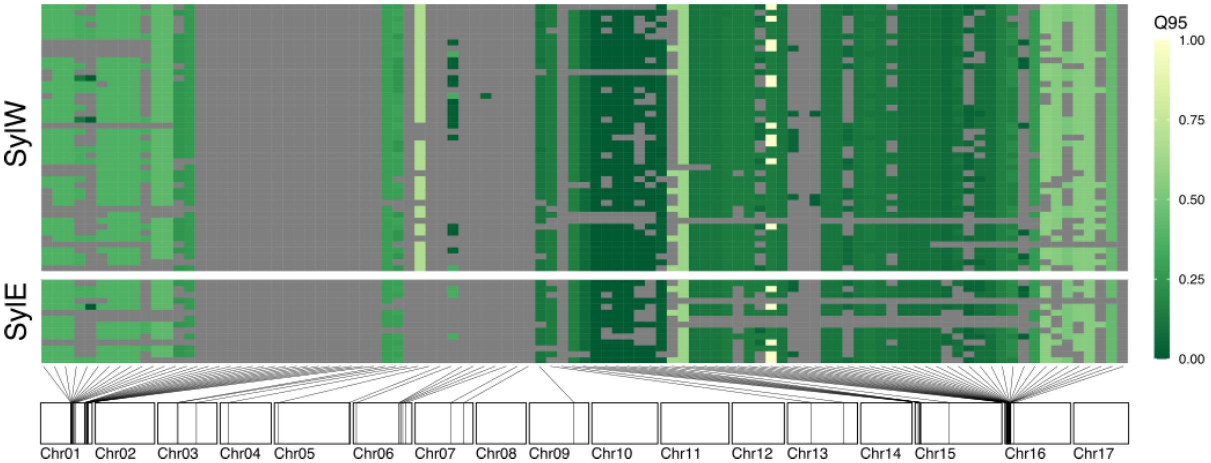


**Fig. S30. Heatmaps of Q95$_{\text{SiOr, DomD, SylW/SylE (10\%, 100\%)}}$ values using each *M. sylvestris* individual as the donor of each selective sweep region in DomD (dessert apple).**

The Q95$_{\text{SiOr, DomD, SylW/SylE}}$ (10%, 100%) refers to the 95th percentile frequency of derived alleles in the DomC populations that are fixed (100% frequency) in the donor panel of SylW and SylE individuals, and occur at a frequency of less than 10% in the combined SiOr panel. The color scale indicates the value of Q95. The color scale indicates the value of Q95. Chromosome diagrams below the plot are drawn to scale based on physical positions, with selective sweep regions positions along each chromosome. DomD, Dessert *M. domestica*; SiOr, *M. sieversii* and *M. orientalis*; Gray indicates sites within the region cannot calculate Q95. SylE, Eastern Europe *M. sylvestris*; SylW, Western Europe *M. sylvestris.*

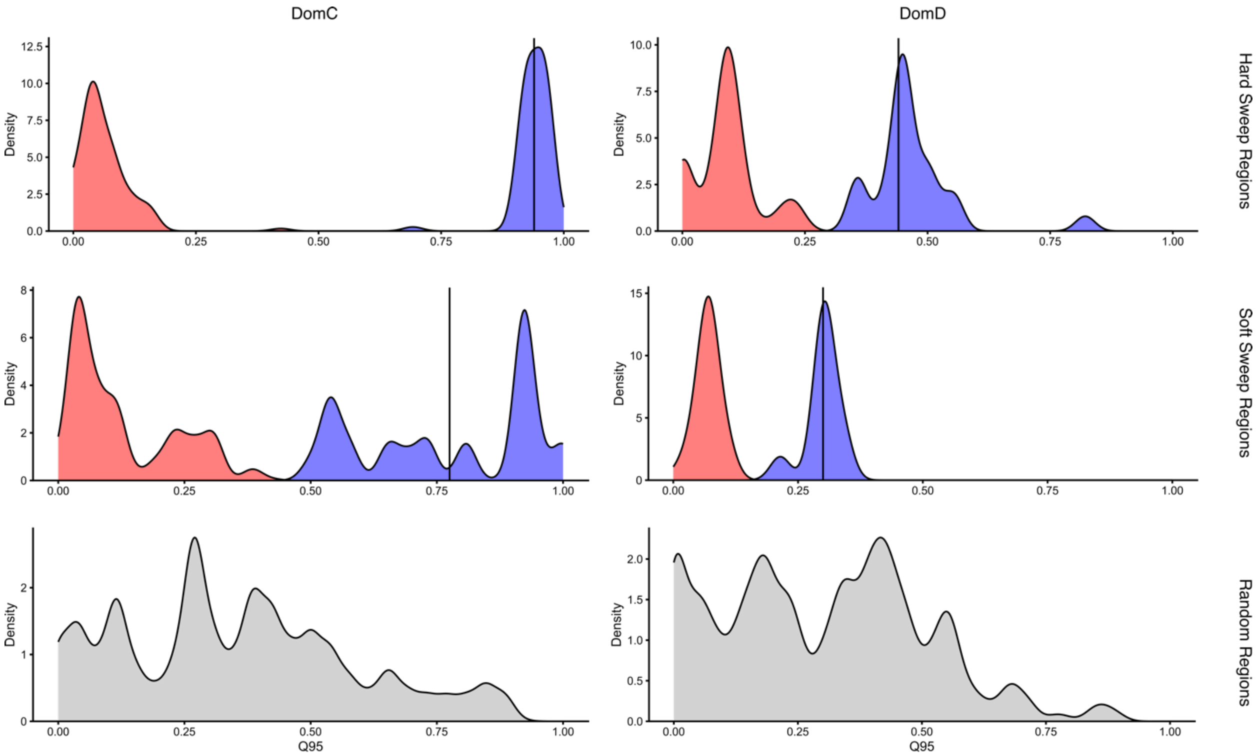


**Fig. S31. The distribution of Q95$_{\text{DomC/DomD, SiOr, SylW/SylE (10\%, 100\%)}}$ values estimated in the hard sweep, soft sweep, and randomly sampled regions.**

The *k*-means algorithm divided the hard and soft sweep region's Q95 value into two groups (red and blue). The vertical line represents the cluster centroid value of the higher Q95 value cluster (blue), serving as the threshold for detecting introgression signals.

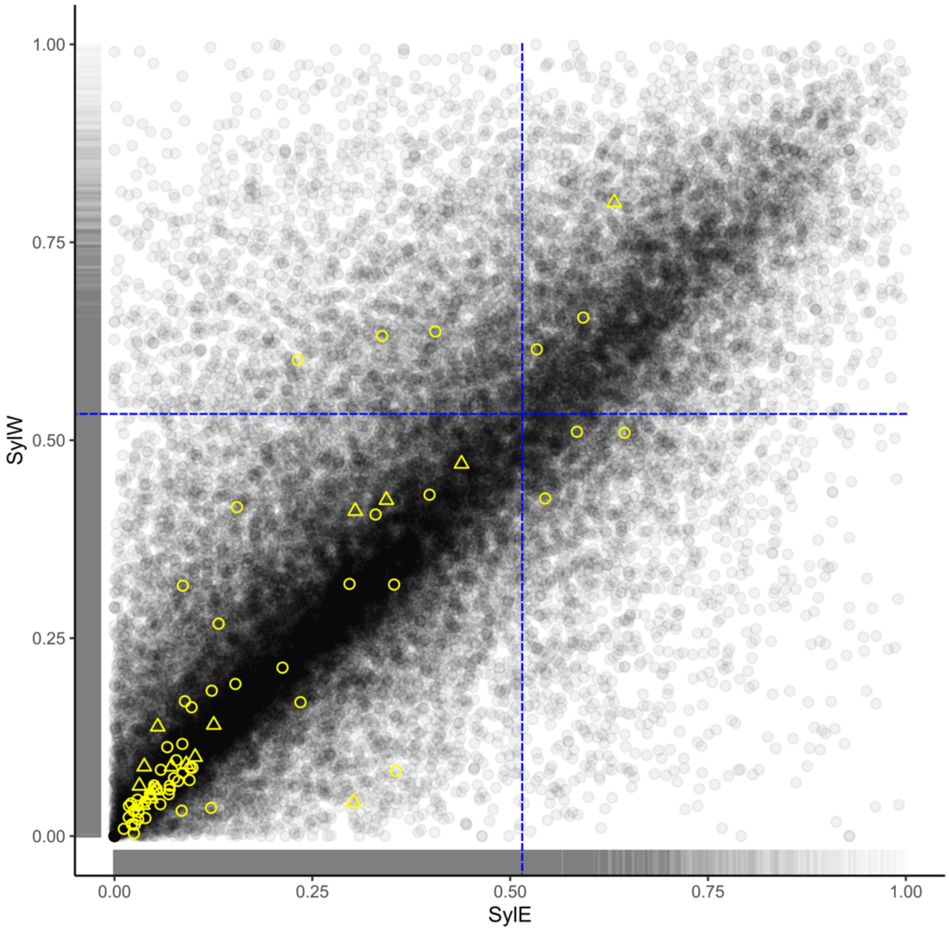


**Fig. S32. Full genome 2D- $f_d$ density plot and the $f_d$ values estimated in hard and soft sweep region of the DomC population.**

The gray dots represent the $f_d$ values (P0, Bacc; P1, SiOr; P2, DomC; P3, SylW/SylE) of all windows in the whole genome. The yellow circles and triangles represent the fd values of hard and soft sweep regions. The blue dashed line represents the threshold for outliers. DomC, Cider *M. domestica*; DomD, Dessert *M. domestica*; SiOr, *M. sieversii* and *M. orientalis*; SylE, Eastern Europe *M. sylvestris*; SylW, Western Europe *M. sylvestris*.

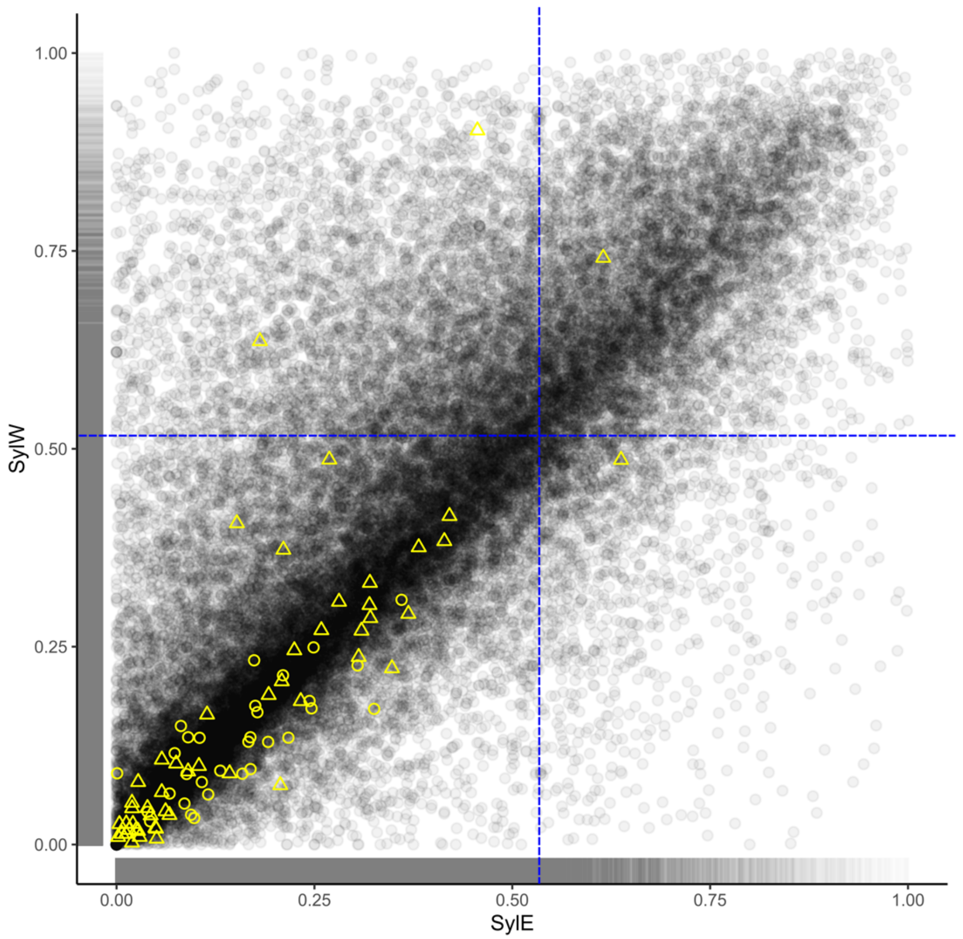


**Fig. S33. Full genome 2D $f_d$ density plot and the hard and soft sweep region $f_d$ values of DomD population.**

The gray dots represent the $f_d$ values (P0, Bacc; P1, SiOr; P2, DomD; P3, SylW/SyE) of all windows in the whole genome. The yellow circles and triangles represent the hard and soft sweep region $f_d$ values. The blue dashed line represents the threshold for outliers. DomD, Dessert *M. domestica*; SiOr, *M. sieversii* and *M. orientalis*; SylE, Eastern Europe *M. sylvestris*; SylW, Western Europe *M. sylvestris*.

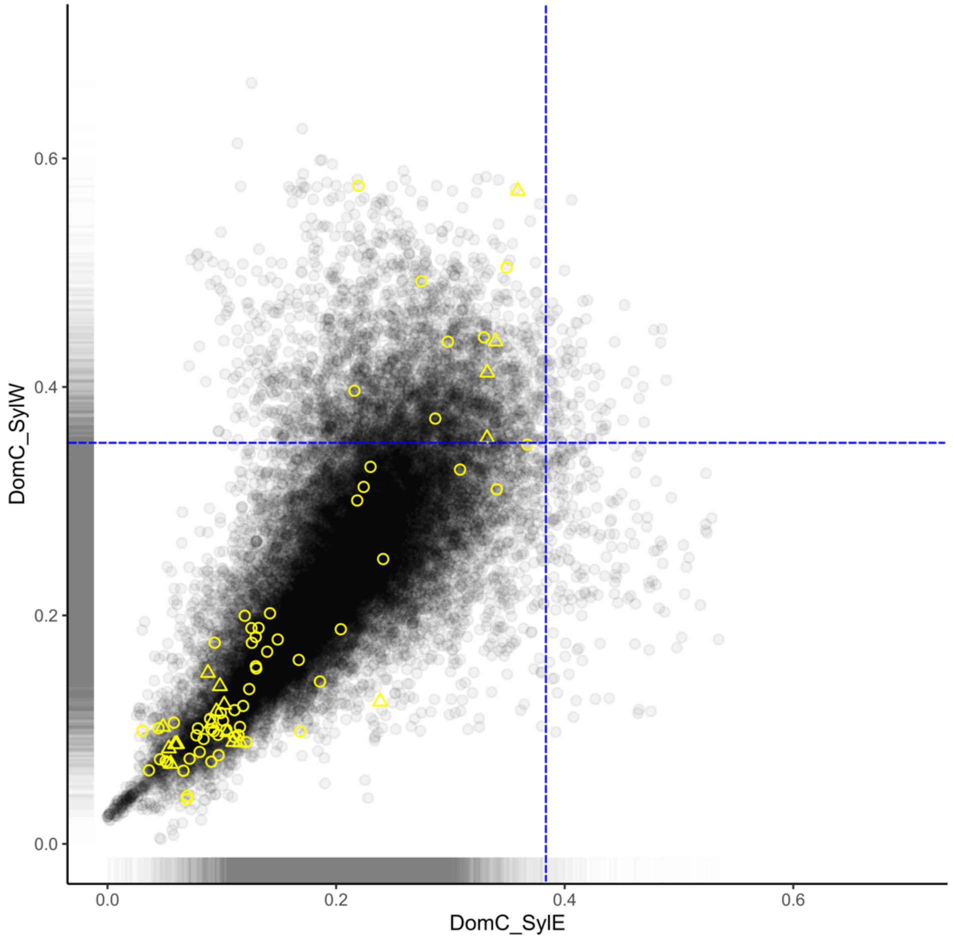


**Fig. S34. Full genome 2D topology weighting density plot and hard and soft sweep region topology weighting values of the DomC population.**

The gray dots represent the topology weighting of phylogenetic groups, indicating that DomC is introgressed by SylW (x-axis) or SylE (y-axis) for all tested windows across the entire genome. The yellow circles and triangles represent the same values for hard and soft sweep regions. The blue dashed line represents the threshold for outliers. DomC, Cider *M. domestica*; SiOr, *M. sieversii* and *M. orientalis*; SylE, Eastern Europe *M. sylvestris*; SylW, Western Europe *M. sylvestris*.

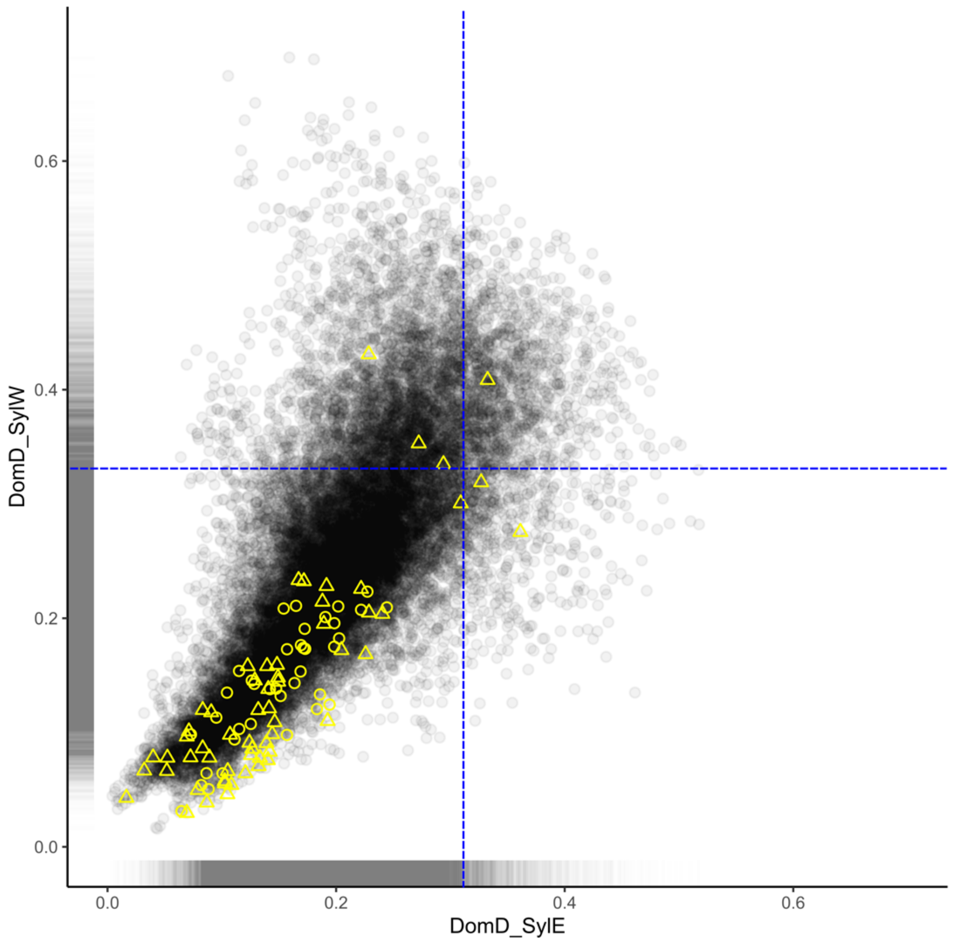


**Fig. S35. Full genome 2D topology weighting density plot and hard and soft sweep region topology weighting values of DomD population.**
The gray dots represent the topology weighting of phylogenies groups that indicated DomD are introgressed by the SylW (x-axis) or SylE (y-axis) for all tested windows in the whole genome. The yellow circles and triangles represent the same values on hard and soft sweep regions. The blue dashed line represents the threshold for outliers. DomD, Dessert *M. domestica*; SiOr, *M. sieversii* and *M. orientalis*; SylE, Eastern Europe *M. sylvestris*; SylW, Western Europe *M. sylvestris*.

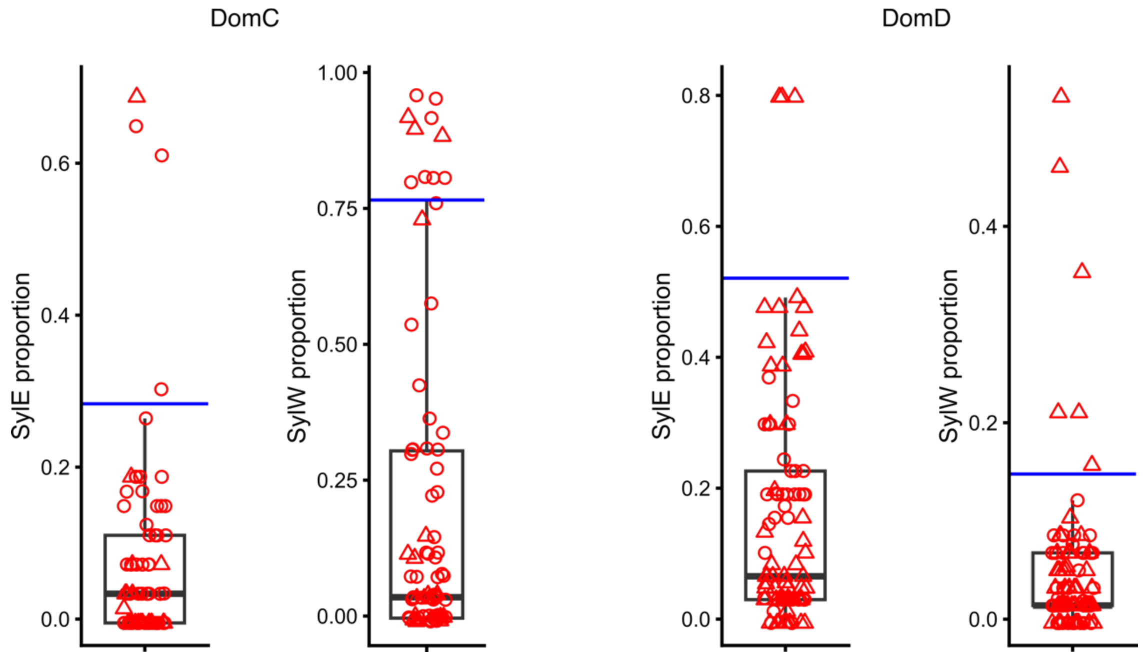


**Fig. S36. Local ancestry proportion inference of Eastern and Western *M. sylvestris* estimated with RFMix in DomC and DomD regions detected under selection.**
The red triangles and circles represent the hard and soft sweep region values. The blue line represents the threshold for outliers. DomC, Cider *M. domestica*; DomD, Dessert *M. domestica*; SiOr, *M. sieversii* and *M. orientalis*; SylE, Eastern Europe *M. sylvestris*; SylW, Western Europe *M. sylvestris*.

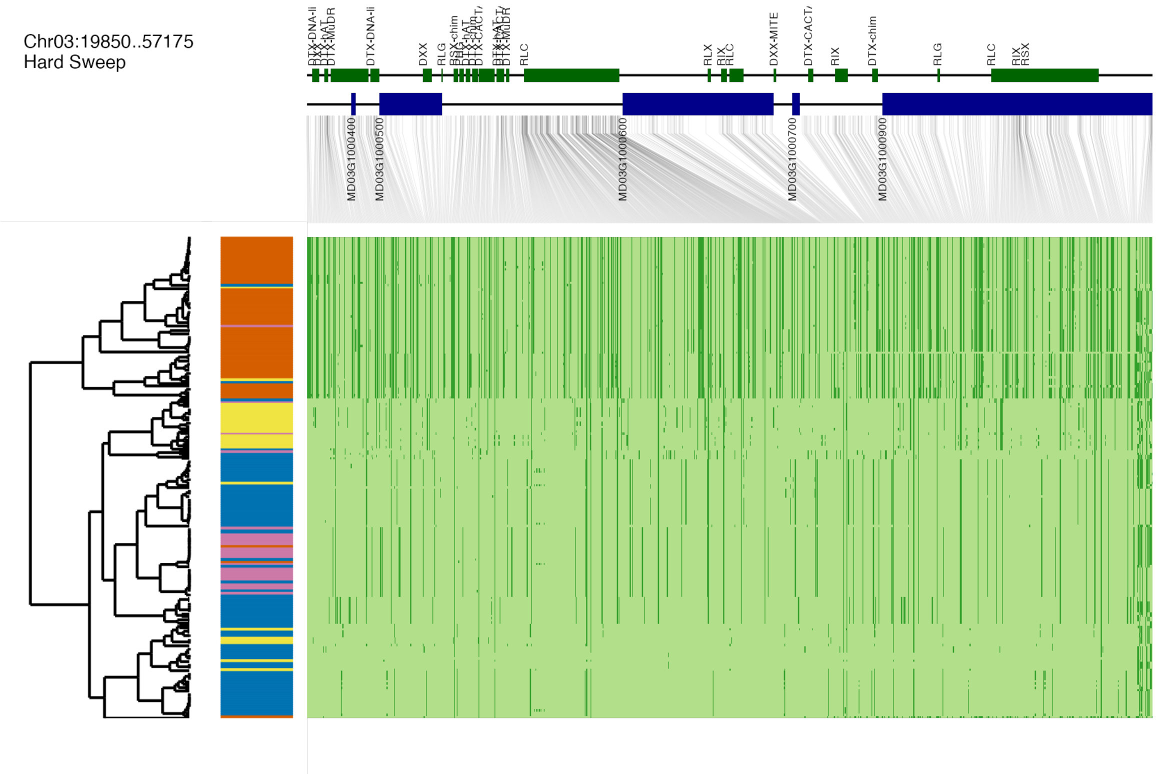


**Fig. S37. Haplotype structure of candidate regions with strong evidence for DomC adaptive introgression.**
For each region, haplotypes are clustered according to PCA and complete linkage method using *dudi.pca()* and *hclust()* (114). The left side is the dendrogram, and haplotypes from different populations are represented in different colors. The upper part shows the genes in this region, and the gene IDs come from the annotations of the GDDH13 v1.1 genome (37). DomC, Cider *M. domestica*; SiOr, *M. sieversii* and *M. orientalis*; SylE, Eastern Europe *M. sylvestris*; SylW, Western Europe *M. sylvestris*.

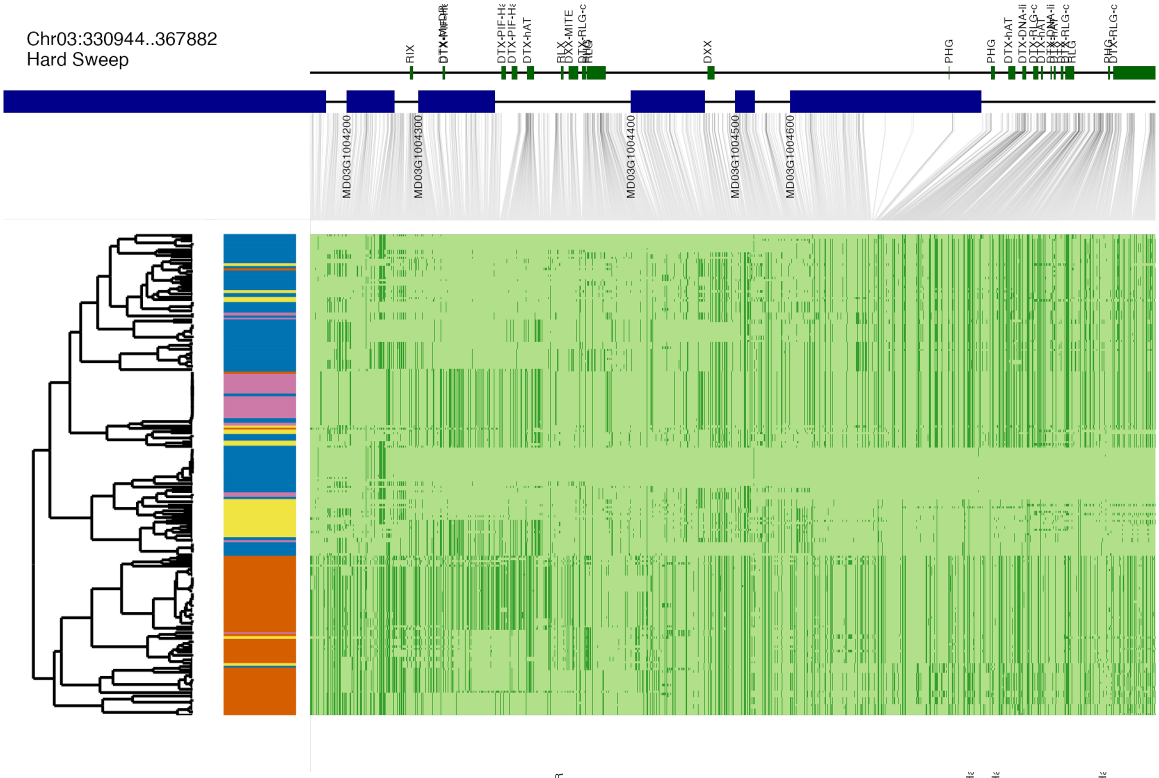


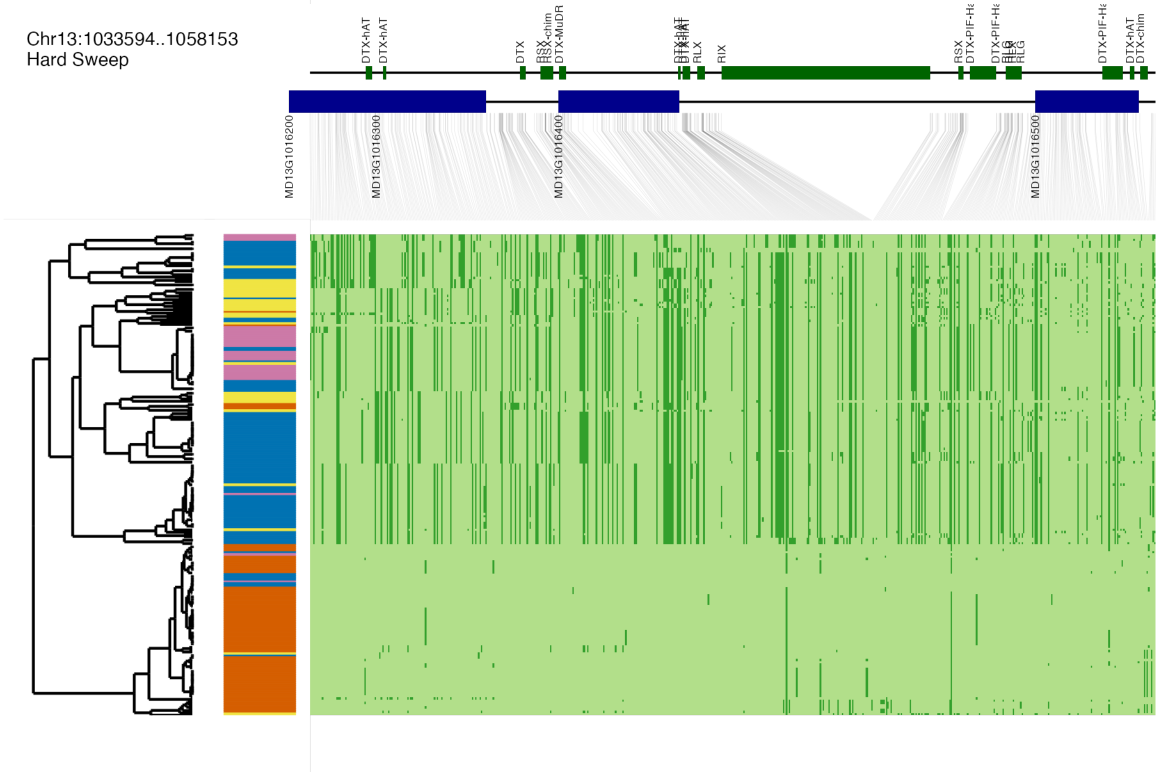


**Fig. S37 (continued)**

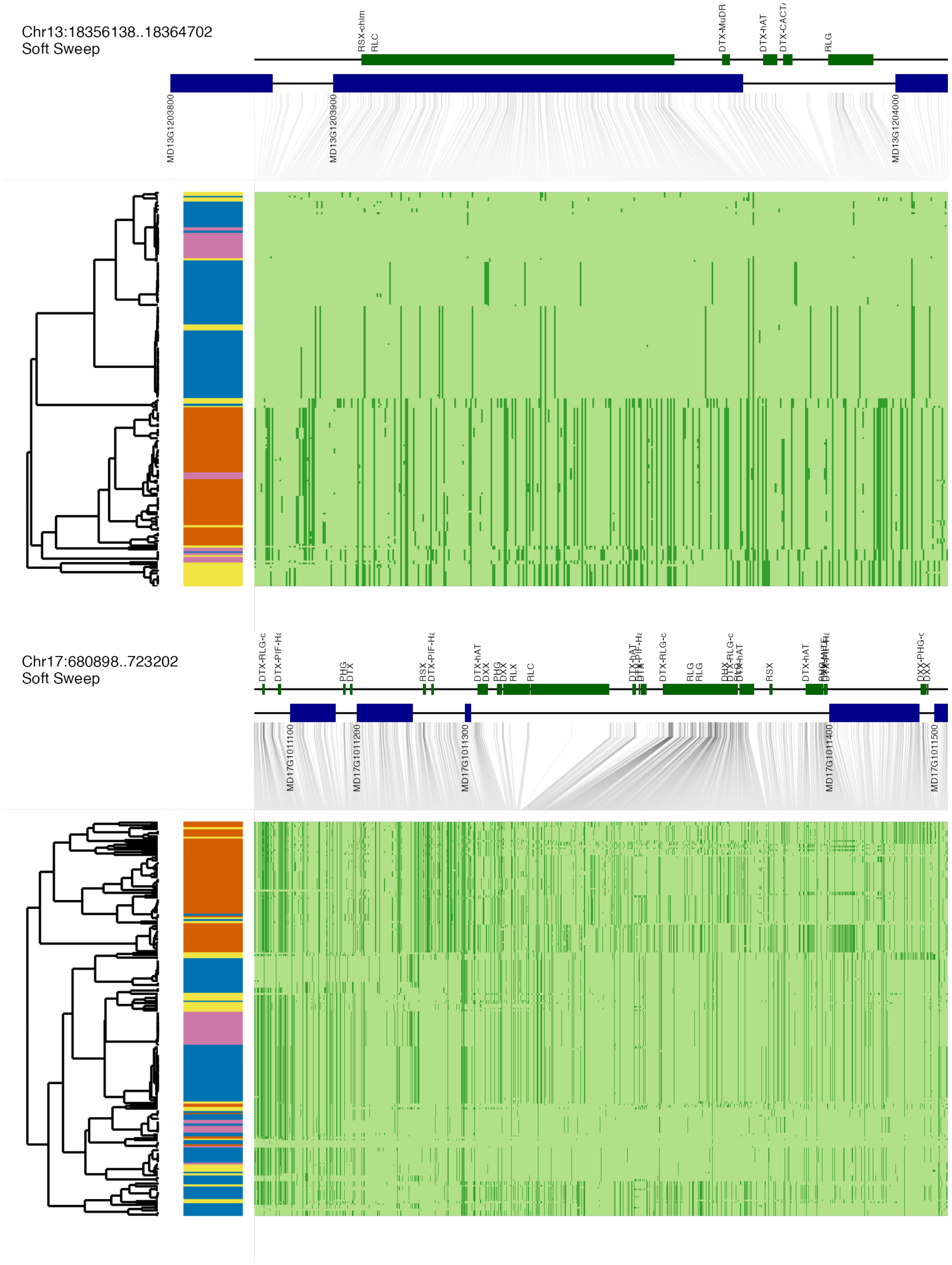


**Fig. S37 (continued)**

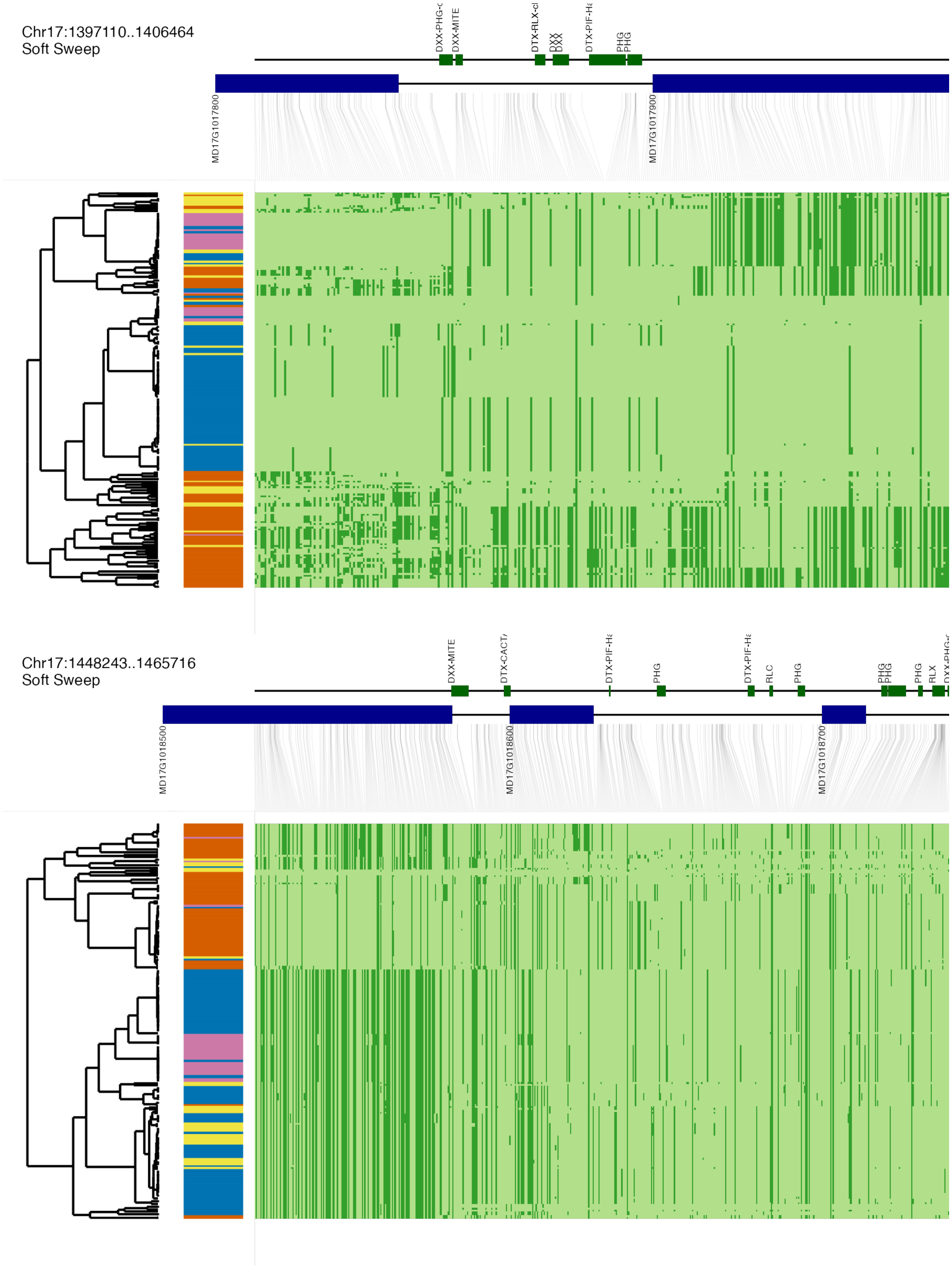


**Fig. S37 (continued)**

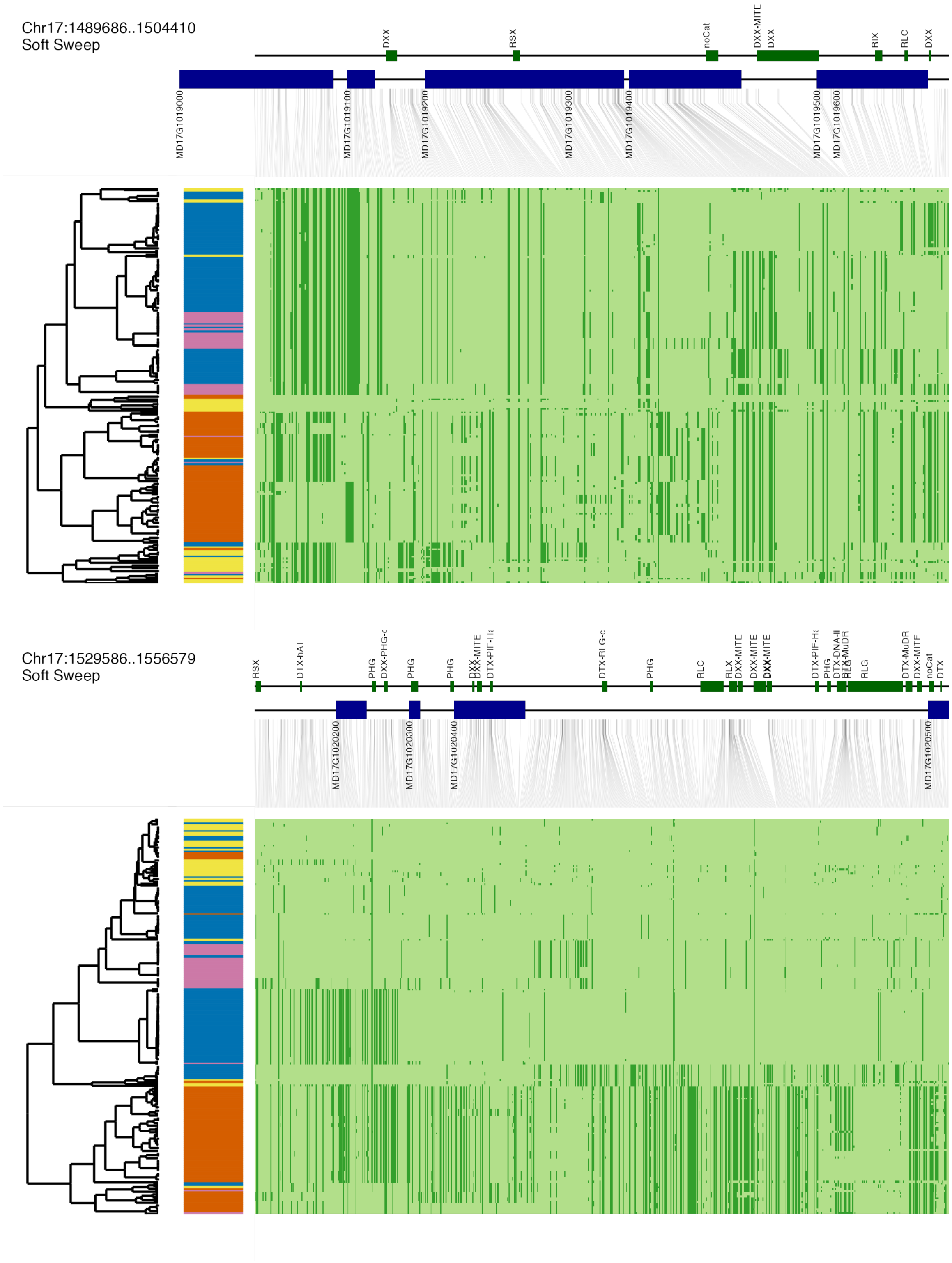


**Fig. S37 (continued)**

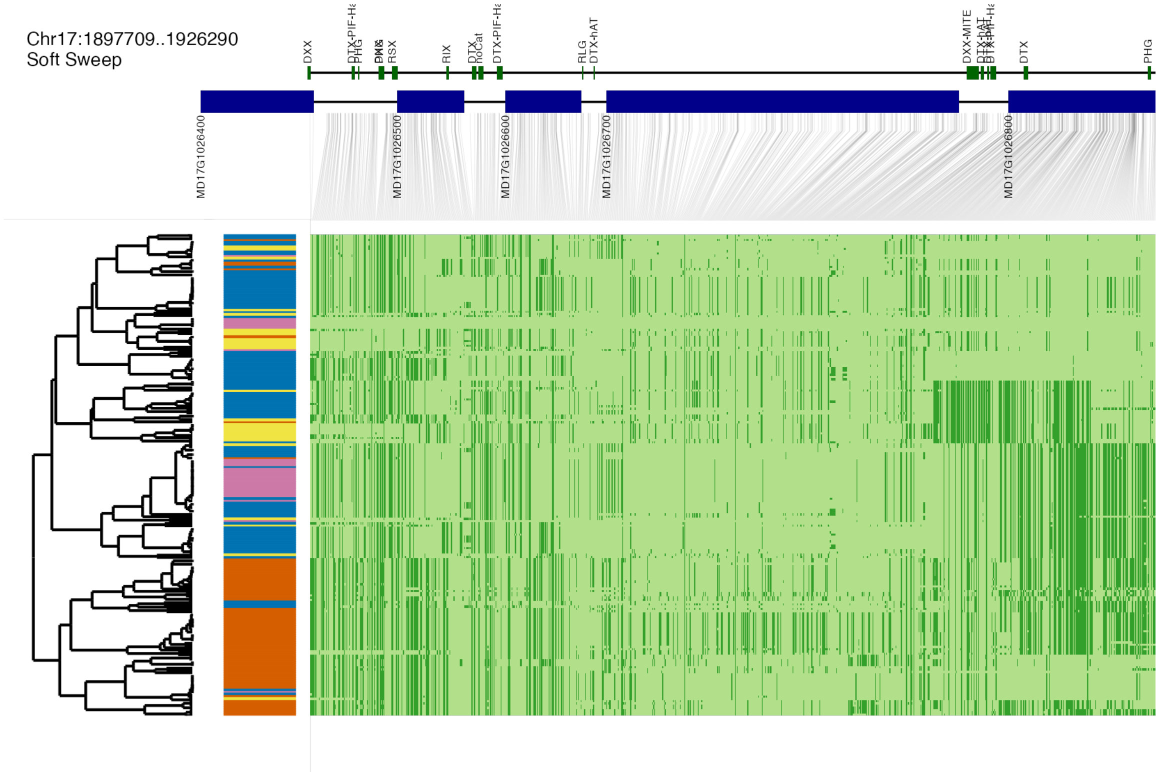


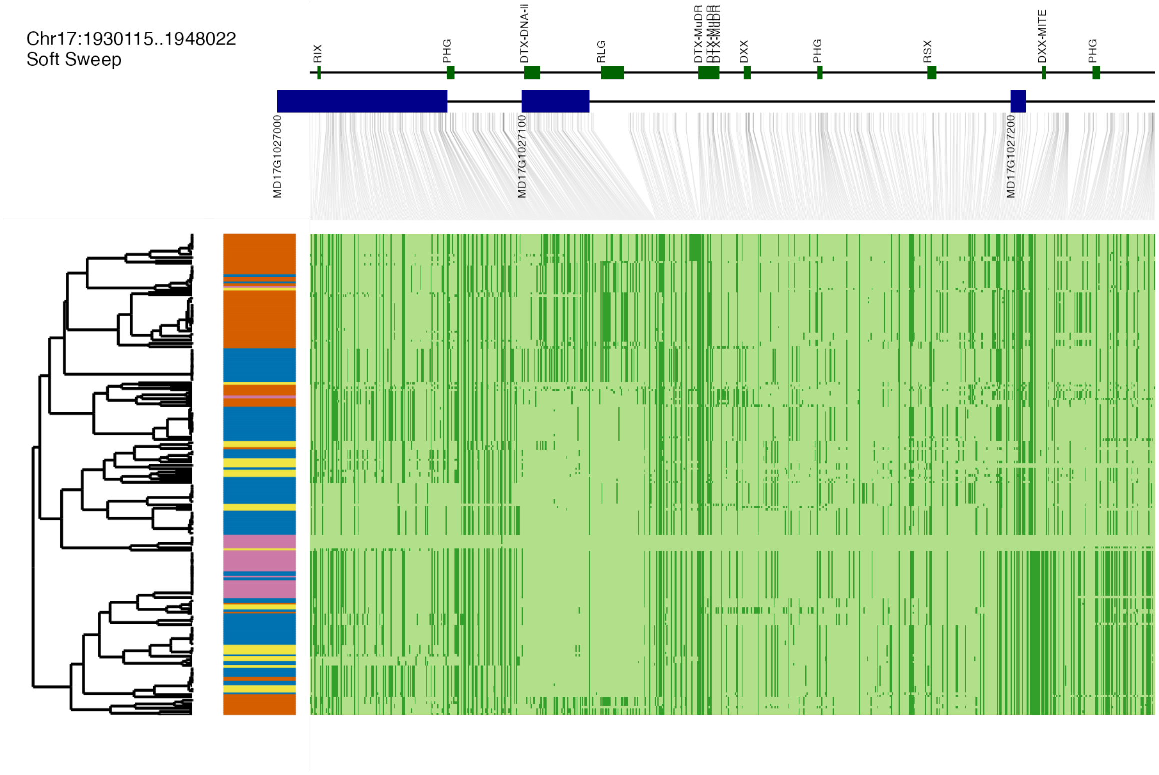


**Fig. S37 (continued)**

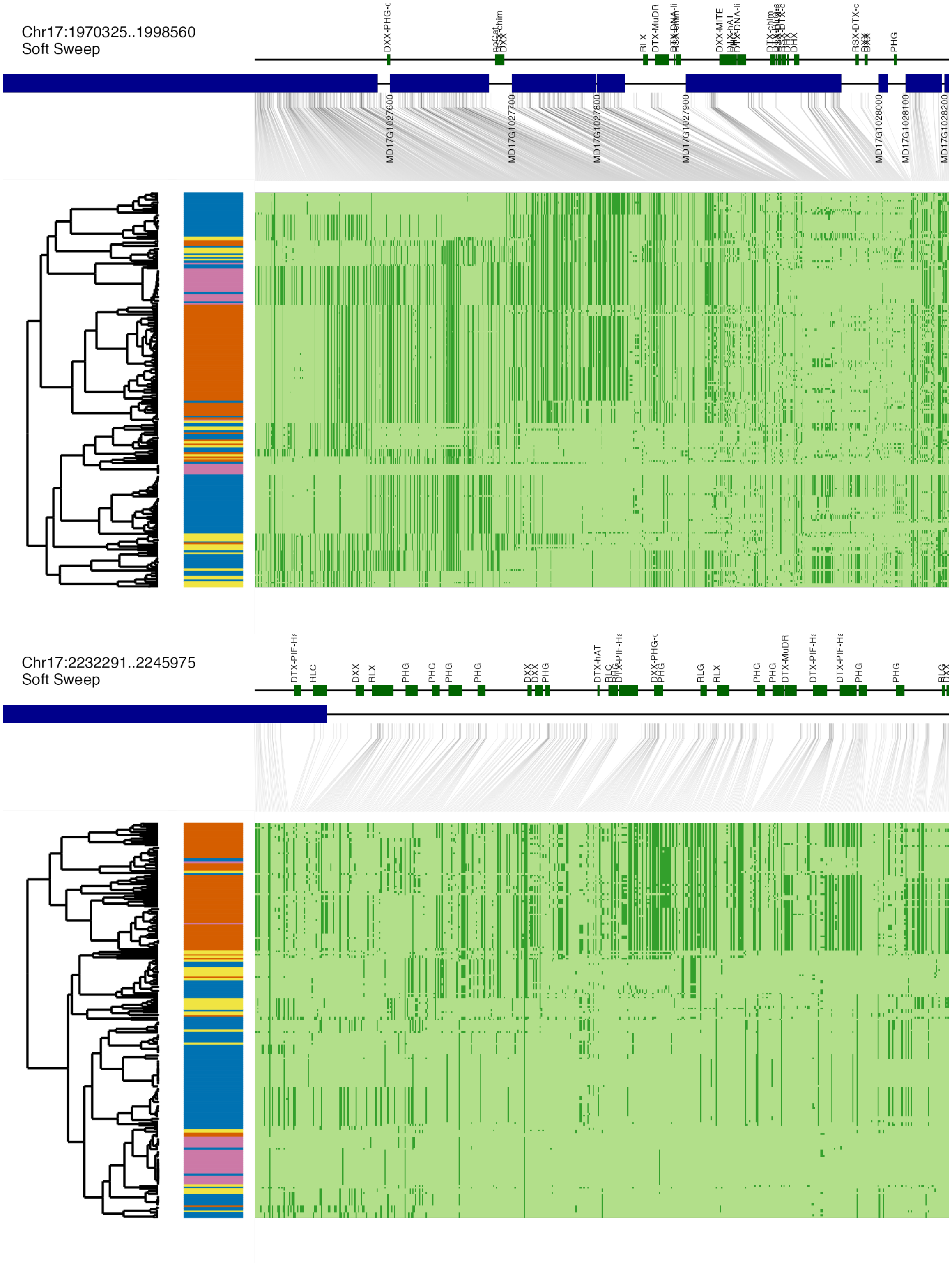


**Fig. S37 (continued)**

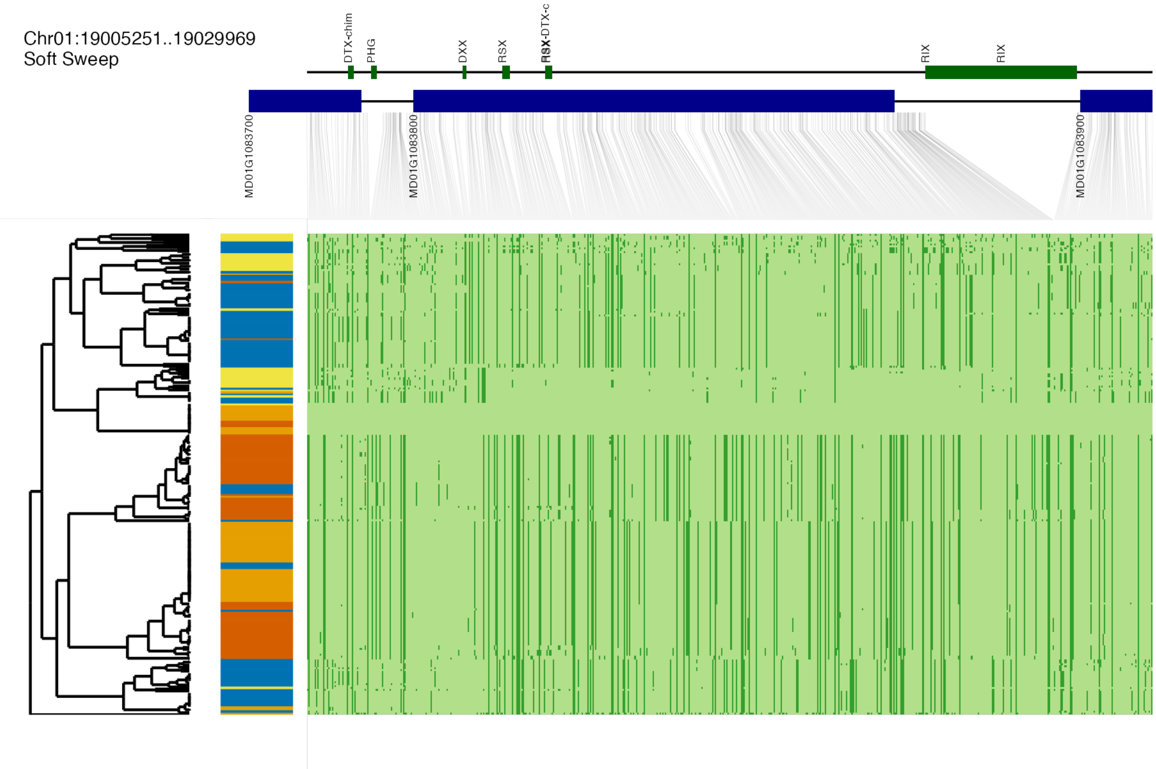


**Fig. S38. Haplotype structure of candidate regions with strong evidence for DomD adaptive introgression.**
For each region, haplotypes are clustered according to PCA analysis and complete linkage method using *dudi.pca()* and *hclust()*(114). The left side is the dendrogram, and haplotypes from different populations are represented in different colors. The upper part shows the genes in this region, and the gene IDs come from the annotations of the GDDH13 v1.1 genome (37). DomD, Dessert *M. domestica*; SiOr, *M. sieversii* and *M. orientalis*; SylE, Eastern Europe *M. sylvestris*; SylW, Western Europe *M. sylvestris*.

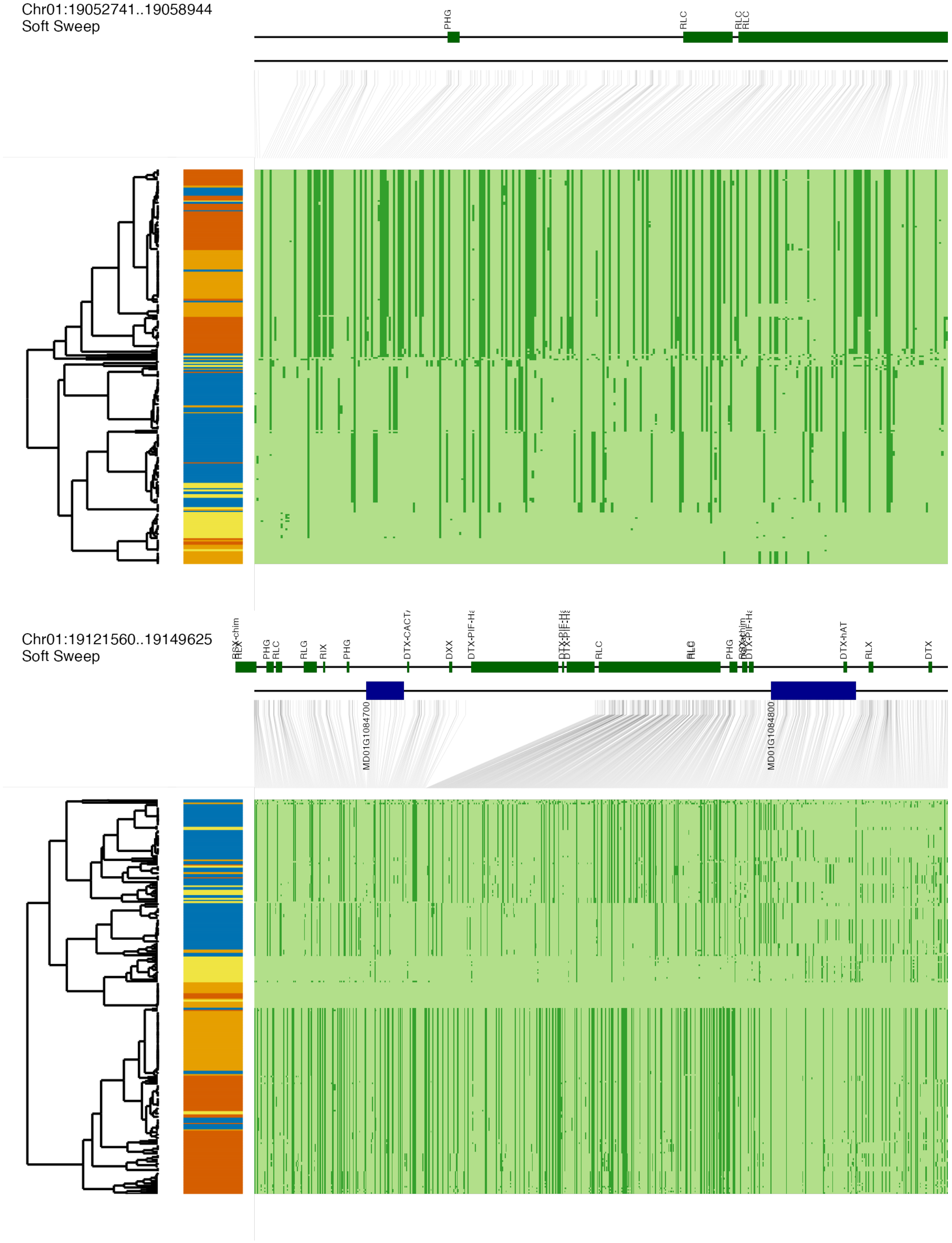


**Fig. S38 (continued)**

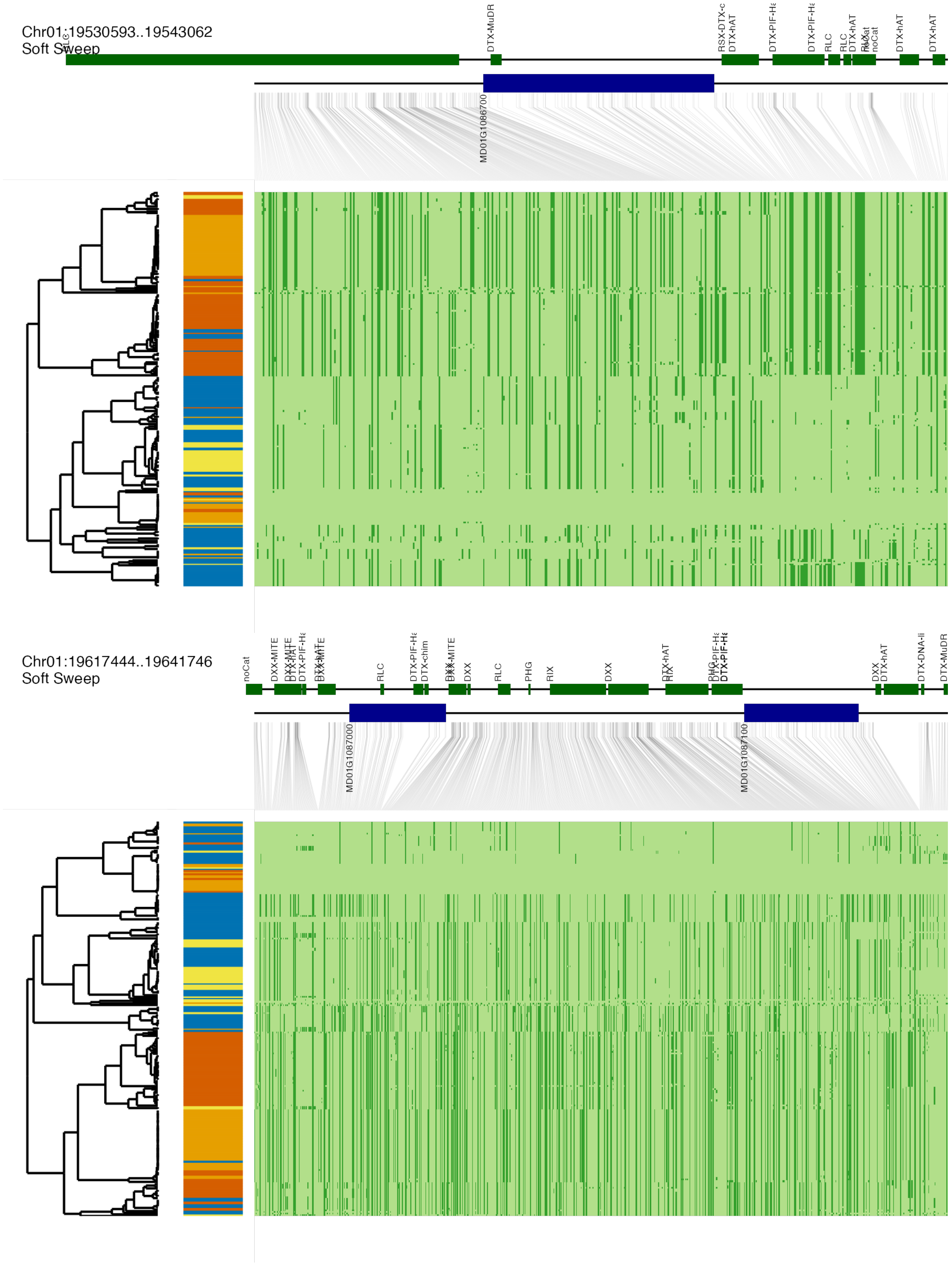


**Fig. S38 (continued)**

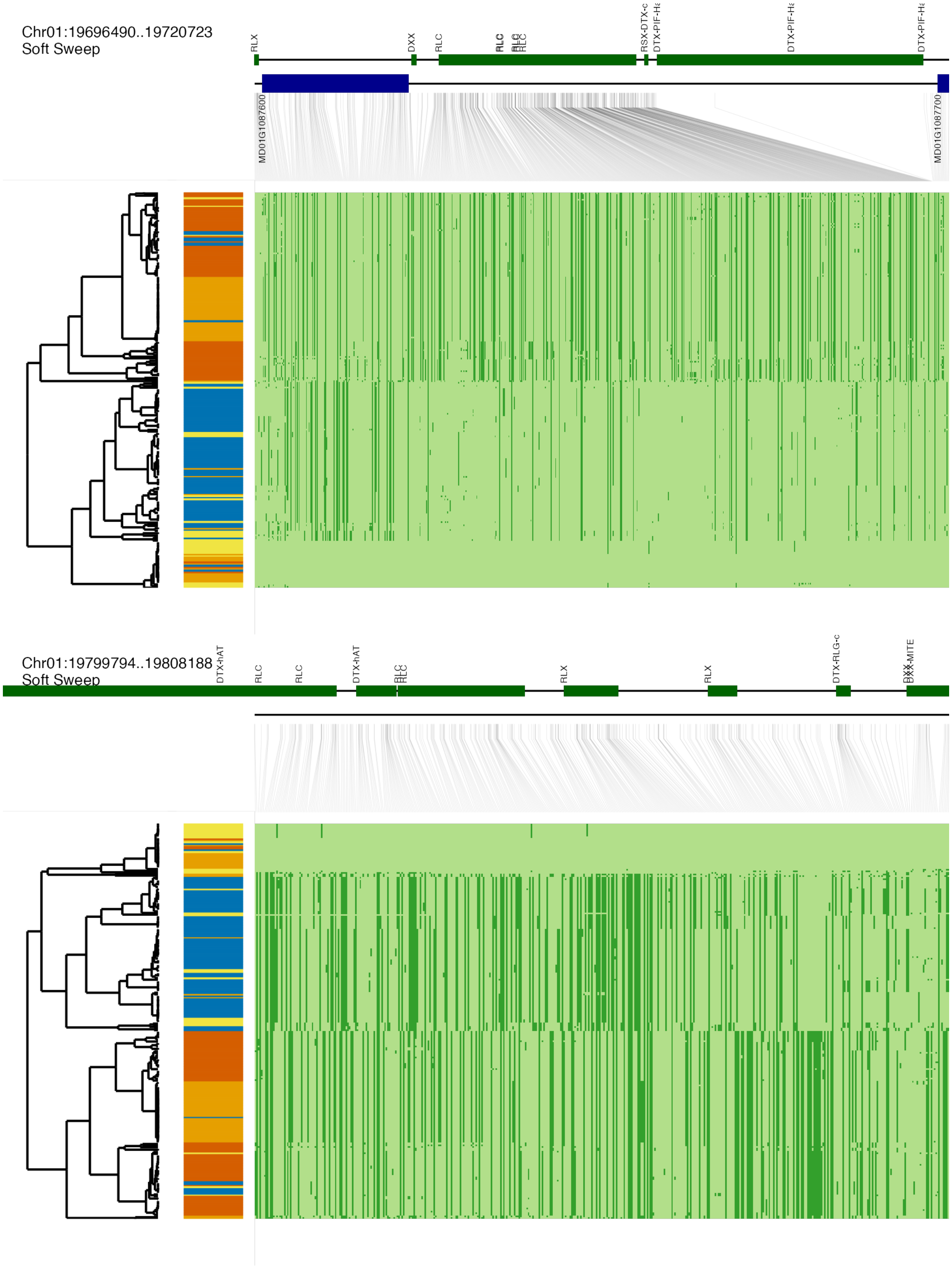


**Fig. S38 (continued)**

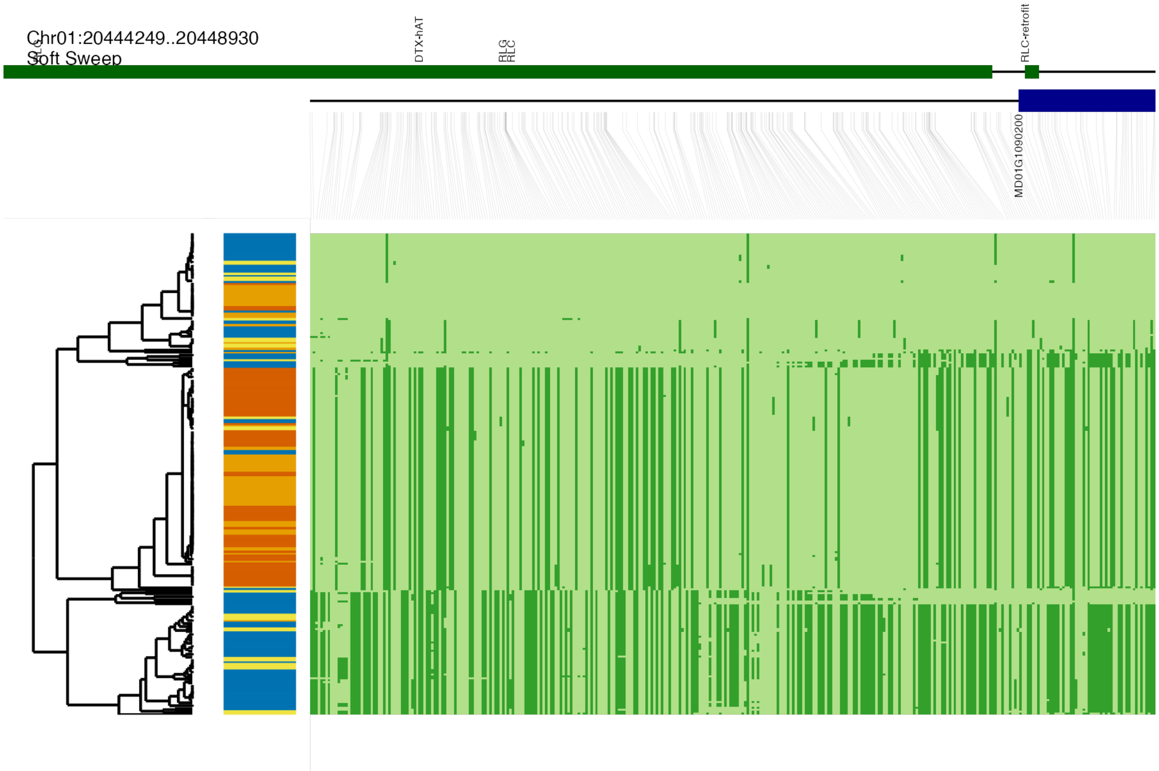


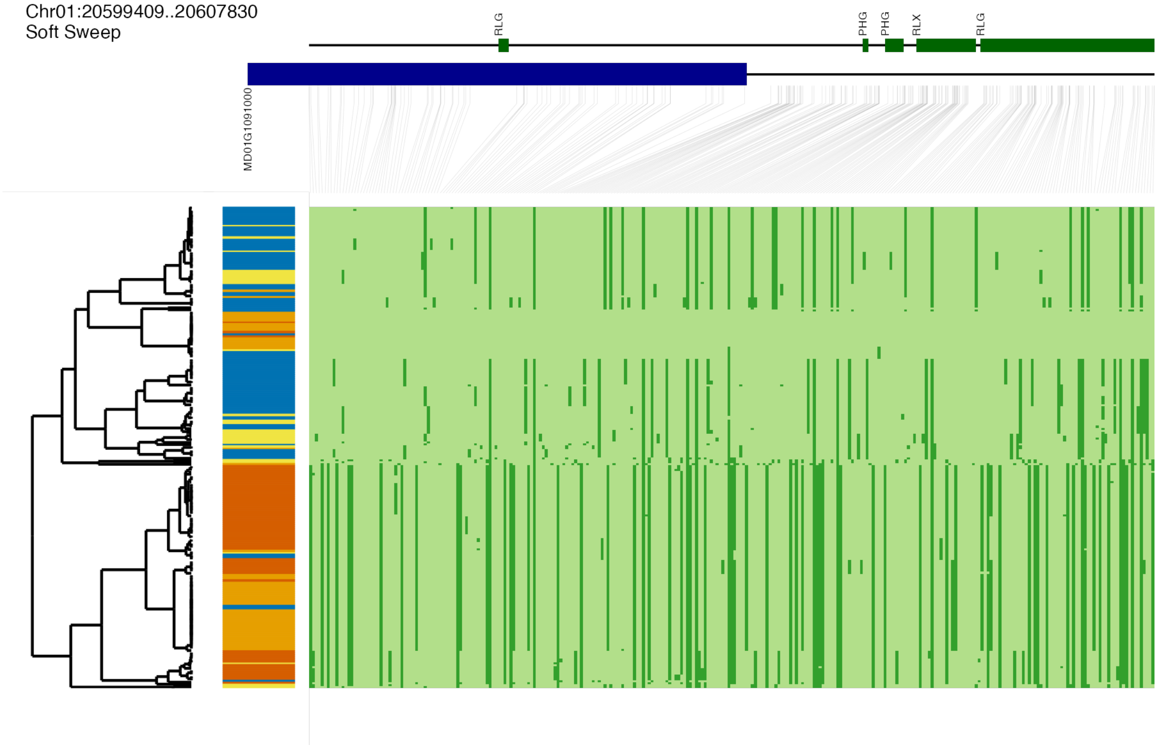


**Fig. S38 (continued)**

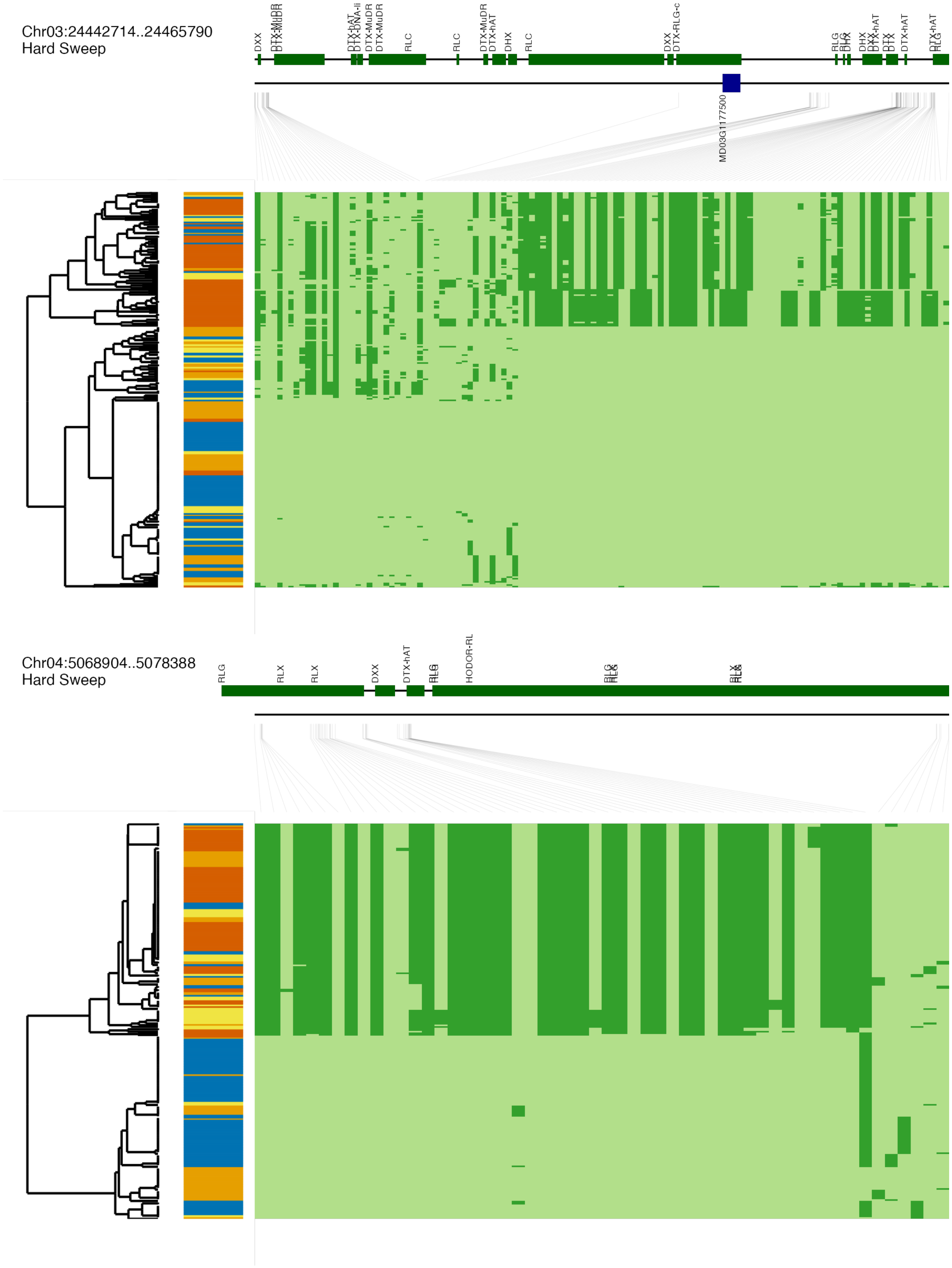


**Fig. S38 (continued)**

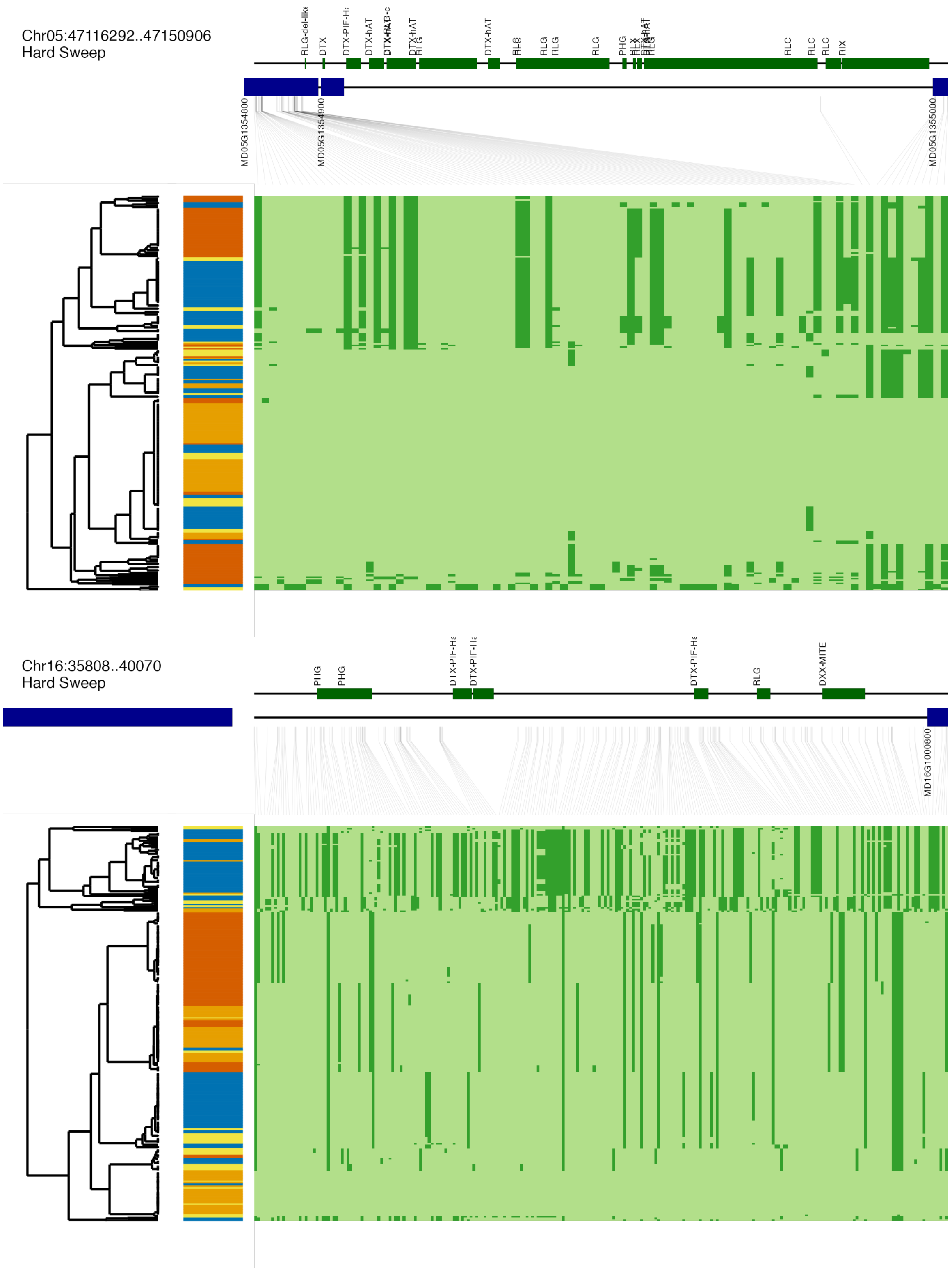


**Fig. S38 (continued)**

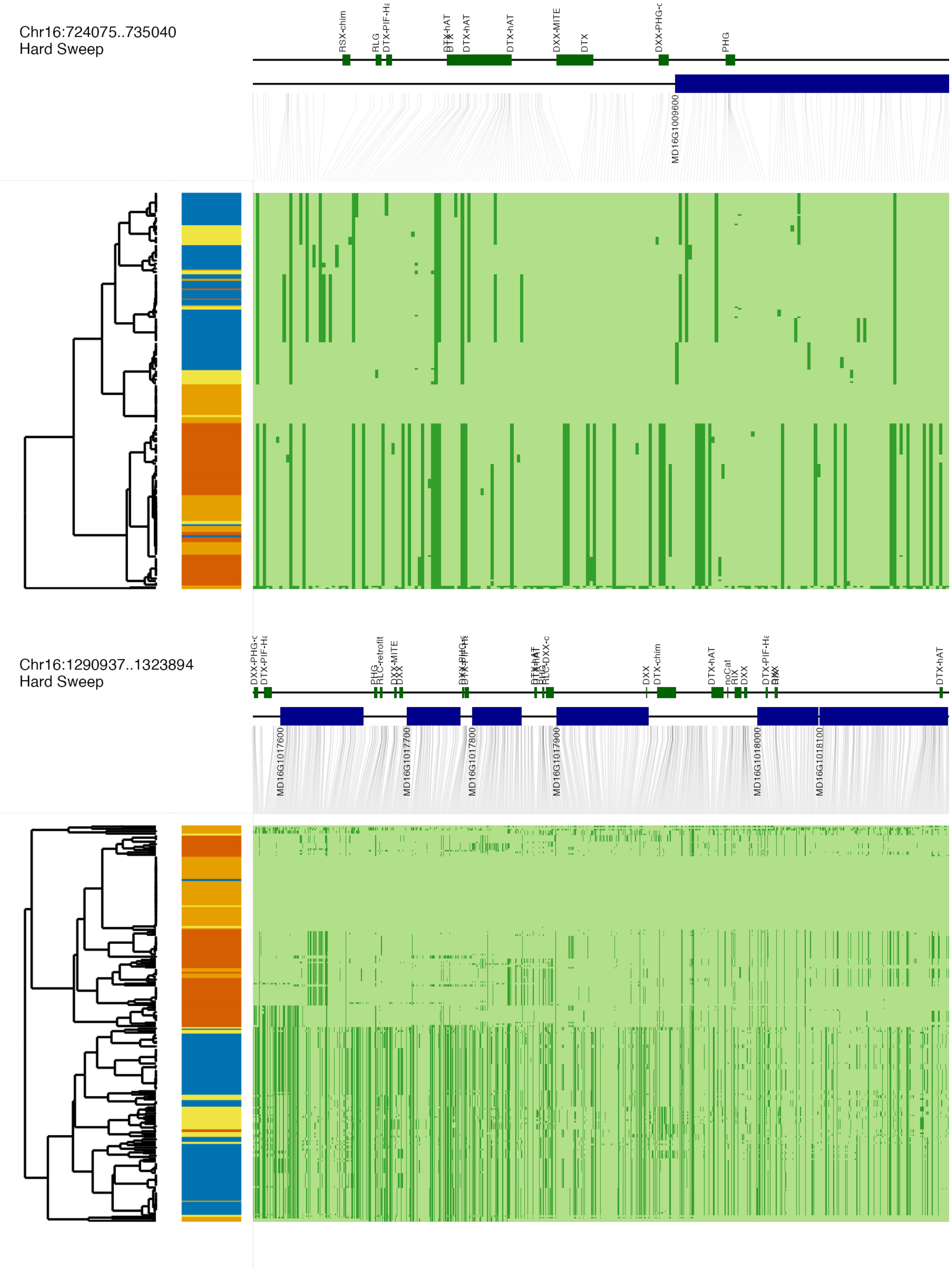


**Fig. S38 (continued)**

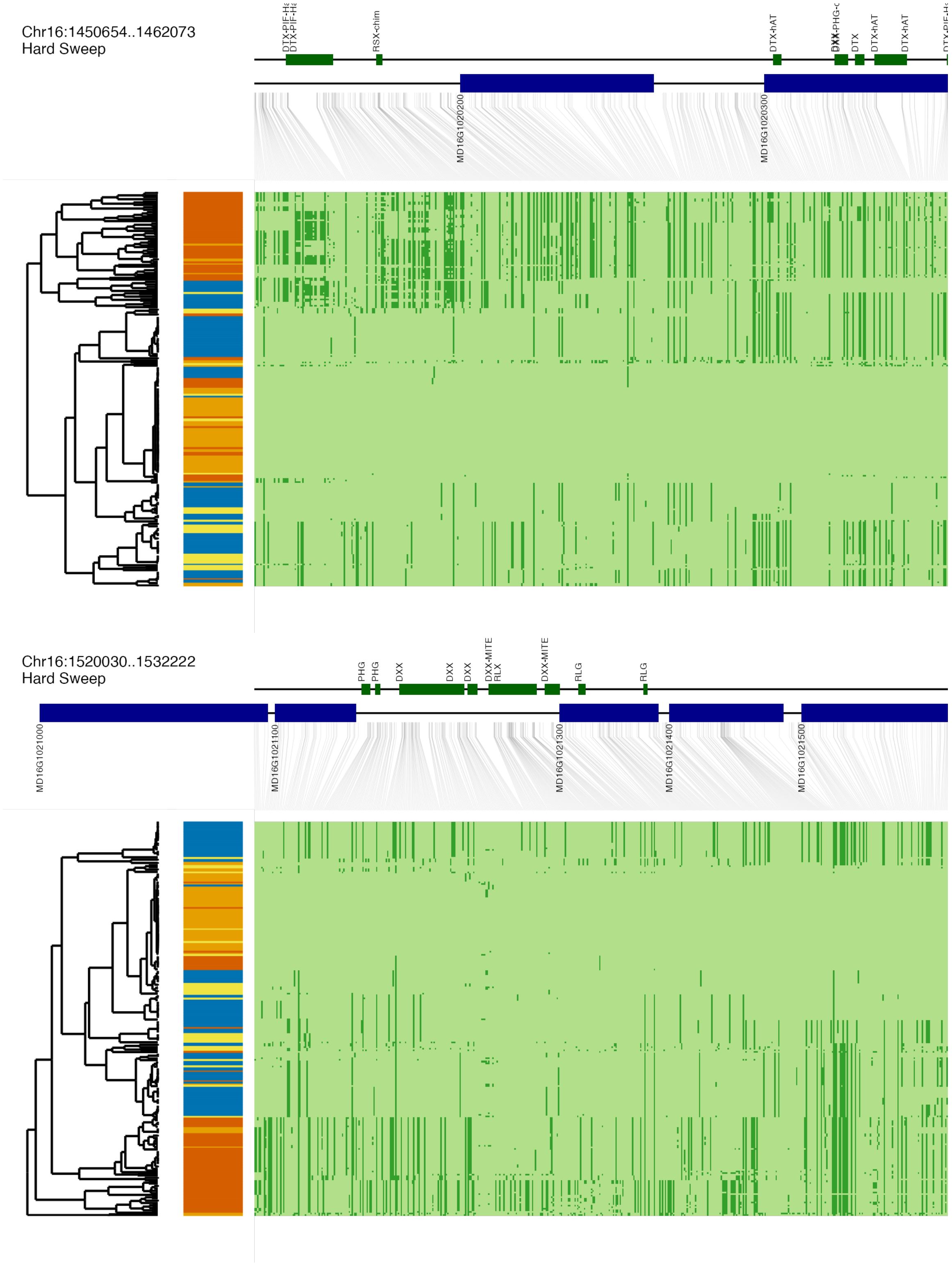


**Fig. S38 (continued)**

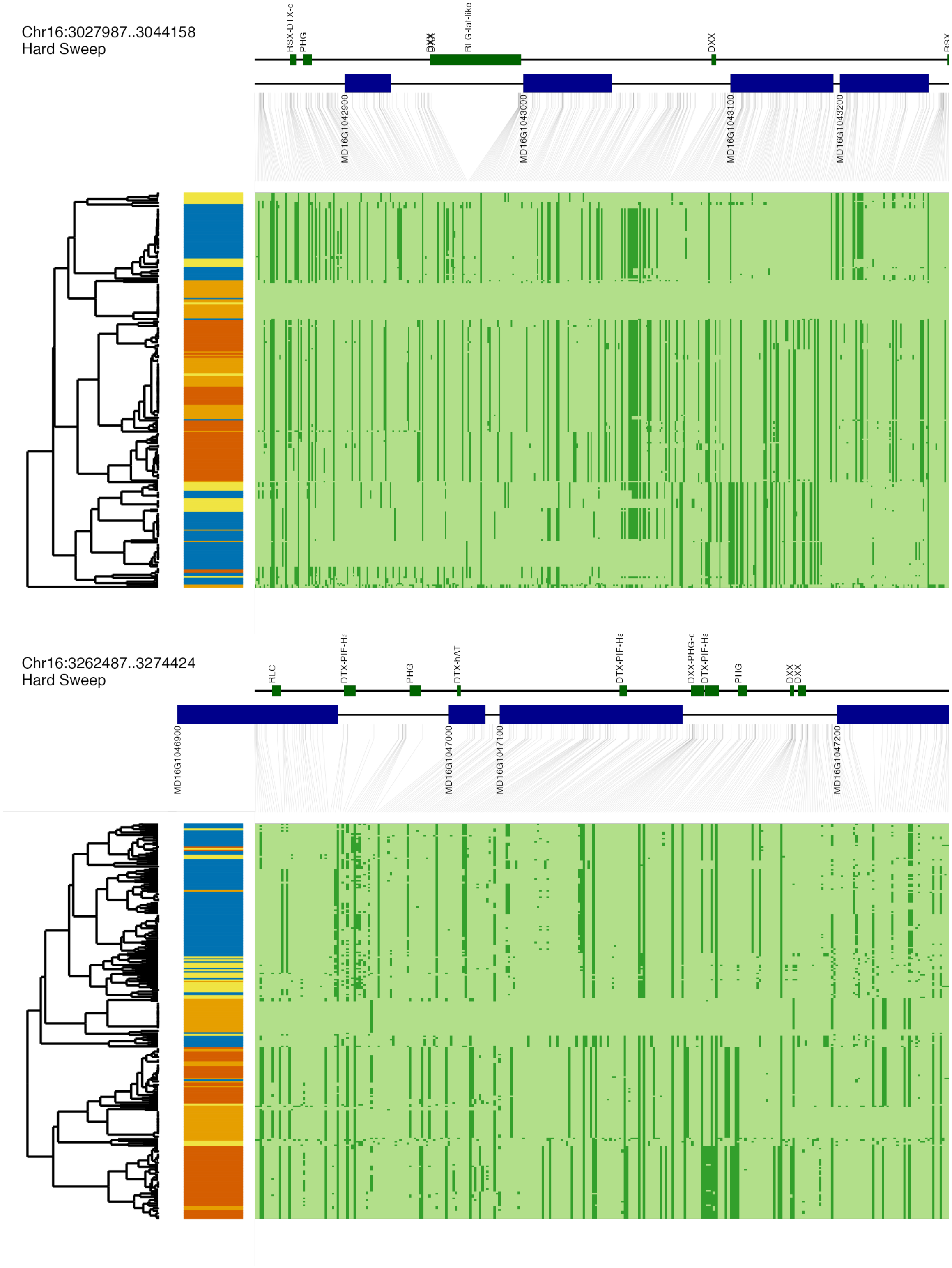


**Fig. S38 (continued)**

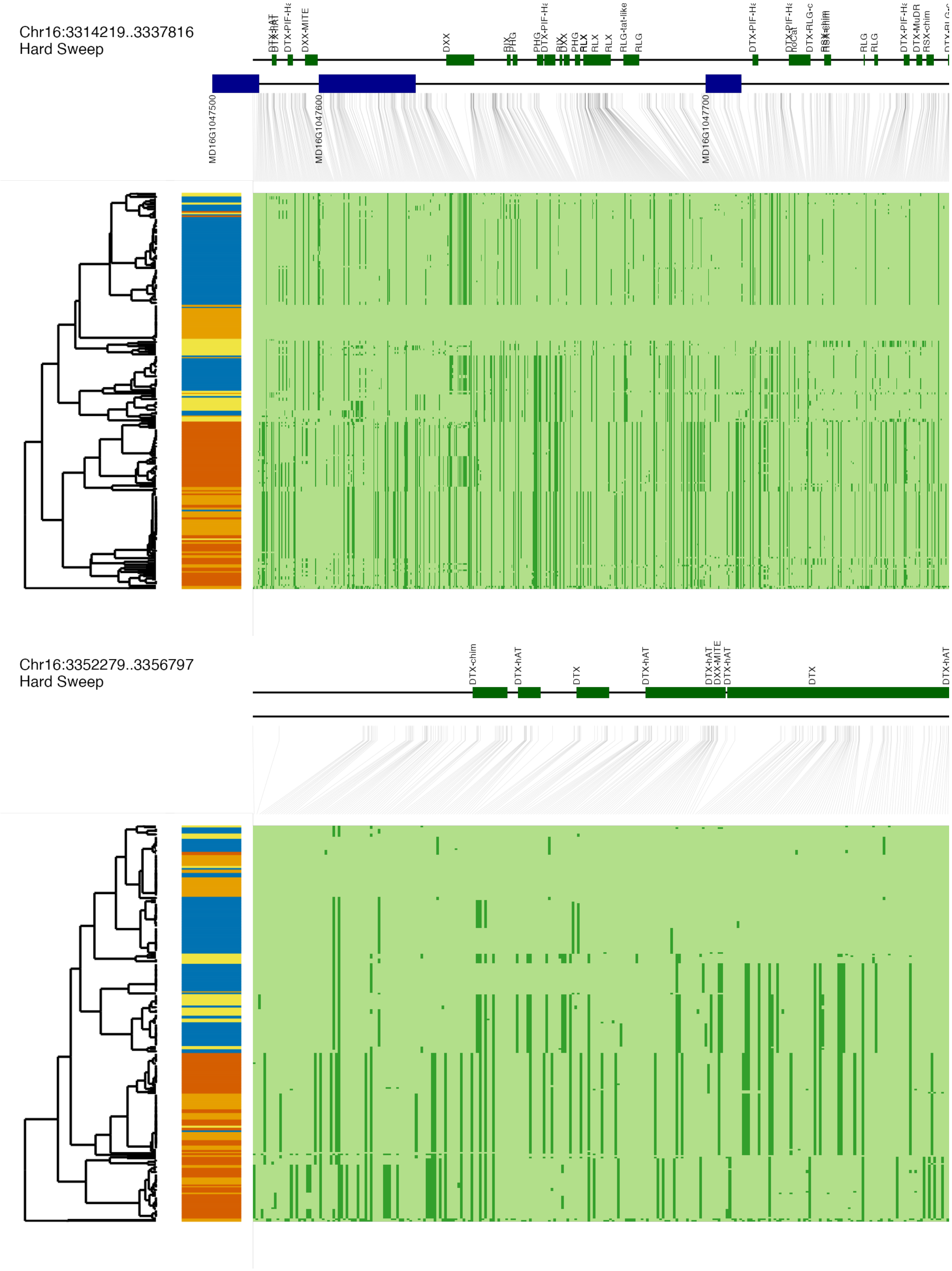


**Fig. S38 (continued)**

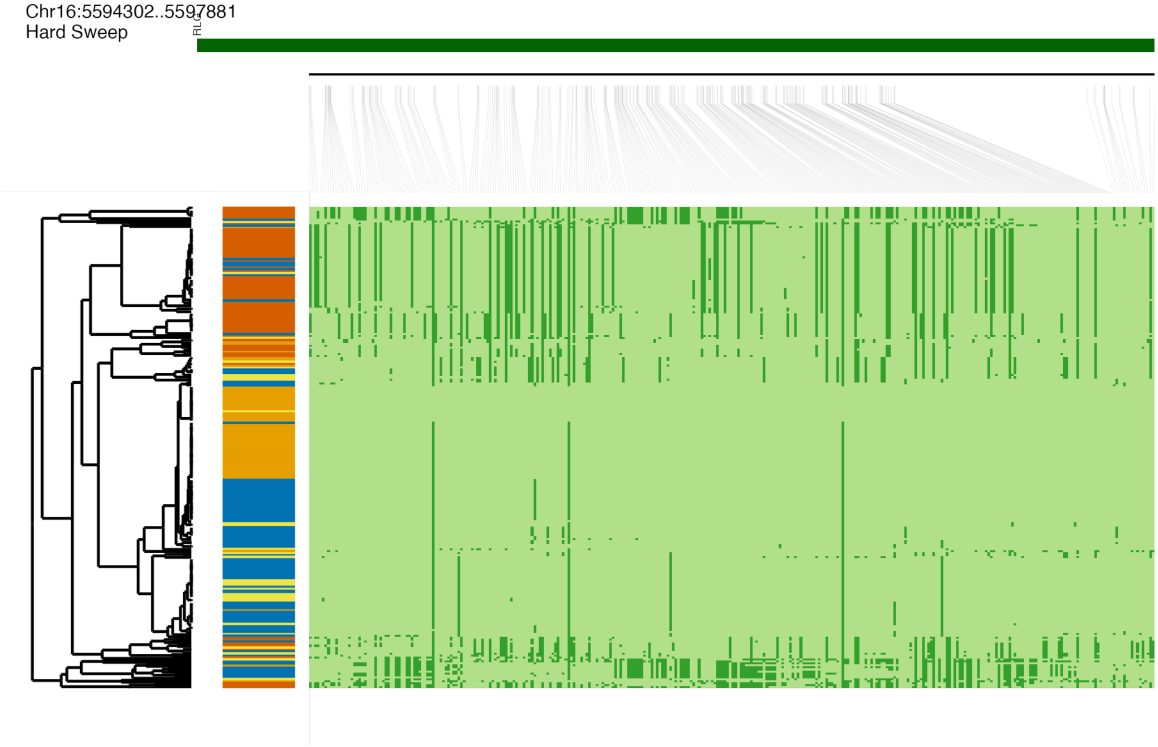


**Fig. S38 (continued)**

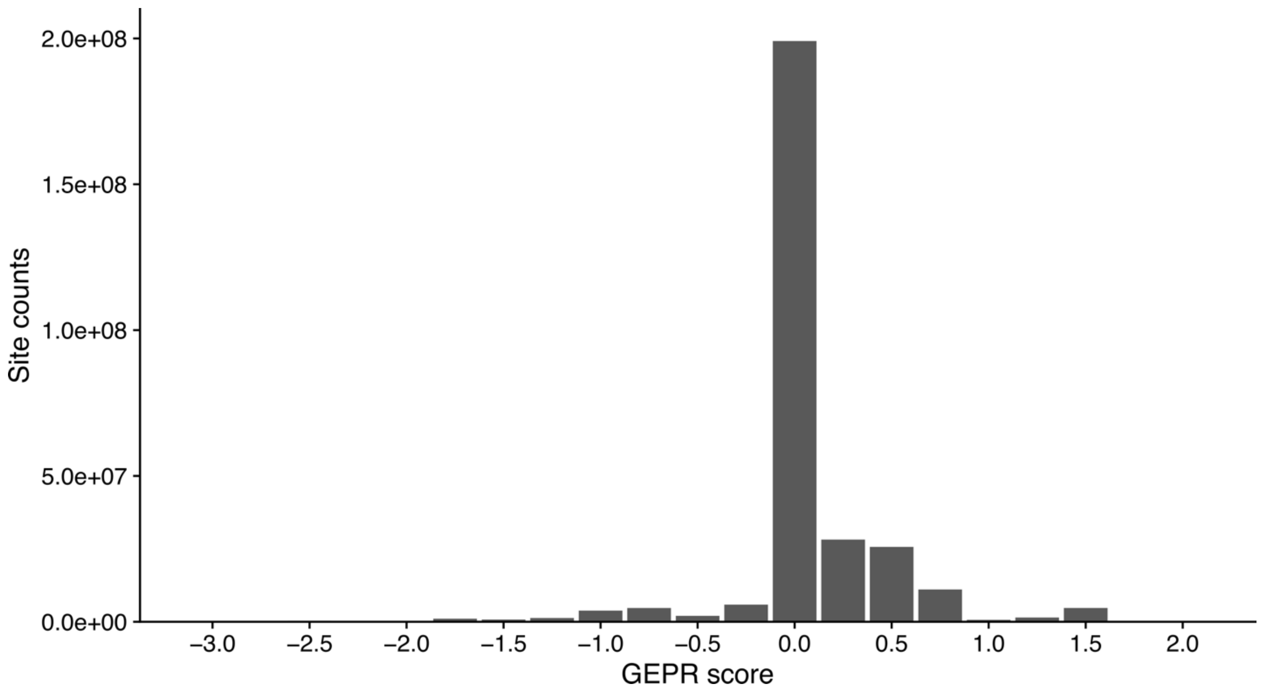


**Fig. S39. Distribution of GERP (Genomic Evolutionary Rate Profiling) scores for apple genome sites.**
The distribution of GERP scores for available sites in the apple genome was obtained from the alignment of seven closely related genome sequences and the thresholds for different types of sites.

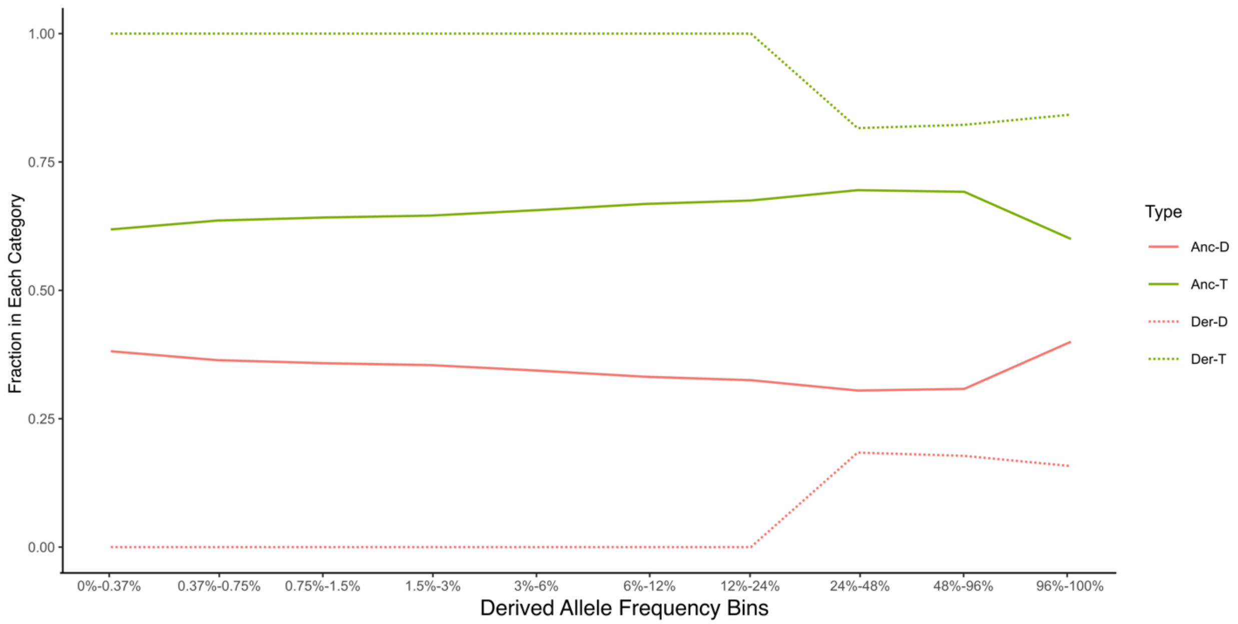


**Fig. S40. The reference bias present in SIFG-4G.**

The x-axis shows the mean population frequency of nonsynonymous SNPs. The y-axis plots the fraction of SNPs in each bin classified as tolerated or deleterious by the SIFT-4G. The y-axis is shown separately according to whether the genome reference sequence carries the ancestral or the derived allele. When the reference carries the ancestral allele, an SNP is classified as deleterious with a probability that ranges from nearly 30% to 40% (solid red line). In contrast, for SNPs where the reference carries the derived allele, the range of fraction of Deleterious alleles is low (dotted red line).

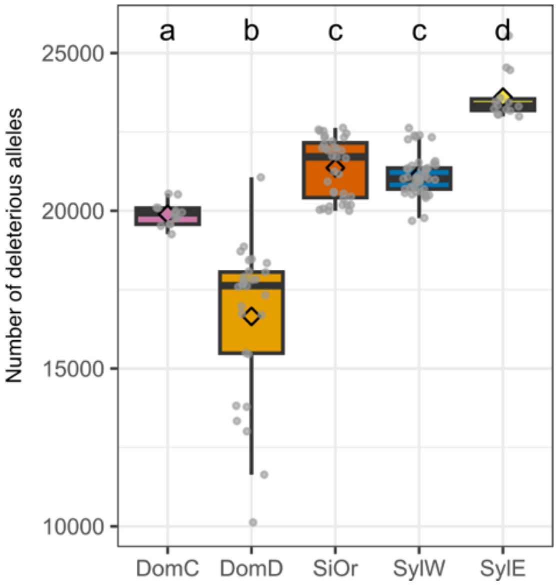


**Fig. S41. Total mutation burden across five apple populations, measured as the number of deleterious alleles individual accessions within each population.**

Different letters represent significant differences (Wilcoxon test, *p*-value < 0.05) between populations. DomC, cider apple individuals; DomD, dessert apple individuals; SiOr, *M. sieversii* and *M. orientalis* individuals; SylE, eastern Europe *M. sylvestris* individuals; SylW, western Europe *M. sylvestris* individuals.

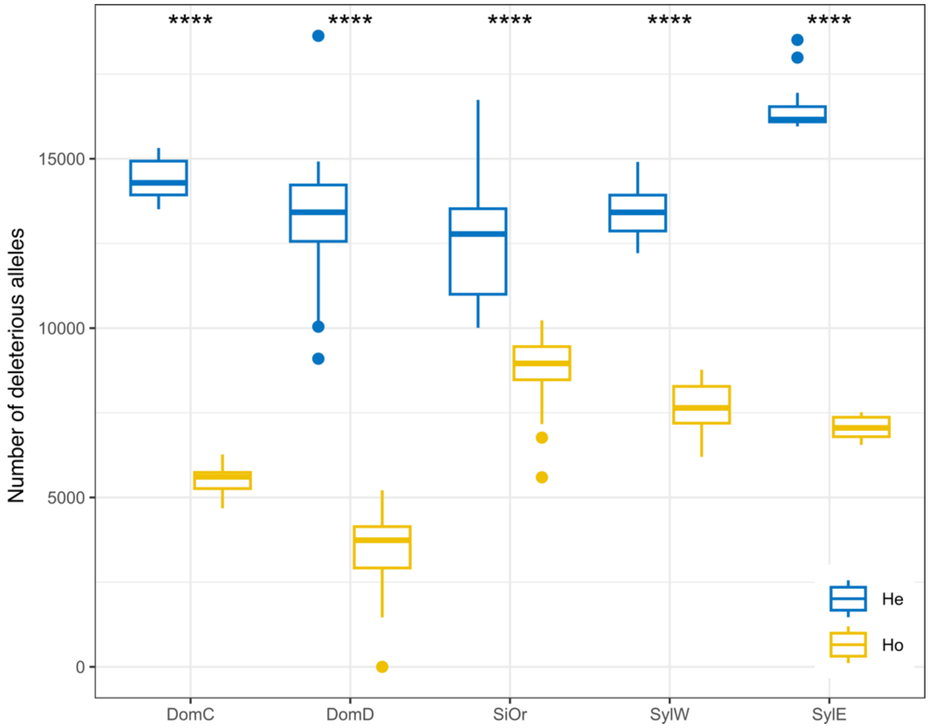


**Fig. S42. Box plot distributions of heterozygote and homozygote mutation burden in cultivar and wild apple populations.**

He, heterozygous site; Ho, homozygous site. DomC, cider apple individuals; DomD, dessert apple individuals; SiOr, *M. sieversii* and *M. orientalis* individuals; SylE, eastern Europe *M. sylvestris* individuals; SylW, western Europe *M. sylvestris* individuals. The star marks significant differences (****: Wilcoxon test *p*-value < 0.0001).

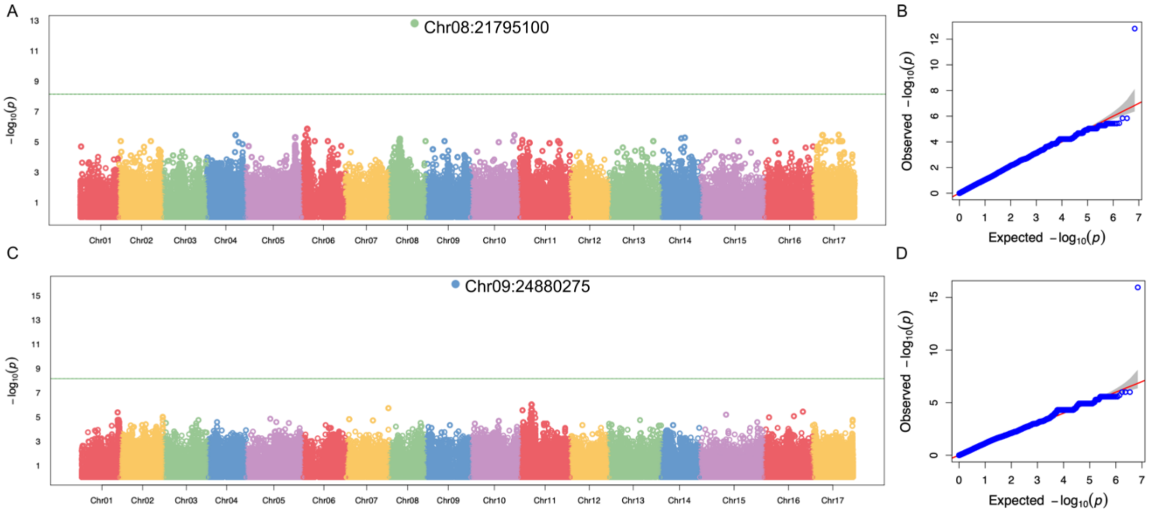


**Fig. S43. Results of genome-wide association study for flowering time.**
(A) Manhattan plot of $–\log_{10}(p)$ values based on the K+Q model. (B) Quantile–quantile (Q-Q) plot comparing the observed and expected $–\log_{10}$(p) values. based on the K+Q model. (C) Manhattan plot of $–\log_{10}(p)$ values based on the K model. (D) QQ plot comparing the observed and expected $–\log_{10}(p)$ values based on the K model. The solid horizontal line indicates the genome-wide significance threshold after Bonferroni correction ($p$ = 7.22 × $10^{-9}$). The dashed horizontal line represents the FDR-adjusted significance threshold at 0.05.

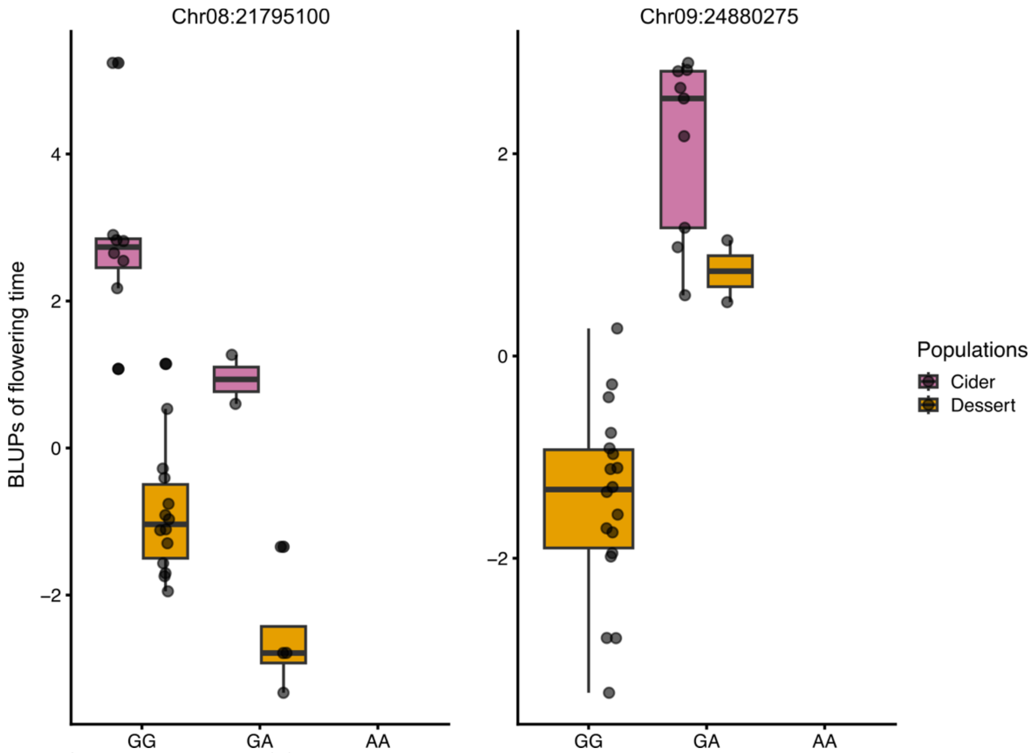


**Fig. S44. Boxplots of flowering time BLUPs across genotypes and cider and dessert apple populations.**
The y-axis shows best linear unbiased predictions (BLUPs) of flowering time, and the x-axis indicates different genotypes. Box colors correspond to different subpopulations.

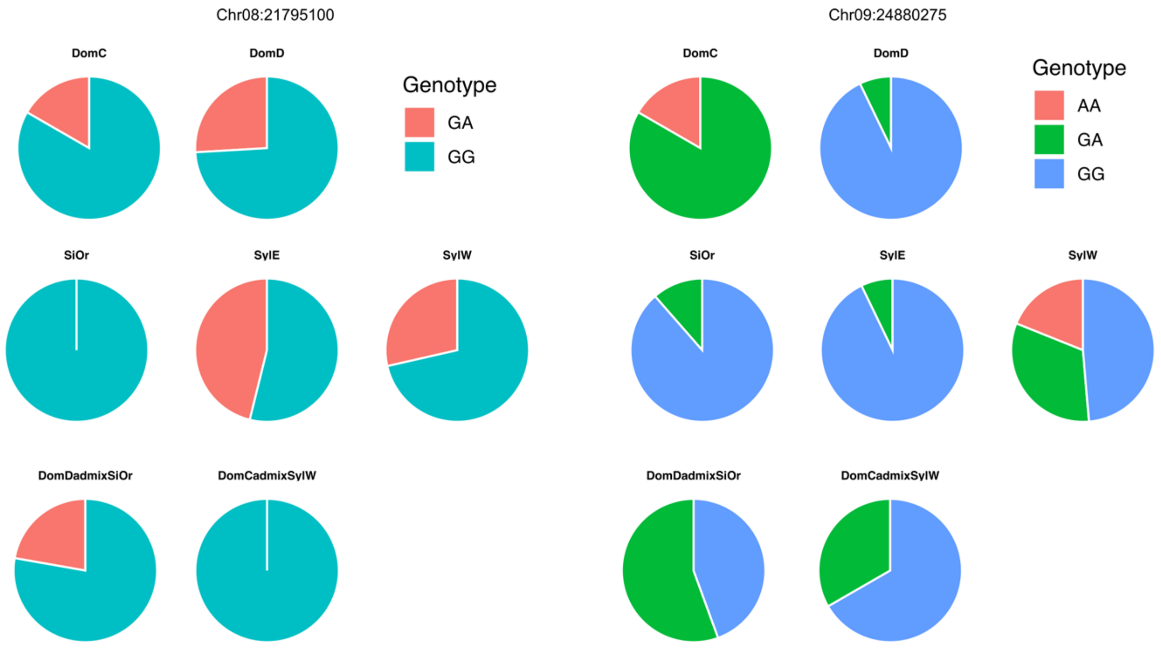


**Fig. S45. Genotype distributions of two SNP loci detected with GWAS across wild and cultivated apple populations.**
Each pie chart represents the proportion of genotypes for a given SNP. Different colors indicate different subpopulations. DomC, Cider *M. domestica*; DomD, Dessert *M. domestica*; SiOr, *M. sieversii* and *M. orientalis*; SylE, Eastern Europe *M. sylvestris*; SylW, Western Europe *M. sylvestris*; Bacc, *M. baccata*; XadmixX, group of individuals classified as admixed between two populations (i.e., individuals assigned with membership coefficient < 0.8 to any given cluster at K=6 with fastSTRUCTURE analyses)

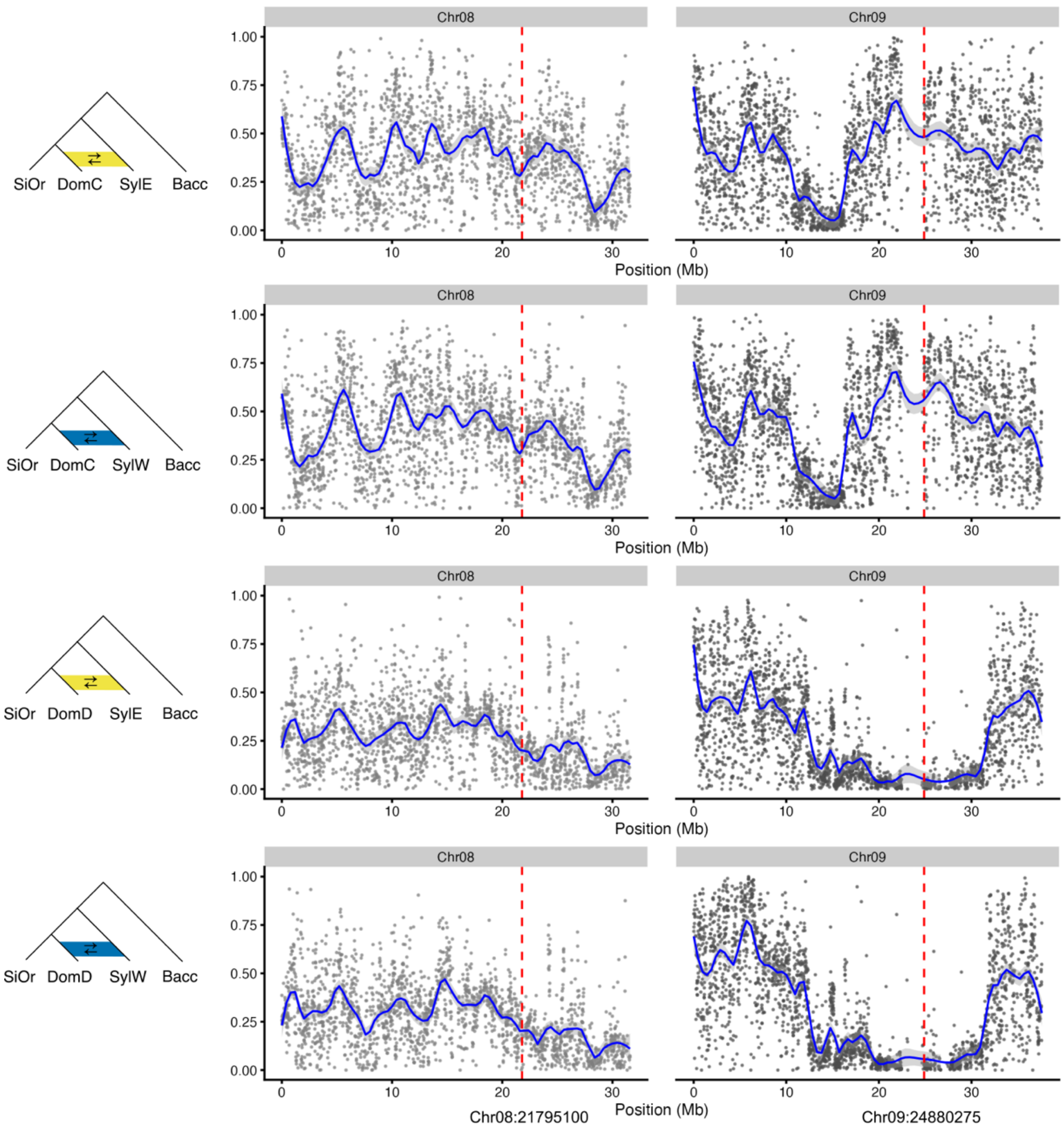


**Fig. S46. $f_d$ values around flowering time-associated GWAS SNPs on chromosomes Chr08 and Chr09.**

Blue lines show a loess fit with 95% confidence intervals. Vertical dashed lines indicate the positions of the GWAS SNPs. $f_d$ analyses used P1 (SiOr), P2 (DomC or DomD), P3 (SylE or SylW), and outgroup (Bacc). DomC, Cider *M. domestica*; DomD, Dessert *M. domestica*; SiOr, *M. sieversii* and *M. orientalis*; SylE, Eastern Europe *M. sylvestris*; SylW, Western Europe *M. sylvestris*; Bacc, *M. baccata*.

**Table S1. Sequencing and mapping statistic of all *Malus* accessions in this study**

| Seq_ID | Raw reads | Raw bases | Clean reads | Clean bases | Effective(%) | Depth(X) | Mapped Reads | Mapped bases | Mapped Depth (X) | Mapped Coverage (%) |
|---|---|---|---|---|---|---|---|---|---|---|
| 858C | 118255938 | 14781992250 | 116814198 | 14585546330 | 98.67% | 23.34 | 109854282 | 13329534935 | 18.78 | 83.45% |
| JFB-siev | 149175598 | 18646949750 | 147291022 | 18390355033 | 98.62% | 29.43 | 140551982 | 17074833384 | 24.06 | 84.82% |
| STO-01 | 145184228 | 18148028500 | 143343254 | 17897348815 | 98.62% | 28.64 | 135705606 | 16467808387 | 23.20 | 83.74% |
| STO-28 | 134989236 | 16873654500 | 133138288 | 16624191036 | 98.52% | 26.61 | 118331861 | 14347933317 | 20.22 | 83.27% |
| X8941 | 153651882 | 19206485250 | 151650182 | 18933236895 | 98.58% | 30.30 | 152470704 | 18486334456 | 26.05 | 84.62% |
| X8942 | 169994834 | 16999483400 | 162369626 | 16204234867 | 95.32% | 25.93 | 162172311 | 15877610943 | 22.37 | 84.36% |
| X8981 | 144428980 | 18053622500 | 142525614 | 17794876030 | 98.57% | 28.48 | 143047664 | 17354149025 | 24.45 | 84.18% |
| X9651 | 178377638 | 17837763800 | 170921146 | 17057662193 | 95.63% | 27.30 | 170641963 | 16717046458 | 23.55 | 83.95% |
| jura-22 | 116962884 | 14620360500 | 115408122 | 14408612703 | 98.55% | 23.06 | 109775007 | 13317293521 | 18.76 | 82.95% |
| A001 | 152905716 | 22935857400 | 152101374 | 22482068683 | 98.02% | 35.98 | 154151830 | 22099162442 | 31.14 | 84.42% |
| A002 | 157922764 | 23688414600 | 157073102 | 23217467468 | 98.01% | 37.16 | 159232059 | 22817647237 | 32.15 | 84.20% |
| A003 | 151637792 | 22745668800 | 150701936 | 22274262285 | 97.93% | 35.65 | 152659329 | 21895359481 | 30.85 | 84.31% |
| A004 | 142941332 | 21441199800 | 142002346 | 20987869338 | 97.89% | 33.59 | 143900789 | 20628149163 | 29.07 | 84.49% |
| A005 | 164813822 | 24722073300 | 163866664 | 24221445943 | 97.97% | 38.76 | 165867575 | 23822715594 | 33.57 | 84.80% |
| A006 | 148112538 | 22216880700 | 147131644 | 21747048086 | 97.89% | 34.80 | 149102507 | 21365372614 | 30.10 | 84.21% |
| A007 | 182831482 | 27424722300 | 181882516 | 26882386896 | 98.02% | 43.02 | 184322281 | 26426849312 | 37.24 | 84.62% |
| A008 | 174418080 | 26162712000 | 173402068 | 25630734145 | 97.97% | 41.02 | 175799940 | 25168046399 | 35.46 | 84.22% |
| A009 | 143018922 | 21452838300 | 142213074 | 21019462541 | 97.98% | 33.64 | 144154480 | 20650544576 | 29.10 | 84.36% |
| A010 | 145361610 | 21804241500 | 144589334 | 21371105801 | 98.01% | 34.20 | 146579382 | 20996708799 | 29.59 | 84.37% |
| A011 | 138874518 | 20831177700 | 138114042 | 20413900625 | 98.00% | 32.67 | 139820354 | 20078262632 | 28.29 | 84.73% |
| A012 | 167466820 | 25120023000 | 166502206 | 24610604766 | 97.97% | 39.39 | 168581514 | 24213401217 | 34.12 | 84.76% |
| A013 | 127469816 | 19120472400 | 126774754 | 18738645183 | 98.00% | 29.99 | 128328205 | 18430066407 | 25.97 | 84.79% |
| A014 | 139403354 | 20910503100 | 138657692 | 20494585857 | 98.01% | 32.80 | 140366512 | 20168819646 | 28.42 | 84.65% |

| | | | | | | | | | | |
|---|---|---|---|---|---|---|---|---|---|---|
| A015 | 125330500 | 18799575000 | 124647620 | 18424202445 | 98.00% | 29.49 | 126201815 | 18123788129 | 25.54 | 84.62% |
| A016 | 168531988 | 25279798200 | 167538642 | 24762553320 | 97.95% | 39.63 | 169471088 | 24364262166 | 34.33 | 85.22% |
| A017 | 142782030 | 21417304500 | 141989264 | 20987390525 | 97.99% | 33.59 | 143711972 | 20647238751 | 29.09 | 84.20% |
| A018 | 137531478 | 20629721700 | 136734402 | 20211034006 | 97.97% | 32.35 | 138401624 | 19888446183 | 28.02 | 84.30% |
| A019 | 155988782 | 23398317300 | 154913278 | 22898081455 | 97.86% | 36.65 | 156213926 | 22595114952 | 31.83 | 85.86% |
| A020 | 147150212 | 22072531800 | 146326428 | 21627996041 | 97.99% | 34.61 | 147214151 | 21396370493 | 30.14 | 86.73% |
| A021 | 167090478 | 25063571700 | 165985100 | 24533698067 | 97.89% | 39.26 | 167992663 | 24137971803 | 34.01 | 85.07% |
| A022 | 150395760 | 22559364000 | 149469680 | 22093172752 | 97.93% | 35.36 | 150743157 | 21807582095 | 30.72 | 86.08% |
| A023 | 159326412 | 23898961800 | 158289252 | 23394624644 | 97.89% | 37.44 | 159449186 | 23088832394 | 32.53 | 85.79% |
| A024 | 155129954 | 23269493100 | 154108490 | 22779333255 | 97.89% | 36.46 | 155564133 | 22463956941 | 31.65 | 85.10% |
| A025 | 193434312 | 29015146800 | 192204940 | 28407237886 | 97.90% | 45.46 | 194456502 | 27963904447 | 39.40 | 84.93% |
| A027 | 172664914 | 25899737100 | 171529172 | 25352368206 | 97.89% | 40.57 | 173431984 | 24955920260 | 35.16 | 85.37% |
| A028 | 150850740 | 22627611000 | 149793972 | 22138346658 | 97.84% | 35.43 | 151981721 | 21731635289 | 30.62 | 83.04% |
| A029 | 154133248 | 23119987200 | 153098472 | 22627893541 | 97.87% | 36.21 | 155380480 | 22206616238 | 31.29 | 83.29% |
| A030 | 161836494 | 24275474100 | 160795640 | 23766818766 | 97.90% | 38.04 | 163141082 | 23328490453 | 32.87 | 83.00% |
| A031 | 164010064 | 24601509600 | 162958292 | 24086545686 | 97.91% | 38.55 | 165273430 | 23649519093 | 33.32 | 84.90% |
| A032 | 141216244 | 21182436600 | 140264398 | 20731528030 | 97.87% | 33.18 | 142395900 | 20349961516 | 28.68 | 82.98% |
| A033 | 153930886 | 23089632900 | 152954402 | 22609129418 | 97.92% | 36.18 | 155301267 | 22185908813 | 31.26 | 83.74% |
| A034 | 132423996 | 19863599400 | 131515150 | 19436919354 | 97.85% | 31.11 | 133530903 | 19071612524 | 26.88 | 82.99% |
| A035 | 153025032 | 22953754800 | 151963646 | 22463077856 | 97.86% | 35.95 | 154008157 | 22054667147 | 31.08 | 83.04% |
| A036 | 137364188 | 20604628200 | 136487616 | 20173054910 | 97.91% | 32.28 | 138605291 | 19797131944 | 27.90 | 83.15% |
| A037 | 145785918 | 21867887700 | 144853984 | 21412158198 | 97.92% | 34.27 | 146732675 | 21037159310 | 29.64 | 83.33% |
| A038 | 125178252 | 18776737800 | 124400002 | 18387043044 | 97.92% | 29.43 | 126269433 | 18044072477 | 25.43 | 83.05% |
| A039 | 181193088 | 27178963200 | 180019148 | 26608331268 | 97.90% | 42.58 | 182542596 | 26109483550 | 36.79 | 83.45% |
| A040 | 136890506 | 20533575900 | 136006572 | 20103494381 | 97.91% | 32.17 | 137973994 | 19727821340 | 27.80 | 82.63% |
| B1 | 177569812 | 26635471800 | 176435348 | 26077942880 | 97.91% | 41.73 | 178373956 | 25545924104 | 36.00 | 84.69% |
| B2 | 156263134 | 23439470100 | 155202682 | 22938455754 | 97.86% | 36.71 | 157260069 | 22552200830 | 31.78 | 85.06% |

| | | | | | | | | | | |
|---|---|---|---|---|---|---|---|---|---|---|
| B3 | 180258960 | 27038844000 | 179034786 | 26461870527 | 97.87% | 42.35 | 181070038 | 26044410348 | 36.70 | 85.12% |
| B4 | 169313776 | 25397066400 | 168196094 | 24859159686 | 97.88% | 39.78 | 169602677 | 24399305117 | 34.38 | 85.10% |
| C01 | 123752202 | 18562830300 | 123007844 | 18178547089 | 97.93% | 29.09 | 124930180 | 17841045758 | 25.14 | 82.75% |
| C02 | 124775016 | 18716252400 | 123971584 | 18322475603 | 97.90% | 29.32 | 125734573 | 17980711874 | 25.34 | 82.73% |
| C03 | 126365524 | 18954828600 | 125625076 | 18568522556 | 97.96% | 29.72 | 127450085 | 18225555311 | 25.68 | 82.55% |
| C04 | 129579290 | 19436893500 | 128781296 | 19034899769 | 97.93% | 30.46 | 130641684 | 18691209982 | 26.34 | 83.18% |
| C05 | 122692684 | 18403902600 | 121977146 | 18028811148 | 97.96% | 28.85 | 123754434 | 17698576664 | 24.94 | 83.08% |
| C06 | 118267772 | 17740165800 | 117386830 | 17350414165 | 97.80% | 27.77 | 119077073 | 17022586978 | 23.99 | 83.26% |
| C07 | 137706264 | 20655939600 | 136853018 | 20227646081 | 97.93% | 32.37 | 138795013 | 19847223170 | 27.97 | 83.90% |
| C08 | 123338022 | 18500703300 | 122586290 | 18118725959 | 97.94% | 29.00 | 124371220 | 17771493075 | 25.04 | 82.76% |
| C09 | 129376128 | 19406419200 | 128487092 | 18990768773 | 97.86% | 30.39 | 130364992 | 18625792318 | 26.25 | 83.19% |
| C10 | 125051202 | 18757680300 | 124389632 | 18385161984 | 98.01% | 29.42 | 126350504 | 18054347446 | 25.44 | 82.96% |
| D01 | 135233278 | 20284991700 | 134404700 | 19864730101 | 97.93% | 31.79 | 136427441 | 19496142066 | 27.47 | 82.98% |
| D02 | 119127864 | 17869179600 | 118446058 | 17506122483 | 97.97% | 28.02 | 120157809 | 17190664148 | 24.22 | 83.30% |
| D03 | 129158800 | 19373820000 | 128260364 | 18956618283 | 97.85% | 30.34 | 130095479 | 18603187538 | 26.21 | 83.61% |
| D04 | 124671108 | 18700666200 | 123936408 | 18318649977 | 97.96% | 29.32 | 125716657 | 17980751173 | 25.34 | 83.49% |
| D05 | 119494930 | 17924239500 | 118816408 | 17561983946 | 97.98% | 28.11 | 120557496 | 17239551929 | 24.29 | 82.44% |
| D06 | 124746484 | 18711972600 | 123982352 | 18325455842 | 97.93% | 29.33 | 125676712 | 17992253793 | 25.35 | 84.11% |
| D07 | 129576042 | 19436406300 | 128754128 | 19029109903 | 97.90% | 30.45 | 130735135 | 18666567442 | 26.30 | 82.96% |
| D08 | 120952744 | 18142911600 | 120211378 | 17767607026 | 97.93% | 28.43 | 122032952 | 17436332572 | 24.57 | 82.66% |
| D09 | 121988662 | 18298299300 | 121229850 | 17916937219 | 97.92% | 28.67 | 123018423 | 17587828905 | 24.78 | 83.03% |
| D10 | 122635082 | 18395262300 | 121898514 | 18016163126 | 97.94% | 28.83 | 123649395 | 17683966191 | 24.92 | 83.40% |
| D11 | 138524606 | 20778690900 | 137666846 | 20348465956 | 97.93% | 32.57 | 139715816 | 19963756984 | 28.13 | 83.02% |
| D12 | 133899098 | 20084864700 | 133054744 | 19664712120 | 97.91% | 31.47 | 135466361 | 19225019445 | 27.09 | 84.17% |
| D13 | 153237986 | 22985697900 | 152348048 | 22517211762 | 97.96% | 36.04 | 154458521 | 22119602591 | 31.17 | 83.31% |
| D14 | 133462876 | 20019431400 | 132638592 | 19602336349 | 97.92% | 31.37 | 134665288 | 19228019428 | 27.10 | 83.40% |
| D15 | 129336242 | 19400436300 | 128562926 | 19001749040 | 97.94% | 30.41 | 130403160 | 18651440103 | 26.28 | 82.97% |

| | | | | | | | | | | |
|---|---|---|---|---|---|---|---|---|---|---|
| D16 | 130910206 | 19636530900 | 130090046 | 19227696916 | 97.92% | 30.77 | 131943174 | 18866902720 | 26.59 | 82.72% |
| D17 | 130312188 | 19546828200 | 129450136 | 19132868034 | 97.88% | 30.62 | 131329305 | 18777691921 | 26.46 | 84.57% |
| D18 | 121022836 | 18153425400 | 120324252 | 17784046102 | 97.97% | 28.46 | 122239233 | 17450211706 | 24.59 | 83.18% |
| D19 | 119705014 | 17955752100 | 119049250 | 17596902630 | 98.00% | 28.16 | 120836393 | 17265768953 | 24.33 | 82.95% |
| D20 | 134357962 | 20153694300 | 133681470 | 19760105035 | 98.05% | 31.62 | 135710551 | 19386801920 | 27.32 | 83.26% |
| E01 | 124325116 | 18648767400 | 123631678 | 18273762744 | 97.99% | 29.24 | 125523284 | 17917565333 | 25.25 | 83.24% |
| E02 | 121801586 | 18270237900 | 121031012 | 17887918628 | 97.91% | 28.63 | 122745848 | 17555916451 | 24.74 | 83.25% |
| E03 | 121840768 | 18276115200 | 121080170 | 17895745844 | 97.92% | 28.64 | 122762942 | 17567106120 | 24.75 | 83.51% |
| E04 | 118492708 | 17773906200 | 117753240 | 17405353699 | 97.93% | 27.86 | 119422492 | 17087457576 | 24.08 | 83.16% |
| E05 | 142632090 | 21394813500 | 141707054 | 20944792091 | 97.90% | 33.52 | 143693044 | 20549786632 | 28.96 | 83.64% |
| E06 | 142638728 | 21395809200 | 141835044 | 20963884196 | 97.98% | 33.55 | 143917142 | 20587817994 | 29.01 | 83.08% |
| E07 | 128758240 | 19313736000 | 128024128 | 18923536644 | 97.98% | 30.28 | 129928484 | 18571332068 | 26.17 | 82.84% |
| E08 | 123631232 | 18544684800 | 122781504 | 18147228808 | 97.86% | 29.04 | 124495979 | 17814657360 | 25.10 | 82.58% |
| E09 | 123793142 | 18568971300 | 123107712 | 18196524917 | 97.99% | 29.12 | 125021194 | 17854924640 | 25.16 | 83.05% |
| E10 | 119807828 | 17971174200 | 119130266 | 17607357784 | 97.98% | 28.18 | 120892646 | 17280169382 | 24.35 | 82.75% |
| E11 | 122821020 | 18423153000 | 122160512 | 18057048009 | 98.01% | 28.90 | 123938795 | 17724870967 | 24.98 | 82.92% |
| E12 | 118462830 | 17769424500 | 117774624 | 17407307027 | 97.96% | 27.86 | 119465843 | 17089229371 | 24.08 | 84.32% |
| E13 | 135176406 | 20276460900 | 134385558 | 19862953673 | 97.96% | 31.79 | 136288436 | 19495464652 | 27.47 | 82.93% |
| E14 | 123814278 | 18572141700 | 122994126 | 18179299703 | 97.88% | 29.09 | 124795847 | 17832008867 | 25.13 | 82.62% |
| E15 | 125170018 | 18775502700 | 124324168 | 18374979424 | 97.87% | 29.41 | 126103280 | 18028664163 | 25.40 | 82.73% |
| E16 | 127217294 | 19082594100 | 126392660 | 18681252931 | 97.90% | 29.90 | 128233105 | 18330321912 | 25.83 | 82.70% |
| E17 | 145247482 | 21787122300 | 144364064 | 21337841942 | 97.94% | 34.15 | 146480943 | 20943343106 | 29.51 | 83.27% |
| E18 | 135569540 | 20335431000 | 134772646 | 19919563988 | 97.95% | 31.88 | 136654994 | 19553588537 | 27.55 | 82.81% |
| E19 | 124161418 | 18624212700 | 123359370 | 18233367613 | 97.90% | 29.18 | 125130329 | 17884424574 | 25.20 | 82.68% |
| E20 | 124385356 | 18657803400 | 123582704 | 18265368943 | 97.90% | 29.23 | 125459467 | 17922925473 | 25.26 | 83.71% |
| F01 | 124645930 | 18696889500 | 123932022 | 18316431214 | 97.97% | 29.31 | 125712382 | 17981922808 | 25.34 | 82.76% |
| F02 | 124783176 | 18717476400 | 123942656 | 18318659625 | 97.87% | 29.32 | 125737499 | 17972672565 | 25.33 | 83.25% |

| | | | | | | | | | | |
|---|---|---|---|---|---|---|---|---|---|---|
| F03 | 122909162 | 18436374300 | 122105266 | 18048164720 | 97.89% | 28.88 | 123866370 | 17709683488 | 24.95 | 82.92% |
| F04 | 120102002 | 18015300300 | 119380864 | 17644306558 | 97.94% | 28.24 | 121232752 | 17308275130 | 24.39 | 83.08% |
| F05 | 124144624 | 18621693600 | 123284890 | 18222917709 | 97.86% | 29.16 | 125047758 | 17877551630 | 25.19 | 83.10% |
| F06 | 119813874 | 17972081100 | 119103840 | 17604377803 | 97.95% | 28.17 | 120754287 | 17293394306 | 24.37 | 84.28% |
| F07 | 133467402 | 20020110300 | 132654498 | 19606725972 | 97.94% | 31.38 | 134644421 | 19232082062 | 27.10 | 82.92% |
| F08 | 126216128 | 18932419200 | 125343524 | 18527136929 | 97.86% | 29.65 | 127147622 | 18180421279 | 25.62 | 82.86% |
| F09 | 121945176 | 18291776400 | 121245032 | 17918728523 | 97.96% | 28.68 | 123016869 | 17587770501 | 24.78 | 83.43% |
| F10 | 124858648 | 18728797200 | 124100956 | 18342009526 | 97.93% | 29.35 | 125839621 | 18011802896 | 25.38 | 84.09% |
| F20 | 126658974 | 18998846100 | 125933378 | 18612677683 | 97.97% | 29.79 | 127782676 | 18264367619 | 25.74 | 82.86% |
| SP01 | 125138514 | 18770777100 | 124426940 | 18390320504 | 97.97% | 29.43 | 125822436 | 18013865884 | 25.38 | 84.30% |
| SP02 | 125529540 | 18829431000 | 124791666 | 18441996135 | 97.94% | 29.51 | 126385522 | 18104109126 | 25.51 | 84.17% |
| SP03 | 125355628 | 18803344200 | 124651258 | 18422526467 | 97.97% | 29.48 | 126263469 | 18070531786 | 25.46 | 84.37% |
| SP04 | 124657858 | 18698678700 | 123915458 | 18314389722 | 97.94% | 29.31 | 125570949 | 17977223221 | 25.33 | 84.40% |
| SP05 | 126735398 | 19010309700 | 125993382 | 18620976370 | 97.95% | 29.80 | 127586348 | 18282271640 | 25.76 | 84.50% |
| SP06 | 122484854 | 18372728100 | 121689228 | 17985134644 | 97.89% | 28.78 | 123409697 | 17666548541 | 24.89 | 84.35% |
| SP07 | 122019860 | 18302979000 | 121282664 | 17924876097 | 97.93% | 28.69 | 122619570 | 17571322828 | 24.76 | 84.40% |
| SP08 | 149432216 | 22414832400 | 148535812 | 21952343470 | 97.94% | 35.13 | 150163596 | 21548132175 | 30.36 | 85.10% |
| SP09 | 120445188 | 18066778200 | 118607662 | 17530394690 | 97.03% | 28.06 | 119937498 | 17114324969 | 24.11 | 84.14% |
| SP10 | 129536590 | 19430488500 | 128398576 | 18975793657 | 97.66% | 30.37 | 129887900 | 18584386552 | 26.19 | 84.91% |
| SP11 | 118180166 | 17727024900 | 116750040 | 17255935202 | 97.34% | 27.62 | 118065324 | 16869030149 | 23.77 | 84.50% |
| SP12 | 118592754 | 17788913100 | 117298474 | 17337246393 | 97.46% | 27.75 | 118617972 | 16930186756 | 23.86 | 84.28% |
| T06 | 122480192 | 18372028800 | 121711012 | 17987570521 | 97.91% | 28.79 | 123733923 | 17646033928 | 24.87 | 82.91% |
| T07 | 125271734 | 18790760100 | 124531270 | 18404554511 | 97.94% | 29.45 | 126613943 | 18054891717 | 25.44 | 82.80% |
| T08 | 143317622 | 21497643300 | 142326102 | 21036701900 | 97.86% | 33.67 | 144585360 | 20629161778 | 29.07 | 83.49% |
| T09 | 146109648 | 21916447200 | 145191226 | 21459793410 | 97.92% | 34.34 | 147438013 | 21050240247 | 29.66 | 83.28% |
| T10 | 135646990 | 20347048500 | 134795476 | 19922642997 | 97.91% | 31.88 | 136705151 | 19560535910 | 27.56 | 84.19% |
| BACC01 | 120063104 | 18009465600 | 119071552 | 17598351865 | 97.72% | 28.16 | 122494725 | 16976188813 | 23.94 | 77.38% |

| | | | | | | | | | | |
|---|---|---|---|---|---|---|---|---|---|---|
| BACC02 | 138651726 | 20797758900 | 137547890 | 20329467975 | 97.75% | 32.53 | 141450943 | 19592490757 | 27.62 | 77.68% |
| BACC03 | 165152022 | 24772803300 | 163798194 | 24210042331 | 97.73% | 38.75 | 168470063 | 23336518103 | 32.90 | 78.19% |
| BACC04 | 167818886 | 25172832900 | 166510716 | 24611221638 | 97.77% | 39.39 | 170982790 | 23686747155 | 33.40 | 78.22% |
| BACC05 | 109344468 | 16401670200 | 108528702 | 16041336791 | 97.80% | 25.67 | 111714998 | 15456793568 | 21.79 | 77.17% |
| BACC06 | 141358472 | 21203770800 | 140164502 | 20716737446 | 97.70% | 33.15 | 140829130 | 19592648195 | 27.62 | 77.92% |
| BACC07 | 121365358 | 18204803700 | 120313500 | 17781288796 | 97.67% | 28.46 | 123588588 | 17120792706 | 24.14 | 77.55% |
| BACC08 | 163988216 | 24598232400 | 162748264 | 24053025551 | 97.78% | 38.49 | 167380720 | 23190182451 | 32.70 | 78.03% |
| BACC09 | 129401124 | 19410168600 | 128536728 | 19001617908 | 97.90% | 30.41 | 130848233 | 18193747008 | 25.65 | 80.20% |
| BACC10 | 132484198 | 19872629700 | 131434744 | 19426066488 | 97.75% | 31.09 | 135224451 | 18736024159 | 26.42 | 77.49% |
| Bac_01 | 124035510 | 12527586510 | 113749970 | 10977043339 | 87.62% | 17.57 | 111850830 | 10390026017 | 14.64 | 82.97% |
| Bac_02 | 73888726 | 11083308900 | 67937306 | 10174772779 | 91.80% | 16.28 | 69541819 | 9852141141 | 13.89 | 74.47% |
| Bac_06 | 71410040 | 10711506000 | 66361912 | 9941038098 | 92.81% | 15.91 | 68097324 | 9606369872 | 13.54 | 74.74% |
| Dom_07 | 154714978 | 19168062178 | 137720558 | 16326370988 | 85.17% | 26.13 | 136159445 | 15821737473 | 22.29 | 84.27% |
| Dom_09 | 169353684 | 21210122894 | 145533474 | 17414820312 | 82.11% | 27.87 | 142810011 | 16819341895 | 23.69 | 85.12% |
| Dom_22 | 174552082 | 17455208200 | 168530470 | 16099222959 | 92.23% | 25.76 | 168026001 | 15813181495 | 22.28 | 85.84% |
| Dom_23 | 160607508 | 16060750800 | 155726526 | 14877014667 | 92.63% | 23.81 | 150517821 | 14082358108 | 19.84 | 84.52% |
| Ori_01 | 167076616 | 20179245616 | 144253876 | 16391962628 | 81.23% | 26.23 | 142813160 | 15931349079 | 22.44 | 84.85% |
| Sie_K_13 | 129297914 | 15928600814 | 110653840 | 12788873638 | 80.29% | 20.47 | 109892121 | 12391008475 | 17.46 | 82.77% |
| Sie_K_15 | 130480520 | 15679594820 | 117034668 | 13341393435 | 85.09% | 21.35 | 115874752 | 12887778605 | 18.16 | 83.88% |
| Sie_X_01 | 162263054 | 14603674860 | 159169758 | 14313725861 | 98.01% | 22.91 | 158124147 | 13974732405 | 19.69 | 83.30% |
| Sie_X_02 | 135829218 | 12224629620 | 133277940 | 11985385055 | 98.04% | 19.18 | 132587971 | 11714357870 | 16.50 | 82.96% |
| Sie_X_03 | 155348698 | 13981382820 | 152804326 | 13741876758 | 98.29% | 21.99 | 152111004 | 13439951720 | 18.94 | 83.21% |
| Sie_X_04 | 174345548 | 15691099320 | 170072336 | 15293393057 | 97.47% | 24.48 | 167636704 | 14796517545 | 20.85 | 83.42% |
| Sie_X_05 | 153499400 | 13814946000 | 150968330 | 13576706979 | 98.28% | 21.73 | 148741621 | 13142250230 | 18.52 | 83.20% |
| Sie_X_07 | 178067162 | 16026044580 | 174535910 | 15697466419 | 97.95% | 25.12 | 173436420 | 15333881798 | 21.60 | 83.31% |
| Sie_X_08 | 181061400 | 16295526000 | 177002700 | 15916199726 | 97.67% | 25.47 | 176094629 | 15549027640 | 21.91 | 83.38% |
| Sie_X_09 | 179670776 | 16170369840 | 175513656 | 15781858973 | 97.60% | 25.26 | 170951364 | 15092788402 | 21.26 | 83.35% |

| | | | | | | | | | | |
|---|---|---|---|---|---|---|---|---|---|---|
| Sie_X_10 | 156207812 | 14058703080 | 153718376 | 13823655928 | 98.33% | 22.12 | 152574626 | 13482269896 | 18.99 | 83.29% |
| Sie_X_11 | 185752776 | 16717749840 | 182313622 | 16395205963 | 98.07% | 26.24 | 179783936 | 15880432883 | 22.37 | 83.42% |
| Sie_X_12 | 150571790 | 13551461100 | 147894528 | 13300300201 | 98.15% | 21.29 | 146852983 | 12975070232 | 18.28 | 83.16% |
| Sie_X_13 | 170185372 | 15316683480 | 163085980 | 14665382565 | 95.75% | 23.47 | 161613118 | 14265832207 | 20.10 | 84.31% |
| Sie_X_14 | 210369524 | 18933257160 | 205412450 | 18473527991 | 97.57% | 29.56 | 204184853 | 18043557604 | 25.42 | 84.04% |
| Sie_X_15 | 154820950 | 13933885500 | 150492632 | 13533197401 | 97.12% | 21.66 | 149265934 | 13179778030 | 18.57 | 84.20% |
| Syl_08 | 156022570 | 19053129070 | 130435130 | 14818738014 | 77.78% | 23.72 | 127809560 | 14154494300 | 19.94 | 84.55% |
| Syl_09 | 110833114 | 11194144514 | 105768730 | 10222458610 | 91.32% | 16.36 | 103617483 | 9806998140 | 13.82 | 84.07% |
| Syl_10 | 150797116 | 19307879816 | 126169756 | 15267610485 | 79.07% | 24.43 | 125581563 | 14611659614 | 20.59 | 83.41% |
| Sun_A01 | 145800000 | 21870000000 | 131652570 | 18662948534 | 85.34% | 29.87 | 132805051 | 18211559762 | 25.66 | 84.08% |
| Sun_A02 | 145800000 | 21870000000 | 139005958 | 19276538186 | 88.14% | 30.85 | 140983113 | 18921438953 | 26.66 | 83.64% |
| Sun_A03 | 145800000 | 21870000000 | 140514980 | 20146417063 | 92.12% | 32.24 | 141686786 | 19754910032 | 27.832 | 84.44% |
| Sun_A04 | 145800000 | 21870000000 | 141823084 | 20431253057 | 93.42% | 32.70 | 143230050 | 20029016274 | 28.22 | 84.08% |
| Sun_A05 | 145800000 | 21870000000 | 142294298 | 20615799214 | 94.27% | 32.99 | 144094581 | 20294071622 | 28.593 | 83.75% |
| Sun_A06 | 145800000 | 21870000000 | 141239032 | 20124040678 | 92.02% | 32.21 | 142921045 | 19777204365 | 27.864 | 84.31% |
| Sun_A08 | 145800000 | 21870000000 | 141362114 | 20302694329 | 92.83% | 32.49 | 142081467 | 19835573303 | 27.946 | 84.44% |
| Sun_A10 | 145800000 | 21870000000 | 138577574 | 18951100593 | 86.65% | 30.33 | 140325738 | 18579969743 | 26.178 | 84.51% |
| Sun_A11 | 145800000 | 21870000000 | 141574758 | 20502365411 | 93.75% | 32.81 | 142512732 | 20083372179 | 28.295 | 84.64% |
| Sun_A12 | 145800000 | 21870000000 | 140993702 | 20115218130 | 91.98% | 32.19 | 142870780 | 19732494937 | 27.803 | 83.30% |
| Sun_A13 | 145800000 | 21870000000 | 137427526 | 19085327926 | 87.27% | 30.54 | 139131730 | 18698482984 | 26.345 | 84.76% |
| Sun_A14 | 145800000 | 21870000000 | 134677630 | 18859031838 | 86.23% | 30.18 | 136341565 | 18534177391 | 26.113 | 84.41% |
| Sun_A15 | 145800000 | 21870000000 | 137056754 | 19460310872 | 88.98% | 31.14 | 139478236 | 18974812841 | 26.739 | 84.19% |
| Sun_A16 | 145800000 | 21870000000 | 135997404 | 19412303771 | 88.76% | 31.07 | 137683549 | 19073386774 | 26.873 | 85.09% |
| Sun_A17 | 145800000 | 21870000000 | 141798200 | 20222039046 | 92.46% | 32.36 | 143188595 | 19850083718 | 27.967 | 84.33% |
| Sun_A18 | 145800000 | 21870000000 | 138427276 | 19847429908 | 90.75% | 31.76 | 140170662 | 19477906880 | 27.443 | 84.49% |
| Sun_A19 | 145800000 | 21870000000 | 142157448 | 20361712174 | 93.10% | 32.59 | 143959282 | 19988197221 | 28.163 | 83.99% |
| Sun_A20 | 145800000 | 21870000000 | 140173806 | 20238863331 | 92.54% | 32.39 | 141989646 | 19925386734 | 28.073 | 84.18% |

| | | | | | | | | | | |
|---|---|---|---|---|---|---|---|---|---|---|
| Sun_A21 | 145800000 | 21870000000 | 139495668 | 19100277911 | 87.34% | 30.57 | 141112446 | 18807035072 | 26.496 | 84.59% |
| Sun_X1 | 145800000 | 21870000000 | 139485772 | 19829926842 | 90.67% | 31.74 | 140566243 | 19518522658 | 27.497 | 85.48% |
| Sun_X2 | 145800000 | 21870000000 | 139531764 | 19449288608 | 88.93% | 31.13 | 140996144 | 19157313913 | 26.989 | 85.11% |
| Sun_A35 | 126921514 | 19038227100 | 124316276 | 17948627368 | 94.28% | 28.72 | 125773435 | 17546175239 | 24.724 | 81.71% |
| Sun_A36 | 145800000 | 21870000000 | 138237838 | 19553875562 | 89.41% | 31.29 | 140111325 | 19088419209 | 26.897 | 82.99% |
| Sun_A37 | 145800000 | 21870000000 | 141021540 | 19932639803 | 91.14% | 31.90 | 142727631 | 19488114976 | 27.46 | 83.94% |
| Sun_A38 | 145800000 | 21870000000 | 141438636 | 20020317653 | 91.54% | 32.04 | 143398177 | 19610320063 | 27.633 | 83.74% |
| Sun_A39 | 145800000 | 21870000000 | 140811722 | 19974771001 | 91.33% | 31.97 | 142939759 | 19564015874 | 27.567 | 84.25% |
| Sun_A40 | 145800000 | 21870000000 | 138859814 | 19760155437 | 90.35% | 31.62 | 140094789 | 19258170962 | 27.135 | 84.20% |
| Sun_A41 | 145800000 | 21870000000 | 139644796 | 19590552496 | 89.58% | 31.35 | 141892918 | 19194671146 | 27.046 | 84.35% |
| Sun_A42 | 145800000 | 21870000000 | 141242380 | 20294279186 | 92.80% | 32.48 | 143366977 | 19922264090 | 28.072 | 83.82% |
| Sun_A43 | 145800000 | 21870000000 | 138042836 | 18191609877 | 83.18% | 29.11 | 140040094 | 17835106312 | 25.13 | 82.34% |
| Sun_A44 | 145800000 | 21870000000 | 139352576 | 18870000212 | 86.28% | 30.20 | 146621187 | 19415076066 | 27.341 | 82.79% |
| Sun_A45 | 145800000 | 21870000000 | 139212262 | 19573775809 | 89.50% | 31.33 | 140467944 | 19062258625 | 26.86 | 83.59% |
| Sun_A46 | 145800000 | 21870000000 | 138482914 | 19179455348 | 87.70% | 30.69 | 140005181 | 18765893369 | 26.441 | 84.45% |
| Sun_A48 | 145800000 | 21870000000 | 135580732 | 17433814748 | 79.72% | 27.90 | 137133061 | 17093003160 | 24.084 | 83.16% |
| Sun_C96 | 154820950 | 13933885500 | 150488970 | 13531629657 | 97.11% | 21.66 | 149261991 | 13178250977 | 18.566 | 84.20% |
| Sun_X3 | 145800000 | 21870000000 | 139908694 | 19865418924 | 90.83% | 31.79 | 141300169 | 19354528943 | 27.272 | 83.84% |
| Sun_X4 | 145800000 | 21870000000 | 140069272 | 19685932070 | 90.01% | 31.50 | 142110719 | 19263472331 | 27.145 | 82.93% |
| Sun_X5 | 145800000 | 21870000000 | 141184106 | 20351878351 | 93.06% | 32.57 | 143667078 | 19825475992 | 27.94 | 83.64% |
| Sun_X6 | 145800000 | 21870000000 | 140201600 | 19967576921 | 91.30% | 31.96 | 142118616 | 19531037659 | 27.521 | 83.34% |
| Sun_X7 | 145800000 | 21870000000 | 142928024 | 20671552704 | 94.52% | 33.08 | 144496196 | 20198391302 | 28.461 | 82.60% |
| Sun_X8 | 145800000 | 21870000000 | 138504990 | 19750615012 | 90.31% | 31.61 | 140629781 | 19376497997 | 27.303 | 83.02% |
| Sun_X9 | 145800000 | 21870000000 | 140206912 | 18925342878 | 86.54% | 30.29 | 142160845 | 18568759197 | 26.164 | 82.11% |
| Sun_A51 | 145800000 | 21870000000 | 129512774 | 18220045883 | 83.31% | 29.16 | 131128180 | 17844561396 | 25.143 | 82.56% |
| Sun_A52 | 145800000 | 21870000000 | 139241144 | 19756361416 | 90.34% | 31.62 | 140538805 | 19265604418 | 27.145 | 83.10% |
| Sun_A53 | 145800000 | 21870000000 | 142520710 | 20497924922 | 93.73% | 32.80 | 144682851 | 20108542557 | 28.334 | 82.76% |

| | | | | | | | | | | |
|---|---|---|---|---|---|---|---|---|---|---|
| Sun_A54 | 145800000 | 21870000000 | 142461294 | 20597404669 | 94.18% | 32.96 | 144325355 | 20172000486 | 28.423 | 82.81% |
| Sun_A55 | 145800000 | 21870000000 | 143023036 | 20603699243 | 94.21% | 32.97 | 144825150 | 20116290204 | 28.345 | 82.85% |
| Sun_X10 | 145800000 | 21870000000 | 141072248 | 20198181263 | 92.36% | 32.32 | 143209119 | 19829556805 | 27.94 | 82.68% |
| Sun_X11 | 145800000 | 21870000000 | 141954676 | 20185363968 | 92.30% | 32.30 | 145257353 | 19805224182 | 27.908 | 83.16% |
| Sun_X12 | 145800000 | 21870000000 | 137931350 | 19886866217 | 90.93% | 31.83 | 139730176 | 19483187373 | 27.453 | 82.60% |
| Sun_X13 | 145800000 | 21870000000 | 137561544 | 18701598582 | 85.51% | 29.93 | 139927473 | 18330422420 | 25.828 | 83.14% |

Note: We take 145.8 million reads per sample from (33), which corresponds to approximately 30-fold coverage.

**Table S2. Summary of identified SNPs and filter conditions.**

<table>
<tr><th>Data set</th><th>Filter name</th><th>Filter condition</th><th>individual nubmer</th><th>variant number</th><th>Note</th></tr>
<tr><td>Raw genotypes</td><td>Level 1 filters</td><td>gatk  VariantFiltration \<br>-filter-expression "(vc.isSNP() &&<br>(vc.hasAttribute('ReadPosRankSum') && ReadPosRankSum<br>< -8.0)) || ((vc.isIndel() || vc.isMixed()) &&<br>(vc.hasAttribute('ReadPosRankSum') && ReadPosRankSum<br>< -20.0)) || (vc.hasAttribute('QD') && QD < 2.0) " \<br>--filter-name "badSeq" \<br>--filter-expression "(vc.isSNP() && ((vc.hasAttribute('FS')<br>&& FS > 60.0) || (vc.hasAttribute('SOR') &&  SOR > 3.0))) ||<br>((vc.isIndel() || vc.isMixed()) && ((vc.hasAttribute('FS') &&<br>FS > 200.0) || (vc.hasAttribute('SOR') &&  SOR > 10.0)))" \<br>--filter-name "badStrand" \<br>--filter-expression "vc.isSNP() && ((vc.hasAttribute('MQ')<br>&& MQ < 40.0) || (vc.hasAttribute('MQRankSum') &&<br>MQRankSum < -12.5))" \<br>--filter-name "badMap"</td><td>218</td><td>485,815,973</td><td>SNPs, Indel<br>and<br>Invariant sites</td></tr>
<tr><td>Filtered variant data</td><td>Level 2 filters</td><td>bcftools filter -S . -e 'FMT/DP<5 | FMT/GQ<20 |<br>FMT/DP>100' \<br>| bcftools filter --SnpGap 10 \<br>| bcftools view -m2 -M2 -v snps \ # biallelic SNPs<br>| bcftools filter -e 'F_MISSING > 0.2'<br>remove clone individaul<br>remove individuals with high missing level (>30%)</td><td>201</td><td>28,377,551</td><td>Biallelic SNPs</td></tr>
<tr><td>Invariant sites</td><td>Level 2' filters</td><td>bcftools filter -S . -e 'FMT/DP<5 | FMT/GQ<20 |<br>FMT/DP>100' \<br>| bcftools filter -e 'F_MISSING > 0.2'<br>remove clone individaul<br>remove individuals with high missing level (>30%)</td><td>201</td><td>303,978,197</td><td>Invariant sites</td></tr>
<tr><td>Unlinked variant data</td><td>Level 3 filters</td><td>bcftools filter -e 'MAF <= 0.05'<br>keep synonymous sites based on SnpEff annoatation<br>information.<br>plink --indep-pairwise 5 kb 1 0.2<br>vcftools  --thin 5000</td><td>201</td><td>31,300</td><td></td></tr>
<tr><td>Final variant data</td><td>-</td><td>remove admixed individuals</td><td>144</td><td>26,767,333</td><td></td></tr>
</table>

**Table S3. Clustering of ancestry components for *Malus* accessions inferred with fastSTRUCTURE at *K* = 6.**

| Seq_ID | Q1(SylW) | Q2(SiOr) | Q3(Bacc) | Q4(DomD) | Q5(SylE) | Q6(DomC) |
|---|---|---|---|---|---|---|
| 858C | 0.9503 | 0.0000 | 0.0000 | 0.0138 | 0.0000 | 0.0359 |
| A001 | 0.0000 | 0.0083 | 0.0000 | 0.1363 | 0.0000 | 0.8554 |
| A002 | 0.0000 | 0.0000 | 0.0000 | 0.0000 | 0.0000 | 1.0000 |
| A003 | 0.1934 | 0.0936 | 0.0000 | 0.6352 | 0.0007 | 0.0772 |
| A004 | 0.0000 | 0.0049 | 0.0000 | 0.0128 | 0.0000 | 0.9822 |
| A005 | 0.0000 | 0.0000 | 0.0000 | 0.3515 | 0.0000 | 0.6485 |
| A006 | 0.0000 | 0.0000 | 0.0000 | 0.0758 | 0.0000 | 0.9242 |
| A007 | 0.0000 | 0.0000 | 0.0000 | 0.0000 | 0.0000 | 1.0000 |
| A008 | 0.0000 | 0.0000 | 0.0000 | 0.0000 | 0.0000 | 1.0000 |
| A009 | 0.0000 | 0.0032 | 0.0000 | 0.0827 | 0.0000 | 0.9141 |
| A010 | 0.0623 | 0.3136 | 0.0000 | 0.3538 | 0.0603 | 0.2100 |
| A011 | 0.0000 | 0.0000 | 0.0000 | 0.8768 | 0.0000 | 0.1232 |
| A012 | 0.0000 | 0.0000 | 0.0000 | 0.8600 | 0.0000 | 0.1400 |
| A013 | 0.0000 | 0.0006 | 0.0000 | 0.6926 | 0.0000 | 0.3068 |
| A014 | 0.0640 | 0.0722 | 0.0000 | 0.7237 | 0.0000 | 0.1400 |
| A015 | 0.0000 | 0.0059 | 0.0000 | 0.9469 | 0.0000 | 0.0472 |
| A016 | 0.0693 | 0.0000 | 0.0000 | 0.8649 | 0.0000 | 0.0658 |
| A017 | 0.0000 | 0.0063 | 0.0000 | 0.9936 | 0.0000 | 0.0000 |
| A018 | 0.0000 | 0.0001 | 0.0000 | 0.9999 | 0.0000 | 0.0000 |
| A019 | 0.0000 | 0.0000 | 0.0000 | 1.0000 | 0.0000 | 0.0000 |
| A020 | 0.0000 | 0.0000 | 0.0000 | 1.0000 | 0.0000 | 0.0000 |
| A021 | 0.0440 | 0.0092 | 0.0000 | 0.8235 | 0.0000 | 0.1233 |
| A022 | 0.0000 | 0.0140 | 0.0000 | 0.9859 | 0.0000 | 0.0000 |
| A023 | 0.0000 | 0.0000 | 0.0000 | 1.0000 | 0.0000 | 0.0000 |
| A024 | 0.0000 | 0.0000 | 0.0000 | 1.0000 | 0.0000 | 0.0000 |
| A025 | 0.0000 | 0.0000 | 0.0000 | 1.0000 | 0.0000 | 0.0000 |
| A027 | 0.0000 | 0.0729 | 0.0056 | 0.9215 | 0.0000 | 0.0000 |
| A028 | 0.0000 | 1.0000 | 0.0000 | 0.0000 | 0.0000 | 0.0000 |

| | | | | | | |
|---|---|---|---|---|---|---|
| A029 | 0.0000 | 1.0000 | 0.0000 | 0.0000 | 0.0000 | 0.0000 |
| A030 | 0.0000 | 1.0000 | 0.0000 | 0.0000 | 0.0000 | 0.0000 |
| A031 | 0.0109 | 0.1972 | 0.0000 | 0.2749 | 0.4663 | 0.0506 |
| A032 | 0.0000 | 0.9930 | 0.0000 | 0.0052 | 0.0000 | 0.0017 |
| A033 | 0.0000 | 0.0000 | 0.0000 | 0.0000 | 1.0000 | 0.0000 |
| A034 | 0.0000 | 1.0000 | 0.0000 | 0.0000 | 0.0000 | 0.0000 |
| A035 | 1.0000 | 0.0000 | 0.0000 | 0.0000 | 0.0000 | 0.0000 |
| A036 | 0.0000 | 1.0000 | 0.0000 | 0.0000 | 0.0000 | 0.0000 |
| A037 | 0.9566 | 0.0033 | 0.0000 | 0.0400 | 0.0000 | 0.0000 |
| A038 | 0.9976 | 0.0000 | 0.0000 | 0.0024 | 0.0000 | 0.0000 |
| A039 | 1.0000 | 0.0000 | 0.0000 | 0.0000 | 0.0000 | 0.0000 |
| A040 | 1.0000 | 0.0000 | 0.0000 | 0.0000 | 0.0000 | 0.0000 |
| B1 | 0.0248 | 0.0027 | 0.0000 | 0.0001 | 0.0000 | 0.9724 |
| B2 | 0.0737 | 0.0344 | 0.0000 | 0.8836 | 0.0000 | 0.0083 |
| B3 | 0.0000 | 0.0000 | 0.0000 | 1.0000 | 0.0000 | 0.0000 |
| B4 | 0.0000 | 0.0000 | 0.0000 | 1.0000 | 0.0000 | 0.0000 |
| BACC01 | 0.0000 | 0.0000 | 1.0000 | 0.0000 | 0.0000 | 0.0000 |
| BACC02 | 0.0000 | 0.0000 | 1.0000 | 0.0000 | 0.0000 | 0.0000 |
| BACC03 | 0.0000 | 0.0000 | 1.0000 | 0.0000 | 0.0000 | 0.0000 |
| BACC05 | 0.0000 | 0.0000 | 1.0000 | 0.0000 | 0.0000 | 0.0000 |
| BACC06 | 0.0000 | 0.0000 | 1.0000 | 0.0000 | 0.0000 | 0.0000 |
| BACC07 | 0.0000 | 0.0000 | 1.0000 | 0.0000 | 0.0000 | 0.0000 |
| BACC08 | 0.0000 | 0.0000 | 1.0000 | 0.0000 | 0.0000 | 0.0000 |
| BACC09 | 0.0000 | 0.0000 | 1.0000 | 0.0000 | 0.0000 | 0.0000 |
| BACC10 | 0.0000 | 0.0000 | 1.0000 | 0.0000 | 0.0000 | 0.0000 |
| C01 | 0.0000 | 1.0000 | 0.0000 | 0.0000 | 0.0000 | 0.0000 |
| C02 | 1.0000 | 0.0000 | 0.0000 | 0.0000 | 0.0000 | 0.0000 |
| C03 | 1.0000 | 0.0000 | 0.0000 | 0.0000 | 0.0000 | 0.0000 |
| C04 | 0.9722 | 0.0000 | 0.0000 | 0.0260 | 0.0000 | 0.0018 |
| C05 | 0.9321 | 0.0210 | 0.0000 | 0.0453 | 0.0000 | 0.0015 |

| | | | | | | |
|---|---|---|---|---|---|---|
| C06 | 0.9625 | 0.0000 | 0.0000 | 0.0308 | 0.0000 | 0.0067 |
| C07 | 0.7047 | 0.0334 | 0.0000 | 0.1931 | 0.0660 | 0.0028 |
| C08 | 0.9571 | 0.0000 | 0.0000 | 0.0000 | 0.0429 | 0.0000 |
| C09 | 0.4492 | 0.0000 | 0.0000 | 0.0000 | 0.5508 | 0.0000 |
| C10 | 0.0000 | 0.0000 | 0.0000 | 0.0000 | 1.0000 | 0.0000 |
| D01 | 0.4532 | 0.0000 | 0.0000 | 0.0000 | 0.5468 | 0.0000 |
| D02 | 0.8778 | 0.0015 | 0.0000 | 0.0869 | 0.0000 | 0.0337 |
| D03 | 0.8099 | 0.0000 | 0.0000 | 0.1878 | 0.0000 | 0.0023 |
| D04 | 0.8752 | 0.0000 | 0.0000 | 0.1247 | 0.0000 | 0.0000 |
| D05 | 1.0000 | 0.0000 | 0.0000 | 0.0000 | 0.0000 | 0.0000 |
| D06 | 0.6530 | 0.0003 | 0.0000 | 0.3124 | 0.0000 | 0.0343 |
| D07 | 0.4275 | 0.0000 | 0.0000 | 0.0000 | 0.5725 | 0.0000 |
| D08 | 1.0000 | 0.0000 | 0.0000 | 0.0000 | 0.0000 | 0.0000 |
| D09 | 0.4280 | 0.0000 | 0.0000 | 0.0000 | 0.5720 | 0.0000 |
| D10 | 0.8627 | 0.0198 | 0.0000 | 0.1175 | 0.0000 | 0.0000 |
| D11 | 0.9343 | 0.0000 | 0.0000 | 0.0565 | 0.0000 | 0.0091 |
| D12 | 0.5556 | 0.1094 | 0.1650 | 0.0121 | 0.1543 | 0.0035 |
| D13 | 0.9460 | 0.0000 | 0.0000 | 0.0540 | 0.0000 | 0.0000 |
| D14 | 0.0006 | 0.0000 | 0.0000 | 0.0000 | 0.9994 | 0.0000 |
| D15 | 0.9630 | 0.0000 | 0.0000 | 0.0331 | 0.0000 | 0.0039 |
| D16 | 1.0000 | 0.0000 | 0.0000 | 0.0000 | 0.0000 | 0.0000 |
| D17 | 0.0555 | 0.2229 | 0.0000 | 0.1898 | 0.4991 | 0.0327 |
| D18 | 0.0000 | 0.0000 | 0.0000 | 0.0000 | 1.0000 | 0.0000 |
| D19 | 0.4178 | 0.0000 | 0.0000 | 0.0000 | 0.5822 | 0.0000 |
| D20 | 0.0018 | 0.0008 | 0.0000 | 0.0001 | 0.9749 | 0.0222 |
| Dom_07 | 0.0000 | 0.0000 | 0.0000 | 1.0000 | 0.0000 | 0.0000 |
| Dom_23 | 0.1250 | 0.2511 | 0.0000 | 0.4182 | 0.0119 | 0.1938 |
| E01 | 0.0003 | 0.0000 | 0.0000 | 0.0000 | 0.9997 | 0.0000 |
| E02 | 0.4777 | 0.0000 | 0.0000 | 0.0579 | 0.3936 | 0.0708 |
| E03 | 0.8648 | 0.0000 | 0.0000 | 0.0987 | 0.0000 | 0.0365 |

| | | | | | | |
|---|---|---|---|---|---|---|
| E04 | 0.4548 | 0.0000 | 0.0000 | 0.0000 | 0.5452 | 0.0000 |
| E05 | 0.8677 | 0.0046 | 0.0000 | 0.1257 | 0.0019 | 0.0000 |
| E06 | 0.4158 | 0.0000 | 0.0000 | 0.0000 | 0.5842 | 0.0000 |
| E07 | 0.4291 | 0.0000 | 0.0000 | 0.0000 | 0.5709 | 0.0000 |
| E08 | 1.0000 | 0.0000 | 0.0000 | 0.0000 | 0.0000 | 0.0000 |
| E09 | 0.0000 | 0.0000 | 0.0000 | 0.0000 | 1.0000 | 0.0000 |
| E10 | 1.0000 | 0.0000 | 0.0000 | 0.0000 | 0.0000 | 0.0000 |
| E11 | 0.9825 | 0.0000 | 0.0000 | 0.0040 | 0.0135 | 0.0000 |
| E12 | 0.2446 | 0.1696 | 0.0000 | 0.2185 | 0.2175 | 0.1498 |
| E13 | 1.0000 | 0.0000 | 0.0000 | 0.0000 | 0.0000 | 0.0000 |
| E14 | 1.0000 | 0.0000 | 0.0000 | 0.0000 | 0.0000 | 0.0000 |
| E15 | 1.0000 | 0.0000 | 0.0000 | 0.0000 | 0.0000 | 0.0000 |
| E16 | 1.0000 | 0.0000 | 0.0000 | 0.0000 | 0.0000 | 0.0000 |
| E17 | 0.3983 | 0.0000 | 0.0000 | 0.0000 | 0.6017 | 0.0000 |
| E18 | 1.0000 | 0.0000 | 0.0000 | 0.0000 | 0.0000 | 0.0000 |
| E19 | 0.9999 | 0.0000 | 0.0000 | 0.0000 | 0.0001 | 0.0000 |
| E20 | 0.0000 | 0.0559 | 0.0000 | 0.0023 | 0.9418 | 0.0000 |
| F01 | 1.0000 | 0.0000 | 0.0000 | 0.0000 | 0.0000 | 0.0000 |
| F02 | 0.3941 | 0.0000 | 0.0000 | 0.0000 | 0.6059 | 0.0000 |
| F03 | 1.0000 | 0.0000 | 0.0000 | 0.0000 | 0.0000 | 0.0000 |
| F04 | 0.0000 | 0.0000 | 0.0000 | 0.0000 | 1.0000 | 0.0000 |
| F05 | 0.9285 | 0.0000 | 0.0000 | 0.0541 | 0.0000 | 0.0174 |
| F06 | 0.0000 | 0.0000 | 0.0000 | 0.3111 | 0.6889 | 0.0000 |
| F07 | 0.3973 | 0.0000 | 0.0000 | 0.0000 | 0.6027 | 0.0000 |
| F08 | 0.4257 | 0.0000 | 0.0000 | 0.0000 | 0.5743 | 0.0000 |
| F09 | 0.9273 | 0.0000 | 0.0000 | 0.0727 | 0.0000 | 0.0000 |
| F10 | 0.7181 | 0.0352 | 0.0000 | 0.1443 | 0.0452 | 0.0572 |
| F20 | 0.5263 | 0.0000 | 0.0000 | 0.0178 | 0.4343 | 0.0215 |
| JFB-siev | 0.0953 | 0.1479 | 0.0000 | 0.6991 | 0.0001 | 0.0578 |
| SP01 | 0.0000 | 0.0008 | 0.0000 | 0.0109 | 0.0000 | 0.9882 |

| | | | | | | |
|---|---|---|---|---|---|---|
| SP02 | 0.0000 | 0.0000 | 0.0000 | 0.2831 | 0.0000 | 0.7169 |
| SP03 | 0.0048 | 0.0060 | 0.0000 | 0.0073 | 0.0000 | 0.9819 |
| SP04 | 0.2245 | 0.0605 | 0.0000 | 0.6498 | 0.0000 | 0.0651 |
| SP05 | 0.2188 | 0.1341 | 0.0000 | 0.5586 | 0.0000 | 0.0886 |
| SP06 | 0.0000 | 0.1059 | 0.0000 | 0.0016 | 0.0000 | 0.8925 |
| SP07 | 0.0000 | 0.0000 | 0.0000 | 0.3552 | 0.0000 | 0.6448 |
| SP08 | 0.0000 | 0.0000 | 0.0000 | 0.6014 | 0.0000 | 0.3986 |
| SP09 | 0.0000 | 0.0000 | 0.0000 | 0.0000 | 0.0000 | 1.0000 |
| SP10 | 0.0000 | 0.0000 | 0.0000 | 0.5943 | 0.0000 | 0.4057 |
| SP11 | 0.0000 | 0.0000 | 0.0000 | 0.3730 | 0.0000 | 0.6270 |
| SP12 | 0.0000 | 0.0000 | 0.0000 | 0.0000 | 0.0000 | 1.0000 |
| STO-01 | 0.4057 | 0.0000 | 0.0000 | 0.0000 | 0.5943 | 0.0000 |
| STO-28 | 0.4637 | 0.0000 | 0.0000 | 0.0000 | 0.5363 | 0.0000 |
| Sie_X_01 | 0.0000 | 1.0000 | 0.0000 | 0.0000 | 0.0000 | 0.0000 |
| Sie_X_02 | 0.0000 | 1.0000 | 0.0000 | 0.0000 | 0.0000 | 0.0000 |
| Sie_X_03 | 0.0000 | 1.0000 | 0.0000 | 0.0000 | 0.0000 | 0.0000 |
| Sie_X_04 | 0.0000 | 1.0000 | 0.0000 | 0.0000 | 0.0000 | 0.0000 |
| Sie_X_05 | 0.0000 | 1.0000 | 0.0000 | 0.0000 | 0.0000 | 0.0000 |
| Sie_X_07 | 0.0000 | 1.0000 | 0.0000 | 0.0000 | 0.0000 | 0.0000 |
| Sie_X_08 | 0.0000 | 1.0000 | 0.0000 | 0.0000 | 0.0000 | 0.0000 |
| Sie_X_09 | 0.0000 | 1.0000 | 0.0000 | 0.0000 | 0.0000 | 0.0000 |
| Sie_X_10 | 0.0000 | 1.0000 | 0.0000 | 0.0000 | 0.0000 | 0.0000 |
| Sie_X_11 | 0.0000 | 1.0000 | 0.0000 | 0.0000 | 0.0000 | 0.0000 |
| Sie_X_12 | 0.0000 | 1.0000 | 0.0000 | 0.0000 | 0.0000 | 0.0000 |
| Sie_X_13 | 0.0000 | 1.0000 | 0.0000 | 0.0000 | 0.0000 | 0.0000 |
| Sie_X_14 | 0.0000 | 1.0000 | 0.0000 | 0.0000 | 0.0000 | 0.0000 |
| Sun_A01 | 0.0131 | 0.2657 | 0.0000 | 0.6604 | 0.0000 | 0.0608 |
| Sun_A02 | 0.1288 | 0.2912 | 0.0000 | 0.4838 | 0.0394 | 0.0568 |
| Sun_A03 | 0.0000 | 0.0171 | 0.0000 | 0.9829 | 0.0000 | 0.0000 |
| Sun_A04 | 0.1579 | 0.0981 | 0.0000 | 0.5106 | 0.0000 | 0.2333 |

| | | | | | | |
|---|---|---|---|---|---|---|
| Sun_A06 | 0.0000 | 0.0000 | 0.0000 | 1.0000 | 0.0000 | 0.0000 |
| Sun_A11 | 0.0000 | 0.0000 | 0.0000 | 1.0000 | 0.0000 | 0.0000 |
| Sun_A12 | 0.0000 | 0.6364 | 0.0000 | 0.2729 | 0.0443 | 0.0464 |
| Sun_A13 | 0.0457 | 0.1510 | 0.0173 | 0.7571 | 0.0289 | 0.0000 |
| Sun_A14 | 0.0000 | 0.0161 | 0.0000 | 0.9838 | 0.0000 | 0.0000 |
| Sun_A15 | 0.0017 | 0.1453 | 0.3532 | 0.4785 | 0.0037 | 0.0175 |
| Sun_A16 | 0.0000 | 0.0824 | 0.0000 | 0.8536 | 0.0000 | 0.0640 |
| Sun_A17 | 0.0106 | 0.0386 | 0.0000 | 0.9107 | 0.0000 | 0.0402 |
| Sun_A18 | 0.0628 | 0.1756 | 0.0000 | 0.7605 | 0.0000 | 0.0010 |
| Sun_A19 | 0.1438 | 0.2003 | 0.0000 | 0.6378 | 0.0181 | 0.0000 |
| Sun_A20 | 0.0000 | 0.0000 | 0.0000 | 1.0000 | 0.0000 | 0.0000 |
| Sun_A21 | 0.1099 | 0.0000 | 0.0057 | 0.8626 | 0.0217 | 0.0001 |
| Sun_A35 | 0.0000 | 1.0000 | 0.0000 | 0.0000 | 0.0000 | 0.0000 |
| Sun_A36 | 0.0000 | 1.0000 | 0.0000 | 0.0000 | 0.0000 | 0.0000 |
| Sun_A37 | 0.0117 | 0.6727 | 0.0000 | 0.2909 | 0.0247 | 0.0000 |
| Sun_A38 | 0.0074 | 0.6872 | 0.0000 | 0.2272 | 0.0024 | 0.0759 |
| Sun_A39 | 0.0866 | 0.6371 | 0.0000 | 0.2762 | 0.0000 | 0.0000 |
| Sun_A40 | 0.0661 | 0.5371 | 0.0000 | 0.3954 | 0.0014 | 0.0000 |
| Sun_A41 | 0.0656 | 0.7019 | 0.0000 | 0.2323 | 0.0002 | 0.0000 |
| Sun_A42 | 0.0067 | 0.6963 | 0.0000 | 0.2892 | 0.0074 | 0.0003 |
| Sun_A43 | 0.0000 | 1.0000 | 0.0000 | 0.0000 | 0.0000 | 0.0000 |
| Sun_A44 | 0.0000 | 1.0000 | 0.0000 | 0.0000 | 0.0000 | 0.0000 |
| Sun_A45 | 0.0293 | 0.8184 | 0.0000 | 0.1503 | 0.0019 | 0.0000 |
| Sun_A46 | 0.0426 | 0.4976 | 0.0000 | 0.4598 | 0.0000 | 0.0000 |
| Sun_A48 | 0.0531 | 0.5862 | 0.0000 | 0.3436 | 0.0000 | 0.0171 |
| Sun_A51 | 0.9466 | 0.0000 | 0.0000 | 0.0013 | 0.0521 | 0.0000 |
| Sun_A52 | 0.9175 | 0.0000 | 0.0000 | 0.0377 | 0.0449 | 0.0000 |
| Sun_A53 | 0.9987 | 0.0000 | 0.0000 | 0.0000 | 0.0013 | 0.0000 |
| Sun_A54 | 0.9717 | 0.0000 | 0.0000 | 0.0074 | 0.0207 | 0.0003 |
| Sun_A55 | 0.0000 | 0.0164 | 0.0000 | 0.0899 | 0.8936 | 0.0000 |

| | | | | | | |
|---|---|---|---|---|---|---|
| Sun_X1 | 0.0000 | 0.0000 | 0.0000 | 1.0000 | 0.0000 | 0.0000 |
| Sun_X10 | 0.9975 | 0.0000 | 0.0000 | 0.0000 | 0.0025 | 0.0000 |
| Sun_X11 | 0.0161 | 0.0690 | 0.0000 | 0.0000 | 0.9149 | 0.0000 |
| Sun_X12 | 0.9844 | 0.0000 | 0.0000 | 0.0106 | 0.0036 | 0.0013 |
| Sun_X13 | 0.0051 | 0.0744 | 0.0000 | 0.0362 | 0.8843 | 0.0000 |
| Sun_X2 | 0.0557 | 0.0000 | 0.0000 | 0.9443 | 0.0000 | 0.0000 |
| Sun_X3 | 0.0000 | 0.8167 | 0.0000 | 0.1699 | 0.0000 | 0.0134 |
| Sun_X4 | 0.0000 | 1.0000 | 0.0000 | 0.0000 | 0.0000 | 0.0000 |
| Sun_X5 | 0.0000 | 0.7386 | 0.2401 | 0.0213 | 0.0000 | 0.0000 |
| Sun_X6 | 0.0018 | 0.9335 | 0.0000 | 0.0487 | 0.0021 | 0.0139 |
| Sun_X7 | 0.0000 | 1.0000 | 0.0000 | 0.0000 | 0.0000 | 0.0000 |
| Sun_X8 | 0.0000 | 0.9008 | 0.0000 | 0.0918 | 0.0000 | 0.0074 |
| Sun_X9 | 0.0000 | 1.0000 | 0.0000 | 0.0000 | 0.0000 | 0.0000 |
| T06 | 0.0000 | 1.0000 | 0.0000 | 0.0000 | 0.0000 | 0.0000 |
| T07 | 0.0000 | 1.0000 | 0.0000 | 0.0000 | 0.0000 | 0.0000 |
| T08 | 0.0000 | 0.0000 | 0.0000 | 0.0000 | 1.0000 | 0.0000 |
| T09 | 0.0000 | 0.0000 | 0.0000 | 0.0000 | 1.0000 | 0.0000 |
| T10 | 0.6556 | 0.0869 | 0.0000 | 0.1041 | 0.1361 | 0.0173 |
| X8941 | 0.0580 | 0.7137 | 0.0000 | 0.2235 | 0.0049 | 0.0000 |
| X8942 | 0.0440 | 0.8333 | 0.0000 | 0.1189 | 0.0038 | 0.0000 |
| X8981 | 0.0003 | 0.8125 | 0.0000 | 0.1872 | 0.0000 | 0.0000 |
| X9651 | 0.9609 | 0.0024 | 0.0000 | 0.0367 | 0.0000 | 0.0000 |
| jura-22 | 0.9435 | 0.0000 | 0.0000 | 0.0000 | 0.0565 | 0.0000 |

Note: DomC, Cider *M. domestica*; DomD, Dessert *M. domestica*; SiOr, *M. sieversii* and *M. orientalis*; SylE, Eastern Europe *M. sylvestris*; SylW, Western Europe *M. sylvestris*; Bacc, *M. baccata*.

**Table S4. Mean proportions of different apple species from different geographical sites assigned to six genetic clusters inferred with fastStructure at *K* = 6.**

| Group | Sample size | Q1(SylW) | Q2(SiOr) | Q3(Bacc) | Q4(DomD) | Q5(SylE) | Q6(DomC) |
|---|---|---|---|---|---|---|---|
| ARM_Kp | 9 | 0.00% | 99.92% | 0.00% | 0.06% | 0.00% | 0.02% |
| AUT_NPDA | 6 | 42.15% | 0.00% | 0.00% | 0.00% | 57.85% | 0.00% |
| AUT_STO | 8 | 42.89% | 0.00% | 0.00% | 0.00% | 57.11% | 0.00% |
| CHN_Liaoning | 2 | 0.00% | 0.00% | 100.00% | 0.00% | 0.00% | 0.00% |
| CHN_Shanxi | 7 | 0.00% | 0.00% | 100.00% | 0.00% | 0.00% | 0.00% |
| CHN_Xinjiang | 13 | 0.00% | 100.00% | 0.00% | 0.00% | 0.00% | 0.00% |
| DEU_Bern | 6 | 96.94% | 0.00% | 0.00% | 0.95% | 2.09% | 0.03% |
| DNK_Aaby | 3 | 95.42% | 0.66% | 0.00% | 3.92% | 0.00% | 0.00% |
| DNK_Hvol | 4 | 100.00% | 0.00% | 0.00% | 0.00% | 0.00% | 0.00% |
| DNK_Norh | 3 | 100.00% | 0.00% | 0.00% | 0.00% | 0.00% | 0.00% |
| DNK_Skov | 4 | 100.00% | 0.00% | 0.00% | 0.00% | 0.00% | 0.00% |
| **DOM_CID** | **23** | **3.58%** | **2.25%** | **0.00%** | **22.78%** | **0.00%** | **71.38%** |
| **DOM_DES** | **40** | **2.76%** | **7.37%** | **0.95%** | **84.08%** | **0.57%** | **4.26%** |
| FRA_DeSe | 6 | 89.99% | 0.03% | 0.00% | 9.32% | 0.00% | 0.67% |
| FRA_Ise | 2 | 60.56% | 9.82% | 8.25% | 5.81% | 14.52% | 1.04% |
| FRA_Jur | 6 | 88.43% | 1.14% | 0.00% | 5.69% | 3.74% | 1.00% |
| FRA_Lor | 3 | 97.81% | 0.00% | 0.00% | 1.88% | 0.00% | 0.30% |
| FRA_PdD | 2 | 80.78% | 0.02% | 0.00% | 17.16% | 0.00% | 2.05% |
| FRA_Senart | 3 | 88.70% | 0.15% | 0.00% | 9.28% | 0.06% | 1.80% |
| FRA_WE | 1 | 100.00% | 0.00% | 0.00% | 0.00% | 0.00% | 0.00% |
| FRA_Yv | 7 | 96.71% | 0.38% | 0.00% | 2.35% | 0.00% | 0.56% |
| ITA_AlDom | 3 | 41.62% | 5.65% | 0.00% | 9.81% | 34.85% | 8.07% |
| KAZ_Site1 | 8 | 3.95% | 65.60% | 0.00% | 29.02% | 0.48% | 0.95% |
| KAZ_Site2 | 15 | 1.05% | 88.90% | 1.60% | 8.03% | 0.07% | 0.35% |
| MKD_Site1 | 3 | 0.71% | 5.33% | 0.00% | 4.20% | 89.76% | 0.00% |
| ROU_Site0 | 3 | 0.36% | 6.57% | 0.00% | 19.53% | 71.84% | 1.69% |
| ROU_Site1 | 3 | 0.06% | 0.03% | 0.00% | 0.00% | 99.16% | 0.74% |

| | | | | | | | |
|---|---|---|---|---|---|---|---|
| ROU_Site13 | 1 | 0.00% | 0.00% | 0.00% | 0.00% | 100.00% | 0.00% |
| ROU_Site14 | 1 | 0.00% | 5.59% | 0.00% | 0.23% | 94.18% | 0.00% |
| ROU_Site15 | 2 | 2.78% | 11.15% | 0.00% | 9.49% | 74.96% | 1.64% |
| ROU_Site16 | 1 | 0.00% | 0.00% | 0.00% | 0.00% | 100.00% | 0.00% |
| ROU_Site2 | 3 | 0.03% | 0.00% | 0.00% | 0.00% | 99.97% | 0.00% |

Note: DOM_CID, cider cultivar apple; DOM_DES, dessert cultivar apple.
In addition to dessert and cider cultivar apples, the name of the group is a combination of the country code and the name of the site (Dataset S1).

**Table S5. Summary of population genetic diversity statistics for each apple population.**

| Population ID | $N$ | $A_p$ | $S$ | $H_o$ | $H_e$ | $F_{IS}$ | $\pi$±SD |
|---|---|---|---|---|---|---|---|
| Bacc | 9 | 2,952 | 8216 | 0.09533 | 0.0813 | -0.01563 | 0.0105±0.0072 |
| DomC | 13 | 0 | 20699 | 0.26001 | 0.22116 | -0.06501 | 0.0075±0.0044 |
| DomD | 28 | 20 | 23974 | 0.26091 | 0.22759 | -0.06929 | 0.0074±0.0042 |
| SiOr | 35 | 360 | 19787 | 0.15945 | 0.14816 | -0.00779 | 0.0057±0.0039 |
| SylE | 14 | 35 | 22092 | 0.21801 | 0.20096 | -0.01796 | 0.0077±0.0044 |
| SylW | 45 | 245 | 24369 | 0.19683 | 0.18829 | -0.00335 | 0.0062±0.0040 |

Notes: $N$ is the number of individuals analyzed. $A_p$ is the number of private alleles per population, $S$ is polymorphic sites $H_o$ is the observed heterozygosity, $H_e$ is the expected heterozygosity under Hardy–Weinberg equilibrium, and $F_{IS}$ is the inbreeding coefficient within populations. The nucleotide diversity ($\pi$) based on 10-kb non-overlapping window. DomC, Cider *M. domestica*; DomD, Dessert *M. domestica*; SiOr, *M. sieversii* and *M. orientalis*; SylE, Eastern Europe *M. sylvestris*; SylW, Western Europe *M. sylvestris*; Bacc, *M. baccata*. The summary statistics are based on 31,300 non-synonymous unlinked SNPs.

**Table S6. Patterson's *D*-statistic (ABBA-BABA statistic) estimated with Dsuite among six *Malus* populations using 31,000 synonymous unlinked SNPs.**

| P1 | P2 | P3 | Dstatistic | Z-score | *p*-value | $f_4$-ratio | BBAA | ABBA | BABA |
|---|---|---|---|---|---|---|---|---|---|
| SiOr | DomC | DomD | 0.1098 | 3.4324 | 0.0003 | 0.4232 | 1444 | 1806 | 1449 |
| DomC | DomD | SylE | 0.0072 | 0.3284 | 0.3713 | 0.0201 | 2410 | 1479 | 1458 |
| DomC | DomD | SylW | 0.0239 | 0.8923 | 0.1861 | 0.0447 | 2260 | 1588 | 1514 |
| SiOr | DomC | SylE | 0.3545 | 15.0053 | 0.0000 | 0.4912 | 2546 | 1956 | 932 |
| SiOr | DomC | SylW | 0.4174 | 16.2926 | 0.0000 | 0.4315 | 2523 | 2139 | 879 |
| SylE | SylW | DomC | 0.0797 | 7.3870 | 0.0000 | 0.1524 | 2225 | 1397 | 1191 |
| SiOr | DomD | SylE | 0.3544 | 14.1882 | 0.0000 | 0.5011 | 2570 | 1997 | 952 |
| SiOr | DomD | SylW | 0.4286 | 15.1797 | 0.0000 | 0.4570 | 2537 | 2223 | 889 |
| SylE | SylW | DomD | 0.0986 | 8.1637 | 0.0000 | 0.1826 | 2197 | 1444 | 1185 |
| SylW | SylE | SiOr | 0.0135 | 1.3597 | 0.0870 | 0.0101 | 3165 | 1107 | 1077 |

Note: P1, P2, P3: three populations used in the test. ABBA, BABA and BBAA: count of the ABBA, BABA and BBAA sites. Z-score and *p*-value associated with the *D*-statistics are computed over 20 jack-knife blocks divided the dataset. DomC, Cider *M. domestica*; DomD, Dessert *M. domestica*; SiOr, *M. sieversii* and *M. orientalis*; SylE, Eastern Europe *M. sylvestris*; SylW, Western Europe *M. sylvestris.* Bacc (*M. baccata*) population are used as the outgroup population.

**Table S7. Summary of the statistically non-rejected admixture graphs allowing for up to four migration events in *qpGraph*.**

| Number of migration events | Score of the best supported admixture graph given the number of migration events | Number of statistically non-rejected admixture graphs ($p$>0.05) | Whether reject graphs with lower number of migrations |
|---|---|---|---|
| 0 | 1296.92 | 0 | / |
| 1 | 124.50 | 0 | Yes |
| 2 | 90.75 | 6 | Yes |
| 3 | 76.76 | 22 | Yes |
| 4 | 78.26 | 76 | No |

**Table S8. Demographic parameters in the best scenarios groups (G3D1 and G3D2) ran with fastsimcoal2 to infer the apple domestication history.**

| Demographic parameter | Scenario G3D1 | | | Scenario G3D2 | | |
|---|---|---|---|---|---|---|
| | Media n | 95% interval low | 95% interval high | Media n | 95% interval low | 95% interval high |
| Effective population size (*Ne*) of Bacc | 17901 | 18552.24 | 20897.28 | 19013 | 19214.35 | 21568.97 |
| Effective population size (*Ne*) of DomC | 1353.5 | 1356.76 | 1539.4 | 1251 | 1345.06 | 1706.82 |
| Effective population size (*Ne*) of DomD | 2140.5 | 2090.11 | 2625.81 | 2077 | 1983.48 | 2625.72 |
| Effective population size (*Ne*) of SiOr | 3356 | 2771.09 | 4786.63 | 3331 | 3402.73 | 6116.11 |
| Effective population size (*Ne*) of SylE | 1793.5 | 1765.65 | 2306.87 | 1982 | 1975.62 | 2632.14 |
| Effective population size (*Ne*) of SylW | 2098.5 | 10377.26 | 24506.46 | 2182 | 12024.13 | 26144.43 |
| Divergence time of SylW from Ance (thousand years ago) | 799.76 | 417.48 | 610.04 | 812.91 | 448.08 | 642.60 |
| Divergence time of SiOr from Ance (thousand years ago) | 158.18 | 329.51 | 528.94 | 152.67 | 311.79 | 504.53 |
| Divergence time of SylE from SylW (thousand years ago) | 18.15 | 19.32 | 27.06 | 19.72 | 16.97 | 23.44 |
| Divergence time of DomD from SiOr (thousand years ago) | 11.10 | 10.21 | 14.33 | 9.73 | 9.74 | 13.24 |
| Divergence time of DomC from SiOr (thousand years ago) | 4.75 | 5.34 | 7.88 | NA | NA | NA |
| Divergence time of DomC from DomD (thousand years ago) | NA | NA | NA | 2.40 | 5.26 | 8.83 |
| Migration rates from DomC to SylE per generation ($\times 10^{-3}$) | 0.4 | 0.34 | 0.43 | 0.43 | 0.36 | 0.48 |
| Migration rates from DomC to SylW per generation ($\times 10^{-3}$) | 0.42 | 0.46 | 1.02 | 0.45 | 0.48 | 1.04 |
| Migration rates from DomD to SylE per generation ($\times 10^{-3}$) | 0.37 | 0.35 | 0.47 | 0.22 | 0.24 | 0.36 |
| Migration rates from DomD to SylWper generation ($\times 10^{-3}$) | 0.41 | 0.44 | 1.03 | 0.32 | 0.52 | 1.37 |
| Migration rates from SiOr to SylE per generation ($\times 10^{-3}$) | 3.97 | 5.11 | 10.12 | 6.39 | 7.64 | 16.22 |
| Migration rates from SiOr to SylW per generation ($\times 10^{-3}$) | 4.79 | 6.98 | 18.27 | 7.7 | 8.77 | 18.58 |
| Migration rates from SylE to DomC per generation ($\times 10^{-3}$) | 0.56 | 0.52 | 0.67 | 0.34 | 0.36 | 0.49 |
| Migration rates from SylE to DomD per generation ($\times 10^{-3}$) | 0.3 | 0.33 | 0.46 | 0.26 | 0.34 | 0.51 |
| Migration rates from SylE to SiOr per generation ($\times 10^{-3}$) | 2.29 | 2.83 | 10.02 | 1.53 | 4.45 | 12.43 |

| | | | | | | |
|---|---|---|---|---|---|---|
| Migration rates from SylW to DomC per generation ($\times 10^{-3}$) | 0.43 | 0.39 | 0.51 | 0.32 | 0.31 | 0.41 |
| Migration rates from SylW to DomD per generation ($\times 10^{-3}$) | 0.4 | 0.42 | 0.62 | 0.31 | 0.37 | 0.61 |
| Migration rates from SylW to SiOr per generation ($\times 10^{-3}$) | 1.96 | 4.77 | 15.14 | 2.04 | 3.93 | 10.54 |

Note: Demographic parameters estimated for the best-supported scenario groups G3D1 and G3G2 (see Methods section), as determined by model selection criteria. Parameter estimates include effective population sizes, migration rates, and divergence times. DomC, Cider *M. domestica*; DomD, Dessert *M. domestica*; SiOr, *M. sieversii* and *M. orientalis*; SylE, Eastern Europe *M. sylvestris*; SylW, Western Europe *M. sylvestris.* Bacc (*M. baccata*) population are used as the outgroup population.

**Table S9. Linkage disequilibrium (LD) decay distances for different populations.**

| Population ID | LD decay distance (bp) |
|---|---|
| Bacc | 22,200 |
| DomC | 123,500 |
| DomD | 27,100 |
| SiOr | 6,600 |
| SylE | 15,900 |
| SylW | 7,100 |

Note: LD decay distance represents the physical distance at which the pairwise $r^2$ between SNPs decreases to half of its maximum value. DomC, Cider *M. domestica*; DomD, Dessert *M. domestica*; SiOr, *M. sieversii* and *M. orientalis*; SylE, Eastern Europe *M. sylvestris*; SylW, Western Europe *M. sylvestris.* Bacc (*M. baccata*) population are used as the outgroup population.

**Dataset S1 (separate file).** Sample information and geographic origin of the *Malus* accessions used in this study.

**Dataset S2 (separate file).** Genes associated with positive selection in cultivar and wild apple population.

**Dataset S3 (separate file).** Hard and soft sweep genes KEGG enrichment analysis in cultivar and wild apple populations.

**Dataset S4 (separate file).** Genes associated with balancing selection in cultivar and wild apple populations.

**Dataset S5 (separate file).** Summary of DomC (cider *M. demostica*) and DomD (dessert *M. demostica*) adaptive introgression regions.

**Dataset S6 (separate file).** Genotypes at GWAS-associated loci (Chr08:21795100 and Chr09:24880275) for *Malus* accessions.
